# Transcriptional and functional consequences of alterations to *MEF2C* and its topological organization in neuronal models

**DOI:** 10.1101/2022.02.15.480098

**Authors:** Kiana Mohajeri, Rachita Yadav, Eva D’haene, Philip M. Boone, Serkan Erdin, Dadi Gao, Mariana Moyses-Oliveira, Riya Bhavsar, Benjamin Currall, Kathryn O’Keefe, Chelsea Lowther, Diane Lucente, Nicholas D. Burt, Monica Salani, Matthew Larson, Björn Menten, Derek J.C. Tai, James F. Gusella, Sarah Vergult, Michael E. Talkowski

**Author notes:** Correspondence: Michael E. Talkowski. These authors contributed equally to the work (Kiana Mohajeri, Rachita Yadav).

## Abstract

Point mutations and structural variants directly disrupting the coding sequence of *MEF2C* have been associated with a spectrum of neurodevelopmental disorders (NDDs), while recent studies have also implicated altered noncoding regulation of *MEF2C* expression in NDDs. However, the impact of haploinsufficiency of *MEF2C* on neurodevelopmental pathways and synaptic processes is not well understood, nor are the complex mechanisms that govern regulation of *MEF2C*. To explore the transcriptional and functional changes associated with coding and noncoding structural variants, we generated an allelic series of 204 isogenic iPSC-derived neuronal cell lines harboring CRISPR-engineered mutations that directly delete predominant isoforms of *MEF2C*, as well as deletions to the boundaries of topologically associating domains (TADs) and chromatin loops encompassing *MEF2C*. We then performed systematic profiling of mutation-specific alterations to transcriptional signatures, regulatory interactions, chromatin contacts, and electrophysiological effects. Our analyses reveal that direct deletion of *MEF2C* causes differential expression of genes enriched for neurodevelopmental and synaptic-associated pathways, accompanied by a significant reduction in synaptic firing and synchrony in neurons. By contrast, we observe robust buffering against *MEF2C* regulatory disruption upon deletion of a distal 5q14.3 TAD and loop boundary; however, homozygous loss of proximal loop boundary resulted in significant down-regulation of *MEF2C* expression and significantly reduced electrophysiological activity that was comparable to direct *MEF2C* disruption. Collectively, our findings demonstrate the functional impact of *MEF2C* haploinsufficiency in human-derived neural models and highlight the complex interactions of gene regulation and chromatin topology that challenge a priori regulatory predictions of structural variant disruption to three-dimensional genome organization.

## INTRODUCTION

Over the last decade, genetic studies have established haploinsufficiency of *MEF2C* as a cause of neurodevelopmental disorders (NDDs)^1–12^. Through microarray analysis, exome and genome sequencing, NDDs have been associated with loss-of-function (LoF) mutations in *MEF2C*, including protein-truncating variants (PTVs), structural variants (SVs) including deletions and balanced chromosomal abnormalities (BCAs), and broader microdeletion of the 5q14.3 locus. Molecular studies of *MEF2C* function have further demonstrated sensitivity to dosage of this synaptic regulator in mouse models of conditionally modulated Mef2c expression in neural tissue, which display fundamentally altered brain development and neuronal activity^13–17^. The extensive complementary data from human patients and mouse models has implicated *MEF2C* LoF mutations as a driver of aberrant neurodevelopment with varied consequences that include developmental delay, intellectual disability, autism spectrum disorder, hypotonia, and epilepsy^3,4,10,18–22^. To date, the mechanisms associated with haploinsufficiency of *MEF2C* that contribute to these phenotypic presentations have not been explored in human neuronal models, which can provide insights into signatures of *MEF2C*-specific changes as well as evidence of transcriptional or functional convergence across NDDs.

In contrast to the abundant molecular studies of direct gene disruption in NDDs, functional interpretation of noncoding variation remains a considerable challenge^23,24^, though multiple studies have described highly penetrant noncoding mutations across rare NDDs and Mendelian disorders^25–27^. One emerging mutational mechanism not captured by exome and genome sequencing is regulatory changes associated with three-dimensional (3D) genome organization^28–30^. Early glimpses into the intricacies of this architecture demonstrated the partitioning of chromatin into topologically associating domains (TADs), and the smaller loops within them. It is presently understood that TADs demarcate neighborhoods of long-range regulatory interaction, while loops facilitate punctate enhancer and promoter connections^31,32^. There are now examples in the literature of pathogenic consequences of positional effects through the disruption of TADs and loops^31,33–35^, as well as studies that have demonstrated an uncoupling of topological rewiring and gene expression^29,32,36,37^. These studies collectively underscore the complexity of long-range regulatory mechanisms and the significant challenges associated with prediction of functional consequences associated with alterations to TAD boundaries and 3D regulatory organization. These studies suggest that the contributions of functional elements are likely to be context-specific and require functional modeling to dissect these diverse regulatory mechanisms of individual loci.

We previously demonstrated through whole-genome sequencing of NDD cases that chromosome 5q14.3 harbored an unusual and genome-wide significant excess of noncoding BCA breakpoints that did not directly disrupt *MEF2C* but that all occurred within the TAD boundaries encompassing *MEF2C*^38^. This distribution of breakpoints in proximity to *MEF2C* was further supported by microdeletions in NDD cases reported in DECIPHER that apparently did not directly alter the gene locus (at the available resolution of chromosomal microarray)^39^. In considering the landscape of de novo SVs across the 5q14.3 locus in NDD cases, the unifying thread appears to be recurrent distal boundary disruption. Taken together, these data suggest that both direct disruption of MEF2C and alterations to its 3D regulatory architecture may result in comparable molecular mechanisms in NDD cases. Motivated by these findings, we have performed a systematic molecular dissection of the 5q14.3 locus to quantify the transcriptomic and electrophysiological effects of *MEF2C* LoF in human neural derivatives. Through the generation of an allelic series of CRISPR-engineered human induced pluripotent stem cell (hiPSC)-derived neural stem cells (NSCs) and glutamatergic neurons (iNs), we interrogated the impact of enhancer, TAD boundary, and loop boundary deletion on local genome organization, local expression effects on *MEF2C*, and global transcriptional signatures. Our analyses reveal that direct *MEF2C* alteration results in both transcriptional and functional changes to the synapse. Moreover, we find that disruption of the distal boundary of the *MEF2C*-containing loop is insufficient to produce indirect *MEF2C* haploinsufficiency, whereas disruption of the proximal boundary of the same 3D structure results in haploinsufficiency of *MEF2C* that is comparable to direct gene disruption. Overall, these data suggest that the effects of direct and indirect *MEF2C* disruption contribute to cell type-specific alterations on neuronal functions that converge on synaptic deficits in neurodevelopment.

## RESULTS

### Haploinsufficiency of MEF2C is associated with altered expression of highly constrained genes in NSCs and synaptic genes in iNs

There is strong evidence for association between microdeletions and LoF point mutations that disrupt *MEF2C* and a spectrum of NDDs. We therefore first sought to determine the transcriptional changes in early neuronal development stem cells and fully differentiated neurons caused by direct disruption of *MEF2C*. We generated targeted heterozygous (DEL^het^) and homozygous (DEL^hom^) deletions of MEF2C in hiPSCs using dual-guide CRISPR/Cas9 genome editing. Following single-cell isolation and screening, we retained both edited clones and, as matched controls, clones that were exposed to all experimental conditions but were not edited. Six replicates per genotype then underwent differentiation to NSCs and iNs for transcriptional profiling using RNAseq **(Figure 1A)**. *MEF2C* expression effects were confirmed to be commensurate with zygosity via protein expression analyses **(SupplementaryMethodsSection1;SupplementaryFigure1-4)**.

**Figure. 1.**
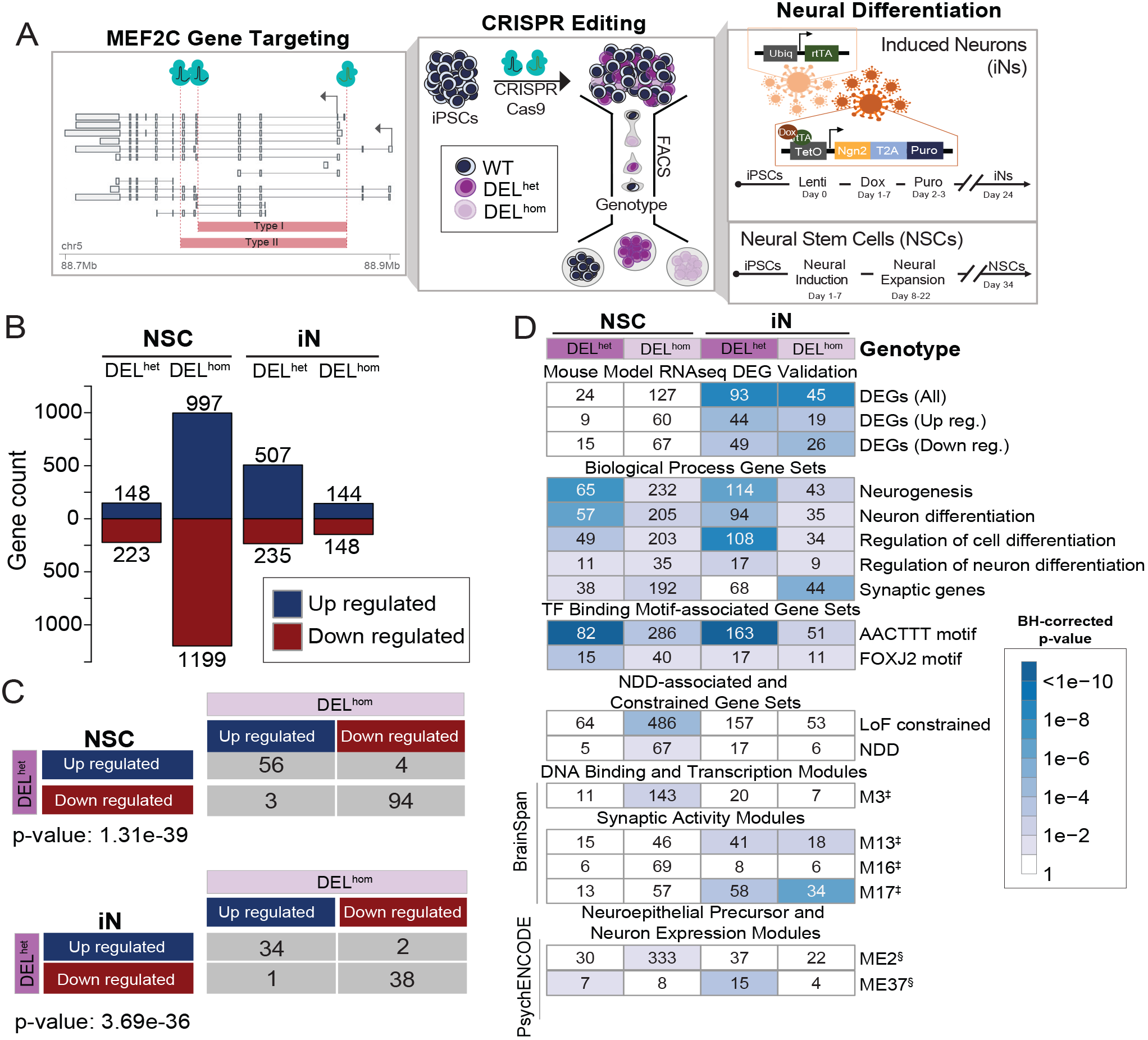
MEF2C deletion yields differential expression of genes enriched for neurodevelopmental and synaptic processes. **A**. Schematic representation of *MEF2C* transcripts and location of CRISPR guide RNAs that introduced two independent deletions (Type I and Type II) into hiPSCs. Fluorescence-activated cell sorting (FACS) was then performed, followed by screening of individual clones that identified both heterozygous (DEL^het^) and homozygous (DEL^hom^) deletions of *MEF2C*, along with unedited wild-type (WT) clones that were exposed to identical CRISPR targeting but remained unedited. Following hiPSC growth, neural differentiation was conducted followed by RNAseq. *MEF2C* CRISPR deletion breakpoints are shown relative to transcripts with >1TPM expression in ≥1 GTEx tissue. Arrows represent positions of canonical transcription start sites of *MEF2C*. **B.** Differentially expressed genes (DEGs) per cell type and genotype. **C.** DEG concordance analysis by cell type and genotype group with statistical significance of shared directionality calculated using Fisher’s exact test. **D.** DEG enrichment for gene sets and modules previously published with functional associations with neurological phenotype, synaptic activity, and *MEF2C* function. Mouse Model RNAseq DEG Validation: Harrington et al, eLife, 2016^40^; Biological Processes Gene Sets: GSEA msigDB^46,47^, Synaptic genes: Syngo v1.1; MEF2C targets: ENCODE LCL ChIPseq^48^; TF Binding Motif-associated Gene Sets: GSEA msigDB^46,47^; LoF constrained: gnomAD^49^; NDD: Neurodevelopmental Disorder-associated genes, Fu, *et al* medRxiv^22^; DNA Binding and Transcription Modules (‡): Parikshak et al, Cell, 2013^41^; Synaptic Activity Modules (‡): Parikshak et al, Cell, 2013^41^; Neuroepithelial Precursor and Neuron Expression Modules (§): Li et al, Science, 2018^42^.

We performed differential expression analysis using DESeq2 and SVAseq, to account for unknown sources of variation in expression data. In this analysis, differentially expressed genes (DEGs) were selected at Benjamini-Hochberg corrected p-values (FDR) < 0.1. In NSCs, we observed a strong zygosity-dependent transcriptional response to *MEF2C* disruption with 371 and 2,196 DEGs in DEL^het^ and DEL^hom^, respectively (**Figure 1B**). By contrast, with iNs we observed 742 DEGs following DEL^het^ that were notable for a predominance of upregulated genes and 292 DEGs in DEL^hom^ cells. Nonetheless, we found highly significant overlap and directionally concordant DEGs between DEL^het^ and DEL^hom^ genotypes in both NSCs (p-value = 1.31e-39) and iNs (p-value = 3.69e-36) (**Figure 1C**).

Gene-set enrichment analysis highlighted differential molecular consequences to *MEF2C* deletion in NSCs and iNs. Haploinsufficiency of *MEF2C* in NSCs led to dysregulation of genes involved in developmental processes including developmental pattern specification, organ morphogenesis, neurogenesis, and neuron differentiation (**Figure 1D**). Additionally, DEGs resulting from homozygous loss of *MEF2C* in NSCs were enriched for LoF constrained genes and gene-sets associated with NDDs from exome sequencing^22^. These LoF and NDD-associated genes are heavily weighted toward genes that display high levels of expression during early neurodevelopment and experience strong negative selection against gene disruptive mutations (**Figure 1D**). In contrast to NSCs, DEGs identified in iNs were significantly enriched for DEGs observed in forebrain excitatory neurons of an Mef2c knockout mouse model published by Harrington et al^40^ (**Figure 1D**). Additionally, DEGs observed in DEL^het^ iNs were enriched for functional terms such as neurogenesis and neuronal differentiation. Homozygous loss of *MEF2C* in iNs also yielded DEGs that were enriched for synaptic genes (**Figure 1D**). When comparing these results to data from the BrainSpan project^41^, which identified neural activity-defining gene co-expression modules using 146 samples from 21 fetal to infant developing brains, iN DEGs were enriched for BrainSpan modules associated with synaptic transmission, synaptic maturation, and genes defined as MEF2C binding targets^41^ (modules M13 and M17; **Figure 1D**). These findings replicated using co-expression modules from a more recent and larger study of 1,230 samples from 48 brains in psychENCODE (Li et al)^42^. We observed the module from Li. et al. (ME37), which includes *MEF2C*, was enriched for DEGs from DEL^het^ iNs (**Figure 1D**). This module from the psychENCODE study demonstrated expression patterns associated with neuron development and was enriched for genes that converged on associations with neurodevelopmental and neuropsychiatric disorders

Overall, the biological pathways and processes shared across DEGs from NSCs and iNs were strongly enriched for neuronal terms including neurogenesis, neuron differentiation, and regulation of cell differentiation and specifically neuron differentiation (FDR < 0.1). Intriguingly, we also observed that DEGs identified in both NSCs and iNs were enriched for AACTTT and FOXJ2 binding motifs (FDR < 0.1). The AACTTT binding motif has previously been associated with the promoter of *MEF2C* and enriched at the promoters of genes involved in neurodevelopment and muscle development^43,44^. The FOXJ2 binding motif has similarly been shown to recruit transcriptional activators that function in early developmental stages^45^. Taken together, these data suggest that the expression signatures associated with LoF mutations of *MEF2C* are consistent with perturbations to highly constrained genes broadly involved in transcriptional regulation during early neural development, as well as genes that display distinct expression patterns in later developmental time points and impact neuronal communication and synaptic functions.

### Altered co-expression of genes in neurodevelopmental and synaptic pathways associated with deletion of MEF2C

We next established modules of co-expressed genes in NSCs and iNs using weighted gene co-expression network analysis (WGCNA; **Figure 2**). In NSCs, four co-expression modules had an eigengene that correlated significantly with MEF2C dosage: violet (P=1e-5), bisque4 (P=5.6e-5), yellow4 (P=8.8e-3), and darkslateblue (P=1.2e-3) (**Figure 2A, C, E**). Yellow4 genes, which showed increased expression with *MEF2C* loss, were notably enriched for processes of heart morphogenesis^13,50–52^ and RHO GTPase activation^53^ (FDR <0.05), both of which have been described previously in relation to MEF2C function. Furthermore, we also considered co-expression modules with an eigengene that correlated with DEL^het^ MEF2C loss alone. Module navajowhite2 (P=5.30e-1) contained genes significantly up-regulated only in DEL^het^ NSCs and was enriched for terms related to synapse assembly and organization.

**Figure 2.**
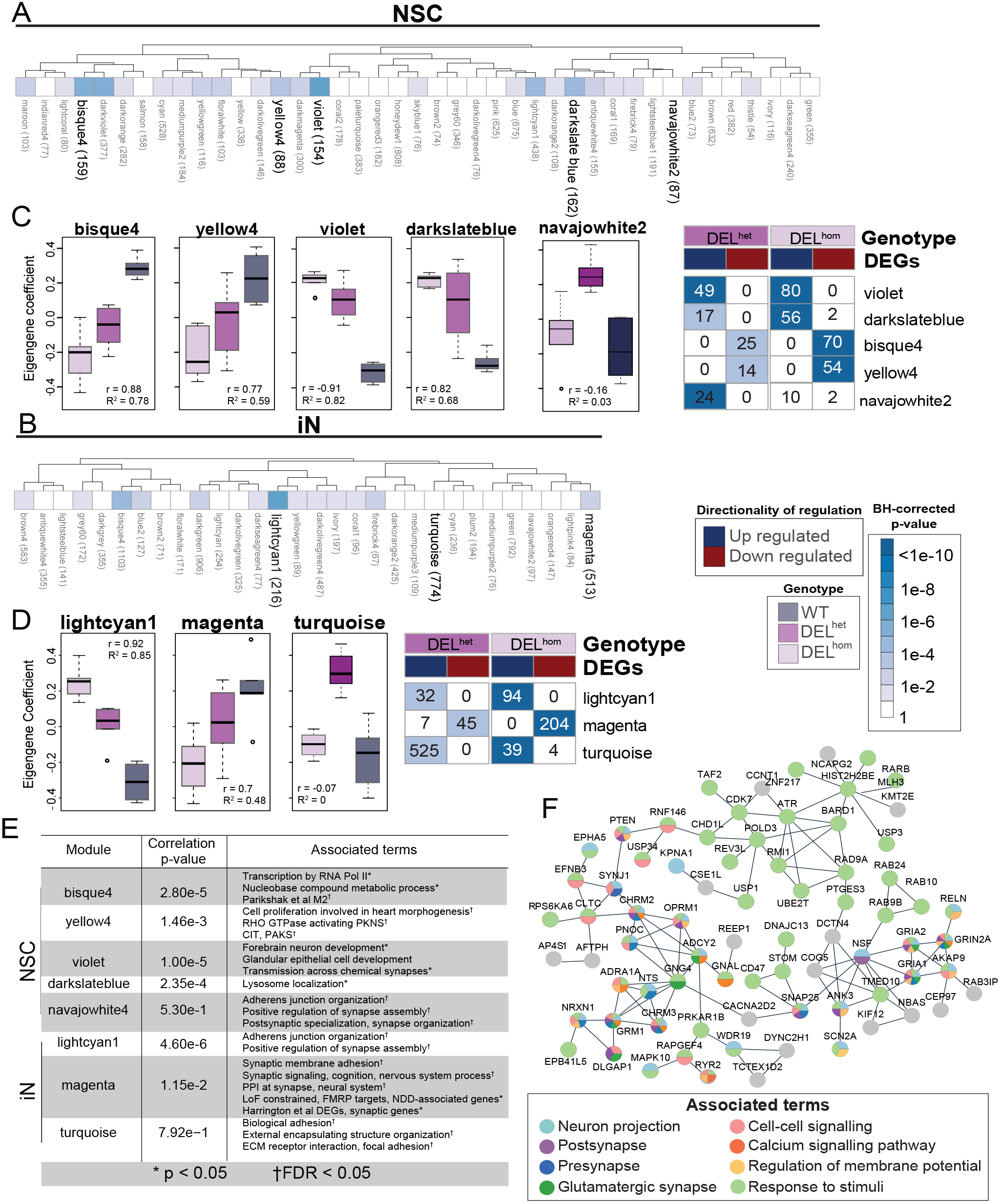
Co-expression modules enriched for constrained genes and synaptic activity are correlated with MEF2C disruption in NSCs and iNs. Dendrogram displaying all modules identified from WGCNA. Modules demonstrating statistically significant correlation with MEF2C genotype (e.g. allelic dosage) are bolded for NSCs (**A**) and iNs (**B**) respectively with p-value of correlation significance indicated by color shade. Boxplots demonstrating each eigengene’s coefficient of correlation per genotype per co-expression module, and corresponding heatmaps, shown for NSCs (**C**) and iNs (**D**). **E**. Associated terms with significant enrichment per highlighted module. **F**. The PPI network of highly connected genes (n=76) of the magenta module with functional annotations. The node colors indicate the functional classes for each protein and the edge represents interaction.

Co-expression analysis in iNs isolated two modules whose eigengenes significantly correlated with *MEF2C* dosage: lightcyan1 (P=1.5e-7) and magenta (P=1.9e-3) and one module turquoise with neuronal function and only affected by single copy edit of MEF2C (P=7.92e-1) (**Figure 2B, D, E**). Genes within module lightcyan1 were enriched for DNA damage repair-associated nucleotide patch replacement (FDR <0.01). In addition to having an essential role in cardiac and neurodevelopment, previous reports have also described *MEF2C* as serving lineage specific roles in regulation of DNA damage repair^54,55^. The genes co-expressed in the magenta module revealed a far greater emphasis on alterations to synaptic functions, being enriched for synaptic membrane adhesion, synaptic signaling, and protein-protein interactions at the synapse (FDR < 0.01). The encoded proteins (n=513) were mapped to a PPI network using StringDB^56,57^ in Cytoscape^58–60^, resulting in a network with 150 members (confidence score > 0.7, evidence=experimental/database, p-value < 1e-16). The proteins from 76 genes contained within module magenta formed a network associated with neuronal and synaptic terms (**Figure 2F**). Module magenta also included genes associated with monogenic forms of epilepsy and other NDDs, such as GRIN2A, SCN2A, and GRIA2, providing evidence of molecular convergence for these genotypically distinct but strongly synaptic activity-associated disease genes^61–63^.

### MEF2C direct disruption yields changes to synaptic firing and synchrony in human neural models as measured by multi-electrode array (MEA)

Given the strong transcriptional changes associated with deletion of *MEF2C* in hiPSC-derived neurons that converged on synaptic activity, we sought to functionally validate this association by defining electrophysiological changes in neurons using multi-electrode array (MEA; **Figure 3, Supplementary Section 2.1, Supplementary Figures 5-7**). We differentiated heterozygous and homozygous *MEF2C* lines and matched controls to iNs and observed a statistically significant (based on t-test) down-regulation of synaptic activity in iNs. We observed statistically significant reductions in firing rate relative to wildtype for both DEL^het^ (26%, P=1.4e-5) and DEL^hom^ (31%, P=1.3e-2) (**Figure 3A**). We also observed statistically significant reductions to spike count relative to wild-type for both DEL^het^ (17%, P=6.7e-3) and DEL^hom^ (48%, P=5.1e-5) (**Figure 3B**). Additionally, we observed statistically significant reductions to synchrony, a measure of uniformity of neuronal firing bursts, relative to wildtype for both DEL^het^ (32%, P=7.1e-3) and DEL^hom^ (73%, P=9.6e-9) (**Figure 3C)**. While we observed clear changes in firing rate, neither DEL^het^ (P=0.68) nor DEL^hom^ (P=0.30) loss of *MEF2C* resulted in significant changes to network burst oscillation (**Figure 3D**). These data provided a complementary measure of *MEF2C* direct disruption resulting in altered synaptic activity.

**Figure 3.**
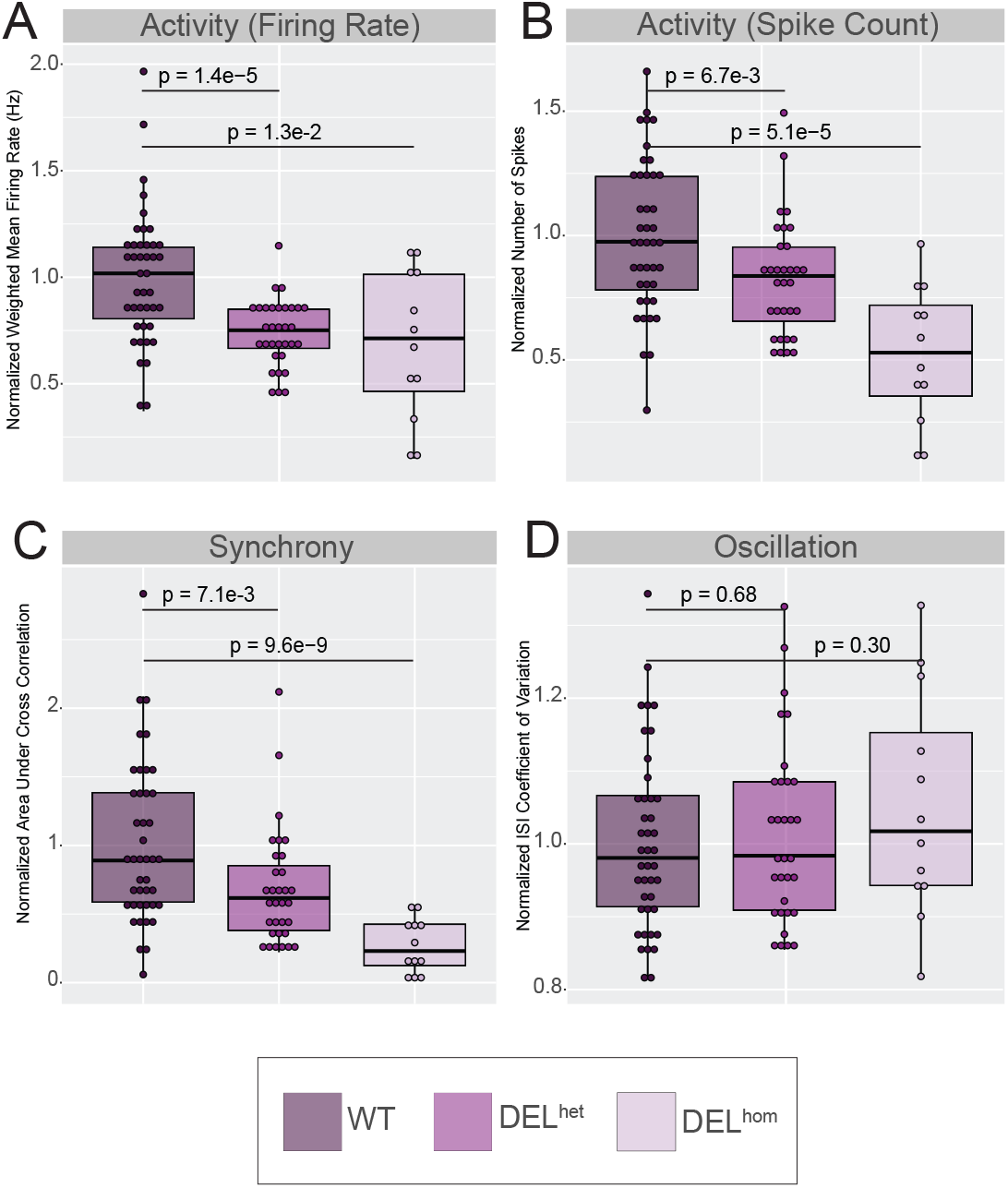
Deletion of MEF2C is associated with reduced synaptic activity and synchrony in neurons. Each datapoint represents a single replicate well. Each resultant measurement was normalized per well against the wild-type mean per plate and normalized data points from two replicate plates plotted. **A.** Normalized weighted mean firing rate (Hz) (Firing Rate). **B**. Normalized number of spikes (Spike Count). **C**. Normalized area under cross correlation (Synchrony). **D**. Normalized ISI (interspike interval) coefficient of variation (Oscillation). P-values calculated using t-test against normalized wells per genotype.

### Dissecting the three-dimensional chromatin topology and regulatory interactions within the 5q14.3 locus

Our prior analyses from whole genome sequencing of individuals with NDDs harboring BCAs^38^, and other recent studies^1,64–69^, have suggested that cis-regulatory disruption by noncoding SVs may underlie NDD phenotype association within the 5q14.3 locus beyond direct LoF *MEF2C* mutation. Collectively, these seven studies have reported CNV and BCA breakpoints 200-500kb distal to *MEF2C* in 13 distinct cases presenting with phenotypes consistent with *MEF2C* haploinsufficiency such as NDD, epilepsy, and hypotonia. As a result, enhancer-promoter decoupling by disruption to 3D chromatin organization has emerged as a mechanistic hypothesis for indirect *MEF2C* disruption. We therefore performed a comprehensive and systematic dissection of the TAD and loop organization of the 5q14.3 region in human neural models. From analyses of existing 2D elements enhancers as well as 3D element like boundaries and structural protein ChIP annotations (CTCF and SMC3) from published datasets^31,70^, we defined both 2D and 3D elements with evidence for a role in MEF2C regulation (**Supplementary Section 3.1, Supplementary Figures 8-9**). We then sought to determine the overarching 3D functional architecture responsible for orchestrating gene-enhancer interactions by generating an allelic series of deletions targeting 2D and 3D functional elements within the 5q14.3 locus (**Figure 4**).

**Figure 4.**
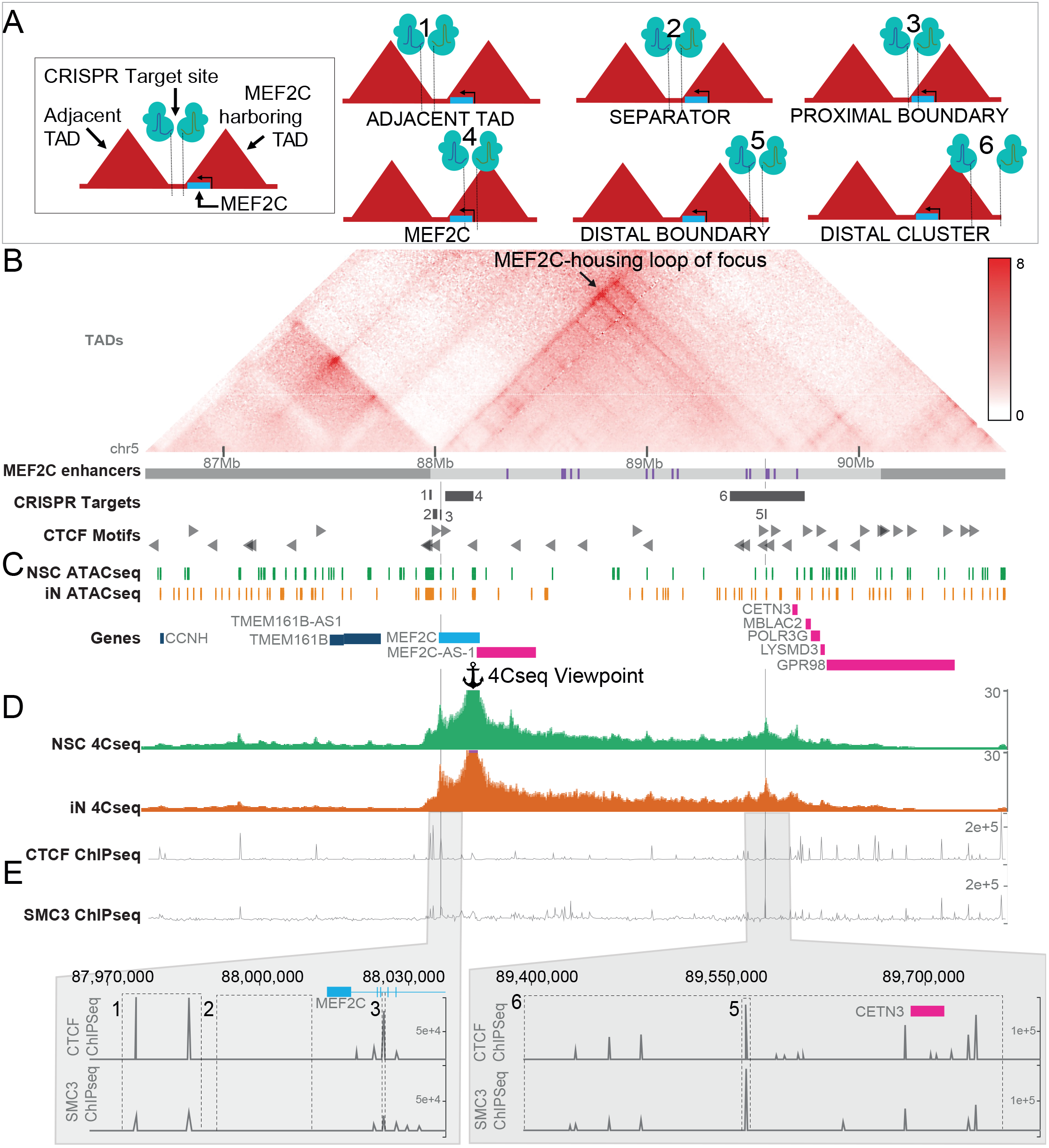
Design of iPS CRISPR deletion models. **A**) Depiction of CRISPR targets relative to MEF2C and local 3D structures. **B**) Knight-Ruiz normalized Hi-C map from GM12878 wildtype LCLs shown^31^. Local TAD annotations^70^ and putative MEF2C enhancers^71^ denoted as grey and purple bars respectively. Black arrow within the Hi-C map and vertical grey lines highlight the MEF2C-housing loop of focus targeted in this study. **C**) Open chromatin regions identified by ATACseq in wildtype NSCs and iNs. **D**) Aggregate contacts from 4Cseq in wildtype NSCs and iNs shown relative to the viewpoint within the MEF2C promoter. **E**) Deletion positions of 3D topology boundaries guided by ChIPseq for structural proteins CTCF and SMC3 in GM12878 LCLs.

We annotated topological structures from LCLs using Hi-C data and integrated CTCF and SMC3 binding sites indicated by LCL ChIP-seq^31,70^. Dual guide CRISPR/Cas9-based genome editing guides were then designed to engineer a series of deletions of the candidate 3D elements as outlined in Figure 4. We targeted deletion of four genomic sites, including the proximal and distal boundaries of the *MEF2C*-containing loop as the key experimental edits (referred to as “Proximal Boundary” and “Distal Boundary”, respectively), and two ‘negative control’ edits, namely the boundary of the TAD adjacent to MEF2C (“Adjacent TAD”), and the genomic sequence spanning the MEF2C-containing and adjacent TAD with no occupied CTCF binding sites in 133 cell/tissue samples from ENCODE (“Separator”). We focused on this MEF2C loop structure as opposed to the larger TAD given that it was largely cell type invariant in both the above resources and additional cell types, as well as the strength of contact with MEF2C and its higher resolution map of the 3D organization encompassing the MEF2C-relevant enhancers (**Supplementary Section 3.1, Supplementary Figures 8-9**). Differentiated hiPSC-derived iN and NSC CRISPR models were generated for each of these four deletion models using six replicates per DEL^het^ and DEL^hom^ genotype, as well as six control clones that were exposed to the CRISPR conditions but not edited (**as described in Materials and Methods**). Differentiation of these iPSCs established 204 individual neuronal lines representing systematic disruption to functional elements within the 5q14.3 locus (**Supplementary Section 3; Supplementary Figures 14-20**).

### Distal Boundary deletion does not result in marked change to MEF2C expression

We observed that DEL^het^ and DEL^hom^ of the Separator and Adjacent TAD boundary deletions resulted in no substantial changes in *MEF2C* expression or to contacts between the MEF2C promoter and published enhancers of MEF2C^71^ based on UMI-4C, as expected (**Figure 5, Supplementary Section 3.6, Supplementary Figures 22-23, 26-29**).

**Figure 5.**
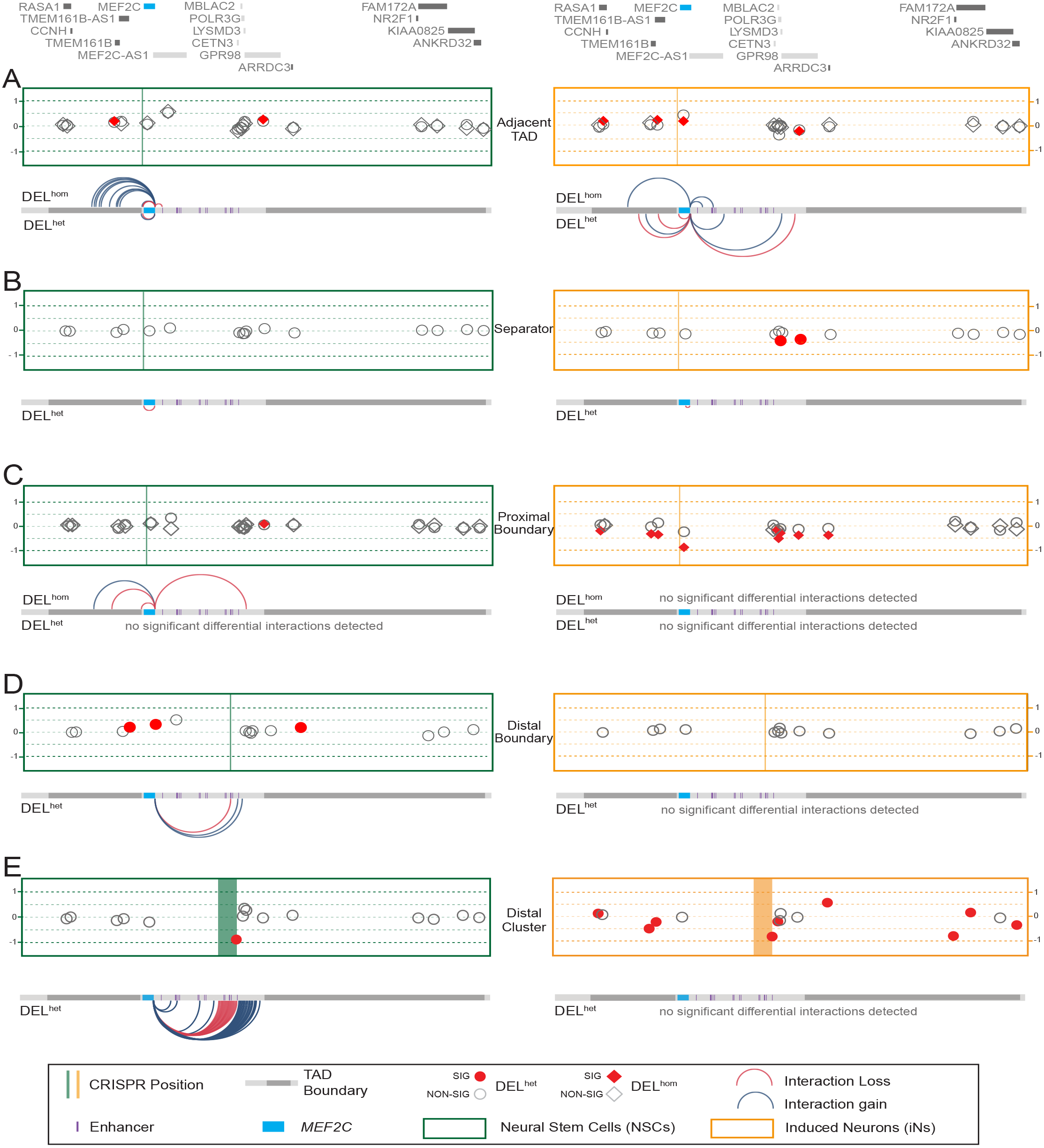
Resultant MEF2C expression and long-range contact changes from 5q14.3 non-coding element disruption cannot be predicted a priori based on cell type, genotype, or functional element type. Y axis shows Log2 fold change of differential expression for *MEF2C* and 15 local genes as defined by their inclusion within the *MEF2C*-containing TAD, TAD immediately proximal, or distal. The significantly differentially expressed (locally corrected FDR<0.1) genes in NSCs and iNs are shown in red and non-significant genes shown outlined in grey. Differential expression of each gene is annotated by a circle for DEL^het^ deletions (all CRISPRs) and a diamond for DEL^hom^ deletions (able to be generated in a subset of CRISPRs). Position of CRISPR deletion annotated by vertical line. Differential contacts with the *MEF2C* promoter in each CRISPR line relative to matched wild-type indicated by curved lines (**Supplementary figure 25-29**). Lines included for significant contact changes with FDR<0.1 based on a sliding window of 5kb with 200bp step. Note: MEF2C-AS1 is not expressed in iNs.

The Distal Boundary deletion served to directly test the necessity of 3D chromatin topology for robust expression of *MEF2C*^1,38,64–67^. Surprisingly, deletion of this site resulted in no change in MEF2C protein expression and a slight increase (25% compared to wildtype) in mRNA expression for DEL^het^ in NSCs (log2FC = 0.33 ; FDR <0.1 when correcting for multiple testing within the TAD, **Figure 5; Supplementary Figure 23**). Aside from *MEF2C*, we also considered expression of genes expressed in NSCs or iNs that were located within either the MEF2C-containing TAD, the TAD directly proximal, or the TAD directly distal, which we refer to herein as “local genes”, using TADs as defined by Dixon et al^70^. 15 genes (not including *MEF2C*) met this criterion for local gene inclusion in NSCs, while 14 were considered in iNs given that MEF2C-AS1 was not expressed in this cell type. We observed no altered expression of any local genes aside from *MEF2C* in Distal Boundary deletion NSCs. To further explore this finding, we employed allele-specific UMI-4C anchored at the *MEF2C* promoter (**Figure 5, Supplementary Section 3.6, Supplementary Figures 26**). These analyses identified an increase in significant (FDR<0.1) contacts between the *MEF2C* promoter and two separate 6kb windows harboring CTCF binding sites immediately distal to the deletion position, suggesting CTCF motif redundancy may buffer deletions to canonical 3D boundary elements, preventing strong dysregulatory effects. In DEL^het^ matched iNs with Distal Boundary deletions, no significant expression or contact changes were observed for *MEF2C* or any other local genes.

To further evaluate this largely negative result, we tested the hypothesis of regulatory element redundancy buffering against pronounced expression effects on *MEF2C*. We did so by deleting what we referred to as the Distal Cluster, a 354kb region that includes six annotated enhancers (e11-e16), five directly oriented CTCF binding site motifs, the Distal Boundary, and one local gene, *CETN3* (**Figure 4**). We note that the generation of this large CRISPR deletion was performed directly in NSCs (by comparison to the hiPSC stage prior to differentiation for all other models), and is thus a technically distinct validation experiment rather than directly comparable to the models in the initial hypothesis test. Nonetheless, in these analyses we observed no significant differential expression of *MEF2C* upon DEL^het^ deletion of the Distal Cluster region when compared to matched wildtypes in either NSCs or NSC-derived iNs (**Supplementary Figures 23-24**). Considering UMI-4C data from the viewpoint of the MEF2C promoter in these lines, we observed a significant (FDR<0.1) increase in contacts with four sites within 5q14.3 alongside a contiguous increase with the 320kb region immediately distal to the deleted region in NSCs. Considering local genes not directly disrupted by the deletion, we observed a cascade of significant (FDR<0.1) dysregulation of 8/15 genes in 5q14.3 locus in iNs, with no genes significantly dysregulated in matched NSCs. Together, these analyses suggest that deletion of the distal boundary of the *MEF2C* encompassing loop is insufficient to indirectly dysregulate this NDD gene.

### Proximal Boundary deletion yields reduction of MEF2C expression and synaptic activity in iNs

In contrast to the weak or largely negative results observed for deletion of the Distal Boundary, deletion of the Proximal Boundary, which is located in an intron of *MEF2C*, had marked effects on the gene’s expression that appeared to be genotype and cell-type dependent. NSCs harboring DEL^het^ or DEL^hom^ of the Proximal Boundary did not display differential expression of *MEF2C*, though protein expression was significantly reduced in DEL^hom^ (49% relative to controls; p-value = 5.7e-4, **Supplementary Figure 22-23**). These DEL^hom^ NSCs also displayed differential contacts with the *MEF2C* promoter at three sites, two of which were significantly increased and one significantly decreased (locally corrected FDR<0.1). Moreover, homozygous deletion of the Proximal Boundary in iNs resulted in pronounced down-regulation of *MEF2C*, five genes localized homozygous deletion of the Proximal Boundary in iNs resulted in pronounced down-regulation of MEF2C, five genes localized to the *MEF2C*-containing TAD, and three genes within the TAD proximal to the *MEF2C*-containing TAD (locally corrected FDR < 0.1; **Figure 5**). This consistent down-regulation thus extended up to 3Mb from the site of the deletion, and was also dosage-dependent as DEL^hom^ deletion resulted in significantly reduced expression compared to unedited and DEL^het^ cells (**Figure 5**). These positional effects were particularly strong for *MEF2C*, and DEL^hom^ deletion largely recapitulated the reduction observed with heterozygous direct gene deletion (e.g. 45% reduction, locally corrected FDR = 1.9e-9, genome-wide FDR = 6.3e-8) (Fi**gure 5**).

We next explored transcriptional and functional commonalities between direct *MEF2C* disruption and indirect expression reduction by Proximal Boundary deletion. We observed shared DEGs from direct *MEF2C* disruption in both NSCs and iNs with Proximal Boundary deletion (**Figure 6**). We also observed shared functional pathways in iNs related to synaptic activity, neurodevelopment, and neural differentiation between these distinct coding and noncoding functional mutations (**Figure 6**). The Proximal Boundary DEL^hom^ and direct *MEF2C* deletion iN DEGs also shared significant enrichment of terms such as axon development alongside sharing an enrichment for genes with promoter containing the sequence motif AACTTT, a binding motif in genes involved in neurodevelopment and muscle development^43,44^. Given our demonstration of shared neuronal pathways between direct MEF2C deletion and Proximal Boundary deletion lines, we next tested whether this noncoding regulatory Proximal Boundary deletion replicated the synaptic deficits observed with MEF2C deletions. From these analyses, we observed a similarly strong reduction in synaptic activity for both DEL^het^ and DEL^hom^ iNs when compared to matched wildtype clones over the differentiation time course measurements (**Figure 6**; Day31 - Day43). The pattern and significance detected for spike number and number of bursts over time was also consistent with the MEF2C deletion MEA time course experiment, suggesting a reproducible synaptic phenotype associated with both direct deletion and noncoding regulatory alterations to the MEF2C locus (**Supplementary Section 3.6, Supplementary Figures 30-32**).

**Figure 6.**
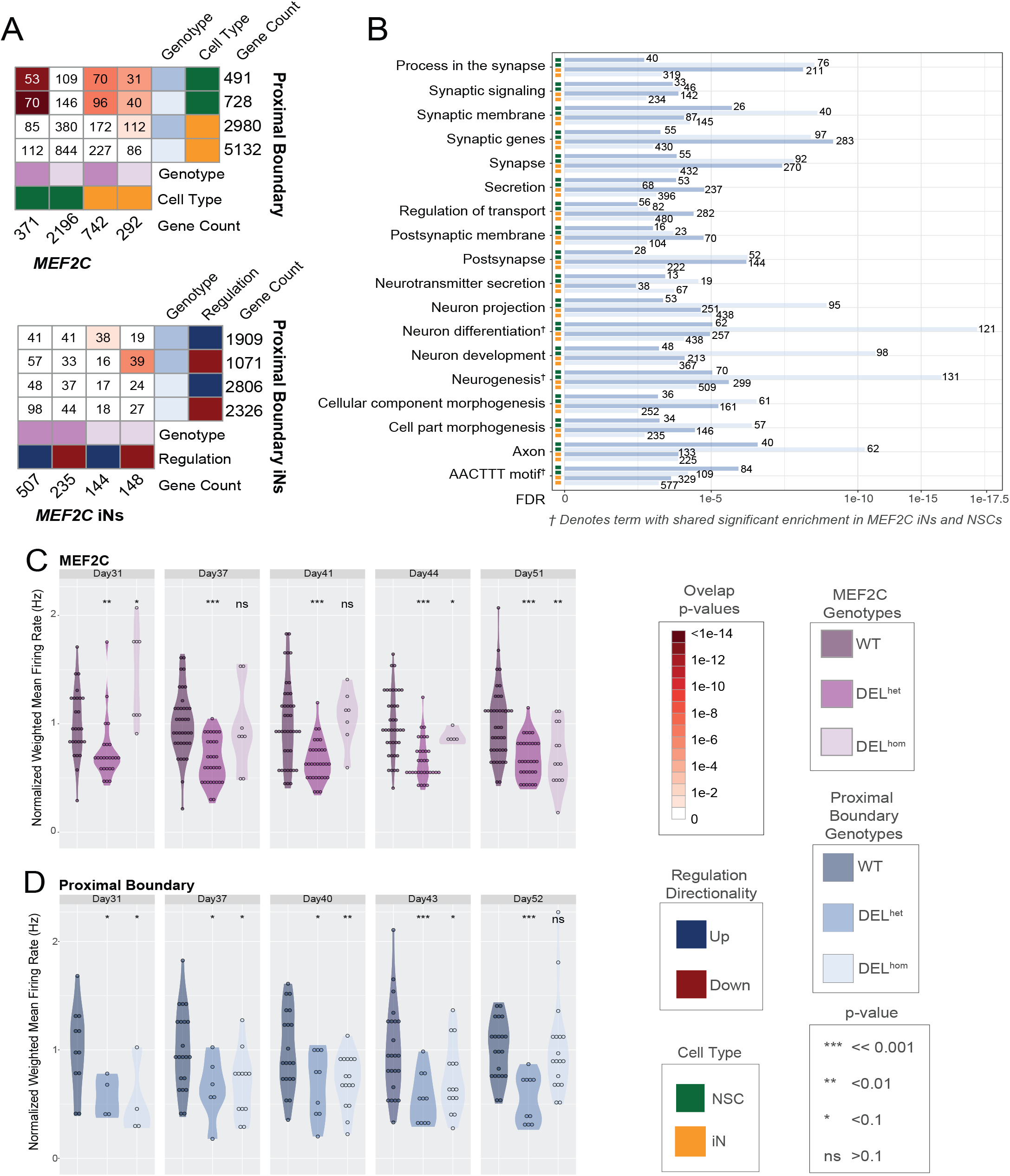
Proximal Boundary disruption yields disruption of genes involved in neuron differentiation and alters synaptic activity. **A**. DEG comparison between Proximal Boundary and *MEF2C* direct disruption lines (top: both NSCs and iNs, bottom: iNs only). **B**. Functional enrichment of Proximal Boundary DEGs. Terms shared in MEF2C iNs and NSCs in both genotypes are also noted. Normalized weighted mean firing rate over time as measured by MEA shown for *MEF2C* iNs (**C**) and Proximal Boundary iNs (**D**), respectively.

## DISCUSSION

We present an initial survey of the transcriptional and electrophysiological consequences of *MEF2C* gene deletion and indirect regulatory changes introduced by alterations to 5q14.3 TAD and loop chromatin topology in human-derived in vitro neural models. Direct LoF mutations (PTVs, deletions, translocations) within MEF2C have repeatedly been associated with NDD risk.^2,3,5,6,10,65,66,72–75^ A variety of murine knockout models, including conditional knockout of *MEF2C* in varying neural cell types and developmental time points, have collectively suggested a role for *MEF2C* in neuronal function,^13,14,16,40^ but no studies have evaluated *MEF2C* LoF mutations in human-derived cell lineages. In the study presented here, we engineered an allelic series of CRISPR mutations within *MEF2C* and its putatively regulatory 3D functional elements, and demonstrated that both direct and indirect alterations to *MEF2C* expression converge on themes related to neurodevelopment and synaptic activity.

Recently, specific studies that have focused on the knockout of *SMC3* in human cell models^29^, genome rearrangements in drosophila^36^, and TAD disruption at the Shh locus^76^ have demonstrated the uncoupling of regulatory changes associated with gene expression and alterations to long-range chromatin contacts. Our data further support those studies, as we observed alterations to 3D topological organization that occur without concomitant changes in gene expression. These demonstrations are in contrast to the hallmark examples of TAD disruption underlying dysregulation of a gene associated with a human disease phenotype3^1,33–35^. Most of the CRISPR models targeting 2D and 3D noncoding element deletions generated in this study demonstrated modest or no functional impact on expression of *MEF2C* and genes encompassed within the adjacent TADs. Furthermore, most CRISPR models also did not result in major changes to long-range contacts with the *MEF2C* promoter. We did note some robust changes in contact patterns from UMI-4C conducted in our NSC CRISPR lines, but most observed differential contacts did not engage previously validated enhancers of *MEF2C*. These findings further illustrate the complexity associated with the dynamic interactions that result from SV alterations of 3D topology and regulation of gene function.

Our approach weighted elements contributing to 3D genome organization given the previous reports highlighting TAD disruption as a putative indirect cause of *MEF2C* haploinsufficiency^38, 65^. In our most recent WGS study of 406 individuals with developmental disorders that harbored a BCA, as well as 304 BCAs from control individuals, the 5q14.3 locus continues to display a genome-wide significant enrichment of noncoding SV breakpoints that localize distal to *MEF2C*. The homozygous Proximal Boundary deletion caused a two-fold reduction of *MEF2C* expression that was comparable to haploinsufficiency resulting from heterozygous deletion. The absence of a coding SNP within MEF2C prevented the determination of whether up-regulation of *MEF2C* on the wild-type allele prevented observed differential expression in heterozygous deletion of Proximal Boundary lines. This region has been recently reported as a cis-regulatory site that was demonstrated to reduce *MEF2C* expression following CRISPRi in K562 cells77. The shared enrichment of gene sets related to neurogenesis and neuronal differentiation between both Proximal Boundary and *MEF2C* DEGs demonstrates transcriptome dysregulation by both direct and indirect *MEF2C* disruption that converge upon biological processes of relevance to NDDs. We also observe significant overlap between indirect disruption of *MEF2C* by Proximal Boundary deletion in NSCs and direct *MEF2C* disruption in iNs, suggesting dysregulatory effects may involve temporal or cell type specific dysregulation. Furthermore, the shared enrichment of genes with AATCCC binding motifs within their promoters, which was previously associated with genes involved in neuronal differentiation, including *MEF2C*, alongside the comparable reductions in synaptic activity as measured by MEA, suggest shared functional consequences from direct and indirect *MEF2C* disruption. However, the deletion of the Distal Boundary did not recapitulate these results. While the data are therefore unambiguous of a localized enrichment of disease associated noncoding SV breakpoints spanning the *MEF2C*-containing 3D organization, the models that displayed the greatest transcriptional and functional consequences were not those that were predicted a priori based on the localization of the SV breakpoints alone. The noncoding regulatory mechanisms that govern NDD risk in this region thus remain elusive and appear to be cell-type and genotype specific. The deletions introduced here do not result in the same degree of genome topological rewiring as would be predicted from the translocations and inversions observed in the BCA cases, and the functional changes observed are not consistent across cell types. These analyses illustrate the highly complex regulatory architecture of alterations to chromatin topology and emphasize the significant challenges for computational prediction of the regulatory features of SVs^78–80^.

We present the first CRISPR engineered isogenic allelic series of *MEF2C* disruption in human-derived in vitro models and functional characterization of resultant transcriptional and electrophysiological effects. These studies also uniquely dissect the parallel consequences of direct and indirect alterations to *MEF2C* by cis-regulatory disruption, revealing that some noncoding mutations can recaptiulate synaptic deficits and transcriptional signatures observed from direct deletion of this gene underlying the well-established 5q14.3 microdeletion syndrome. While direct deletion of *MEF2C* revealed functional changes in human neurons that were consistent with previous studies in mouse models^14–17,40^, our results following SV alteration to 3D organization of 5q14.3 underscore the complexity of regulatory interactions at this locus, and more broadly in defining the features associated with the functional impact of noncoding SVs genome-wide. These data clearly demonstrate that alteration to annotated boundaries of 3D regulatory architecture encompassing established human disease genes is insufficient evidence to presume alterations to gene regulation or phenotypic impact. Future in silico, in vitro, and in vivo studies targeting more loci and classes of SVs with designs that either recapitulate case rearrangements or are agnostic to regulatory element type will be necessary to further expand these findings into systematic analysis of the features associated with 3D genome architecture and noncoding disease association.

## METHODS & SUPPLEMENTARY INFO

Detailed methods and supplementary information for this manuscript have been provided in a separate document, which will be linked directly from *bioRxiv*.

## ACKNOWLEDGMENTS

We would like to thank Maris Handley and the staff of the Harvard Stem Cell Institute FACS core at Massachusetts General Hospital for helpful discussion and technical assistance with cell sorting for iPS clone generation. We would also like to thank Lies Vantomme for excellent technical assistance.

This research was supported by grants from the National Institutes of Health: P01GM061354, R01HD096326, R01MH115957, R01MH123155, U01HG011755, R03HD099547, R01NS093200, K08NS117891, T32GM007748, G044615N and 1520518N. Support was also provided by the Simons Foundation for Autism Research Initiative (#573206). KM was supported by the National Science Foundation Graduate Research Fellowship Program (NSF GRFP) doctoral fellowship. RY was supported by the Mass General Hospital Fund for Medical Discovery. ED and SV were respectively supported by a doctoral and postdoctoral fellowship of the Research Foundation Flanders (FWO). PB was supported by 1K08NS117891 and T32GM007748. MMO was supported by the Autism Speaks Postdoctoral Fellowship.

## AUTHOR CONTRIBUTIONS

Study design: KM, RY, ED, BM, JFG, SV, MT; Experiments: KM, ED, PMB, MMO, RB, BC, KK, NB, DT, MS, ML. Data analysis: KM, RY, ED, SE, DG, CL; Data interpretation: KM, RY, SE, ED, MT; Manuscript writing: KM, RY, SE, ED, JFG, SV, MT. All authors have reviewed and accepted final version of the manuscript.

## COMPETING INTERESTS

M.E.T. receives research funding and/or reagents from Levo Therapeutics, Microsoft Inc, and Illumina Inc. KM initiated employment with Tornado Bio during the writing of this manuscript and is at present an employee of the company. All other authors declare no competing interests. All other authors declare no competing interests.

