## Supplementary material for "Transcriptional and functional consequences of alterations to *MEF2C* and its topological organization in neuronal models": Methods_SupplementaryInformation_Supplementaryfigures

Kiana Mohajeri[1-4*], Rachita Yadav[1-3*] et al.
*Full list of authors is available in the main text*

#### Table of Contents

***Materials and Methods*** ***3***
**iPSC Culture:** **3**
**in silico CRISPR guide selection:** **3**
**CRISPR/Cas9 Transfection:** **3**
**FACS and Consolidation for Isolation of Clonal CRISPR Populations:** **4**
**Genotyping Clonal CRISPR Populations:** **4**
**Selection of TRA-1-60 Positive iPSC CRISPR Lines:** **4**
**Differentiation of Induced Neurons (iNs):** **5**
**Differentiation of Neural Stem Cells (NSCs)** **5**
**RNA Extraction** **6**
**Whole Transcriptome Sequencing (RNA-seq) Library Preparation** **6**
**RNAseq data processing** **6**
**Differential Gene Expression** **7**
Differential Exon Expression 7
Co-expression Analysis 7
Functional enrichment 7
**ATACseq Sample and Library Preparation from WT iNs and NSCs** **7**
**ATACseq Peak Calling from WT iNs and NSCs** **8**
**Differentiation Marker Staining** **8**
**Western Blot** **8**
**Multi-Electrode Array (MEA)** **9**
**4C Sample Preparation** **9**
4C Library Preparation and Sequencing 10
4C Data Processing – Delta Map Generation 10
4C Data Processing – FDR-Corrected Differential Contact Annotation 10
***Supplemental Methods*** ***11***
**Section 1: *MEF2C* direct disruption molecular QC:** **11**
Section 1.1: Design of MEF2C deletion iPSC lines: 11
Section 1.2: Deletion confirmation in *MEF2C* iPSC lines: 11
**Section 1.3: Confirmation of MEF2C expression reduction in direct disruption CRISPR edits:** **13**
Section 1.4: Differentiation marker expression validation in *MEF2C* iNs and NSCs: 15

**Section 2: *MEF2C* direct disruption multi-electrode array (MEA): 16**
Section 2.1: Analysis of *MEF2C* direct disruption iNs for multi-electrode array (MEA): 16

**Section 3: Characterization of 5q14.3 positional effect CRISPR targets 21**
Section 3.1: Determination of Proximal and Distal Boundary targets for CRISPR 21
Section 3.2: Design of five positional effect CRISPR target positions within 5q14.3 locus 26
Section 3.3: Genotype characterization of 5q14.3 iPSC CRISPR lines 27
Section 3.4: Differentiation marker expression validation in positional effect iNs and NSCs 31
Section 3.5: MEF2C protein expression in positional effect CRISPR NSCs 35
Section 3.6: mRNA expression of local genes within 5q14.3 in CRISPR iNs and NSCs 40
Section 3.7: Identification of distal contacts with *MEF2C* promoter using UMI-4C 40
Section 3.8: Proximal Boundary CRISPR multi-electrode array (MEA) assessment of synaptic activity 45

***Supplementary Tables 49***

***Supplementary References 49***

#### Supplementary Figures:

### Materials and Methods

#### iPSC Culture:

GM08330 human iPSCs between P40-50 were used for all CRISPR experiments. Cells were grown in feeder-free culture using Matrigel hESC-Qualified Matrix (Corning, 08-774-552) with Essential 8 media (Gibco, A1517001). Cells were passaged using ReLeSR (STEMCELL Technologies, 05873). mFreSR freeze media (STEMCELL Technologies, 05855) was used for cryopreservation. Y-27632 dihydrochloride (MedChem Express, HY-10583) was added to media at 1mg/mL for up to 24 hours for initial plating and at various points in the engineering protocol as indicated below.

#### in silico CRISPR guide selection:

Dual guides were designed with multiple considerations to maximize on-target cutting efficiency for deletion generation while minimizing both off-target hits as well as off-target effects. Guides with NGG PAM for Cas9 were identified using the MIT CRISPR design tool (http://crispr.mit.edu). Dual guides were selected to encompass each targeted functional element within the intended deletion with at least 200bp of buffer sequence. Buffer sequence was often larger given guide placement limitations from predicted off targets. Off-target characterization was conducted per guide using Off-Spotter (https://cm.jefferson.edu/Off-Spotter/). We restricted to guides without identified coding, lncRNA, rRNA, or tRNA off-target hits with ≤4 mismatches between the guide and the off-target position. Guide sequences used are listed in **Supplementary Table 1**.

#### CRISPR/Cas9 Transfection:

For the generation of CRISPR lines for 5/6 target regions (Separator, Adjacent TAD, Proximal Boundary, Distal Boundary, and *MEF2C* direct gene deletion), iPSCs were transiently transfected with breakpoint targeting guides and S.p. Cas9 ribonuclear protein as ribonuclear proteins (RNP). Alt-R S.p. Cas9 Nuclease V3 (IDT, 1081058) was diluted to 25µM in 20mM HEPES Buffer and 150mM KCl. Custom designed Alt-R crRNAs (IDT) and Alt-R tracrRNA (IDT, 1072532) were diluted to 25µM and annealed using manufacturer recommendations to create crRNA::tracrRNA duplexes. Single cell iPSC suspension of Y-27632 dihydrochloride-treated cells was generated using Accutase (STEMCELL Technologies, 07920) and filtering through a 40µM mesh strainer (Falcon, 08-771-1). Diluted Cas9 (1µg) and annealed guides (37.5µM total) were combined with 1µg pmaxGFP plasmid (Lonza, VPH-5012) and nucleofected into iPSCs ($5x10^5$ cells per transfection). Nucleofections were conducted with an Amaxa Nucleofector II, program A-033 using Human Stem Cell Nucleofector Kit 1 (Lonza, VPH-5012), following manufacturer's instructions. Nucleofected cells were plated onto matrigel-coated 12-well plates, maintained in E8 media supplemented with Y-27632 dihydrochloride at 1mg/mL for 24 hours. Cell-containing wells were replenished with E8 media daily with subsequent FACS 36-48 hours post-transfection.

Given that the Distal Cluster edited lines were generated as a follow-up to the first 5 CRISPR sets, RNPs were delivered using lipofection given reduced recovery associated with the generation of this large deletion with nucleofection. As was conducted with the other CRISPR sets, custom designed Alt-R crRNAs (IDT) along with commercially available Alt-R tracrRNA (IDT, 1072532) were used. Single cell, P6 NCAM+ NSC suspension of Y-27632 dihydrochloride-treated cells (0.5mg/mL media) was generated using Accutase (STEMCELL Technologies, 07920) and filtering through a 40µM mesh strainer (Falcon, 08-771-1). NSCs were plated at 30-50% confluence on 12-well plates 3-5 hours prior to addition of lipofection reagent. Lipofectamine Stem reagent (Invitrogen, STEM00003) was combined with diluted Alt-R S.p. Cas9 Nuclease V3 (1µg; IDT, 1081058), EGFP mRNA (0.2µg; TriLink Biotechnologies, L-7201) and crRNA::tracrRNAs (37.5uM) annealed following manufacturer protocol as indicated above. Lipofection reagent was maintained on cells for 24 hours. Cell-containing wells were replenished with E8 media daily with subsequent FACS 36-48 hours post-transfection. Transfection and isolation of the Distal Cluster deletion was attempted starting with both wildtype iPSCs and NSCs. Distal Cluster deletion was not recovered in iPSC background but was generated in NSC background. NSCs within 3-5 passages of transfection were utilized for subsequent steps (expansion for RNAseq, 4Cseq, marker staining, transduction for NSC-derived iN generation, etc.).

#### FACS and Consolidation for Isolation of Clonal CRISPR Populations:

Thirty-six to 48 hours following transfection, single cell suspensions were made using Accutase (STEMCELL Technologies, 07920), re-suspended in 500uL DPBS with Y-27632 dihydrochloride (1mg/mL for iPSCs, 0.5mg/mL for NSCs). Single cells were sorted using 100 micron, low pressure nozzle with a 37 degree chamber in a BD FACSAria II machine run by staff at the Harvard Stem Cell Institute Flow Cytometry Core Facility. GFP+/TO-PRO-3- single cells were sorted into Matrigel-coated 96-well flat bottom plates with Essential 8 media supplemented with 10% CloneR (STEMCELL Technologies, 05889). CloneR supplemented media was changed following manufacturer recommendations. Once reaching 70-80% confluence (approximately 2 weeks post-FACS), cell populations derived from each original single cell per well (clone) were treated with ReLeSR and split into two 96-well plates, one matrigel-coated plate for continued growth along with a matched 96-well plate for PCR genotyping.

#### Genotyping Clonal CRISPR Populations:

DNA from clonal lines was extracted in 96-well format using a Quick-DNA 96 Kit (Zymo Research, D3012). Lines were genotyped for presence of deletion, direct tandem duplication, and wild type alleles at both distal and proximal dual guide cut sites for each respective target locus. Genotyping was conducted using custom designed PCR primers (IDT, **Supplementary Table 2**) with Taq polymerase (NEB, M0273S; Promega, M3001). PCR products were run on 2% Agarose 96-well E-Gels (ThermoFisher, G700802) for visualization. Positive products were cleaned up using ExoSAP-IT (ThermoFisher, 78201.1ML) and submitted for Sanger sequencing using the MGH CCIB DNA Core (**Supplementary Table 3**).

As an orthogonal method of genotyping and for confirmation of clonality, edited lines and a subset of wildtype lines identified by Sanger were also genotyped using ddPCR with DNA copy number probes. Predesigned FAM copy number probes targeting within each CRISPR deletion site were used (ThermoFisher, **Supplementary Table 2**) alongside a VIC reference probe targeting RNase P (ThermoFisher, 4403326) with ddPCR Supermix for Probes, no dUTP (Bio-Rad, 186-3023) following manufacturer instructions. Droplets were generated using a Bio-Rad AutoDG (Biorad, 1864101) and droplets analyzed for fluorescence using the Bio-Rad QX200 (Biorad, 1864003) and QuantaSoft Software.

For inclusion in subsequent differentiation, lines were required to display the edit breakpoint with Sanger sequencing as well as harbor the anticipated allele balance of test to reference material based on ddPCR (1:2 heterozygous deletions, 0:2 homozygous deletions). Wildtype lines for differentiation were selected for a 1:1 allele balance by ddPCR and sequence confirmation of wildtype allele by Sanger. Harboring small InDels at the guide target sites was not used as a criteria for wildtype line exclusion from differentiation given that most of the target sites were in non-coding space. The one exception is in the case of *MEF2C* coding sequence disruption, where wildtype lines were chosen with greater stringency and required completely unedited at the guide target sites given that these fell within coding sequence.
When possible, we required 6 independent clonal lines per genotype at each target locus in the interest of maximizing the number of biological replicates in each experiment. We were unable to generate homozygous deletions for some target regions and in some instances we split single lines in two in order to differentiate 6 lines per genotype and capture technical variation attributable to differentiation as outlined in **Supplementary Table 7**.

#### Selection of TRA-1-60 Positive iPSC CRISPR Lines:

Three - 5 passages following FACS, iPS lines selected for differentiation underwent magnetic activated cell sorting (MACS) for expression of the TRA-1-60 cell surface marker in order to select pluripotent cells for differentiation. Cells were separated using a MiniMACS Separator (Miltenyi Biotec, 130-090-312) with Anti-TRA-1-60 microbeads (Miltenyi Biotec, 130-100-832) following manufacturer instructions (~$2x10^6$ cells per line). TRA-1-60 positive cells were plated with Y-27632 dihydrochloride (1mg/mL), expanded, and cryopreserved using mFreSR. Cells within three passages of TRA-1-60 selection were used for differentiation.

#### Differentiation of Induced Neurons (iNs):

All cell preparations for lentiviral transduction and subsequent iN differentiation were conducted in parallel in a single batch for all lines representing a given locus in order to minimize technical artifacts: 12-18 lines per batch, depending on the locus (6 wildtype, 6 heterozygous deletion, 6 homozygous deletion - for loci where homozygous deletion lines were recovered).

TRA-1-60 positive iPSCs were plated as single cells at 80% confluence on a Matrigel-coated 6-well plate with Y-27632 dihydrochloride (1mg/mL). Polybrene was added at 8 mg/mL 1 hour after re-plating. Cells were incubated with polybrene for 10-15 minutes prior to the addition of lentivirus. Lentiviral constructs for directed differentiation of iPSCs into iNs were made as described previously (Zhang, Neuron 2013). Cells were incubated with lentivirus for 24 hours, followed by a media change with regular E8. Forty-eight to 72 hours following single-cell re-plating, transduced iPSCs were passaged for expansion and subsequent use or cryopreservation in mFreSR.

Transduced iPSCs were expanded onto matrigel-coated T-25 flasks. Once all lines in a batch reached 70-80% confluence, cells were re-plated as single cells onto a new T-25 flask with Neural Maintenance Media (NMM) supplemented with Y-27632 dihydrochloride and 2µg/mL Doxycyline (Clontech, NC0424034) to begin induction of TetO gene driving Ngn2 expression and Puromycin resistance (Day 0). Twenty-four hours after re-plating, media was changed to NMM supplemented with Doxycyline and 1µg/mL Puromycin (Sigma), to begin selection of Ngn2-expressing cells (Day 1). Fresh NMM with Doxycycline and Puromycin was added to cells to continue selection on Days 2 and 3. On Day 4, cells were detached using Accutase and re-plated onto Poly-Ornithine/Laminin (Sigma-Aldrich, P4957/L2020) coated plates with NMM supplemented with 2ug/mL Doxycyline, 10 mg/L human BDNF (Pro-Spec, CYT-207), and 10 mg/L human NT-3 (PeproTech, 450-03). Cells were counted prior to re-plating using a Countess II Automated Cell Counter (Invitrogen, AMQAF1000) with $5x10^5$ cells plated per well of a 12-well plate. Following re-plating, iNs were not exposed to air and required half-media changes every third day. On Day 6, a half media change was conducted with NMM supplemented with Doxycyline, human BDNF, human NT-3, and 2g/L Cytosine β-D-arabinofuranoside (Sigma, C1768-100MG) to prevent glial growth. On Day 8, a half-media change with fresh NMM with Doxycycline, BDNF, and NT-3 was conducted. For subsequent media changes (Day 10+), NMM supplemented with only BDNF and NT-3 was added until cells reached Day 24 of differentiation, at which time differentiation was terminated for RNA extraction, conformation capture crosslinking, or fixation for staining, as described below.

For generation of NSC-derived iNs utilized for the Distal Cluster CRISPR set, we followed the same methods as described above with the following modifications: no MACS selection was conducted, polybrene was omitted (toxic to NSCs), and neural expansion media (NEM) was used in place fo E8 along with supplementation of 0.5mg/mL Y-27632 dihydrochloride during single cell manipulations.

##### Neural Maintenance Media (500mL working stock):

*Modified from Livesey, Nature Protocols 2012*
250mL DMEM/F12 Media (ThermoFisher)
250mL Neurobasal Media (ThermoFisher)
5mL B-27 Supplement (ThermoFisher)
2.5mL N2 Supplement (ThermoFisher)
5mL GlutaMAX (ThermoFisher)
2.5mL NEAA (ThermoFisher)
5mL Penicillin/Streptomycin (ThermoFisher)
125µL Insulin solution from bovine pancreas (Sigma)
250µL Beta-mercaptoethanol (ThermoFisher)
Recipe can be scaled up or down accordingly

#### Differentiation of Neural Stem Cells (NSCs)

All cell preparations for NSC differentiation were conducted with all lines for a given locus handled in the same batch to minimize technical artifacts: 12-18 lines per batch, depending on the locus. TRA-1-60 positive lines were differentiated into NSCs using the protocol detailed by ThermoFisher (MAN0008031). For all target regions, Passage 7 stage NSCs were harvested for RNA extraction, conformation capture crosslinking, or fixation for staining, as described below.

#### RNA Extraction

iNs were harvested on day 24 of differentiation, $5x10^5$ cells per 1mL TRIzol Reagent. NSCs were harvested at the completion of differentiation P7, $2\text{-}3x10^6$ cells per 1mL TRIzol Reagent. Media was aspirated completely from cells with subsequent direct addition of TRIzol Reagent (ThermoFisher, 15596026). Cells were stored at -80C in TRIzol until ready for extraction. Samples in the same differentiation batch underwent TRIzol addition and extraction as a unit. Standard Phenol-Chloroform extraction was used for isolation of RNA using Phasemaker tubes (ThermoFisher, A33249) to ensure extraction purity. Briefly, cells suspended in TRIzol were allowed to reach room temperature and transferred to Phasemaker tubes. 200uL molecular-grade Chloroform (VWR, AAJ67241) was added per 1mL TRIzol reagent. Tubes were capped, vigorously inverted 10x to ensure mixing, and left at room temperature for 10min. Tubes were then spun at 13,000rpm for 15min at 4C. Aqueous phase was removed, transferred to new, LoBind Eppendorf tubes (Fisher Scientific, 13-698-791) with 100uL molecular-grade Isopropanol (Fisher Scientific, 67-63-0) supplemented with 10ug/mL glycogen (Roche, 10901393001), and vigorously inverted 10x. Tubes were left at room temperature for 10min and again spun at 13,000rpm for 15min at 4C. Solution was removed from each tube with care to not disturb the RNA pellet formed. 1mL molecular-grade 70% Ethanol (Fisher Scientific, 64-17-5) was added to each pellet and each pellet dislodged carefully from the side of the tube. RNA pellets in Ethanol were stored at -20C overnight up to one week prior to further processing. TRIzol-extracted RNA from iNs underwent subsequent RNeasy cleanup with on-column digest (Qiagen, 74104, 79254) to ensure removal of residual DNA we often observe carrying over in phenol-chloroform extracted RNA from this cell type. RNA concentration and quality was assessed by Tapestation (Agilent, 5067-5576) with RIN ≥8 required for downstream use for RNAseq or qPCR.

#### Whole Transcriptome Sequencing (RNA-seq) Library Preparation

RNA-seq libraries were prepared by the Genomic and Technology Core (GTC) of Massachusetts General Hospital (MGH) using TruSeq® Stranded mRNA Library Kit (Illumina, San Diego, CA) and prepared according to manufacturer's instructions. In brief, RNA sample quality (based on RNA Integrity Number, or RIN) and quantity was determined based on Agilent 2200 TapeStation (Agilent, Santa Clara, CA) and between 500-100 ng of total RNA was used to prepare libraries. PolyA bead capture was used to enrich for mRNA, followed by stranded reverse transcription and chemical shearing to make appropriate stranded cDNA inserts for library construction. Libraries are finished by adding sample specific, dual barcoded adapters for Illumina sequencing followed by 15 rounds of PCR amplification. Final concentration and size distribution of libraries were evaluated by Agilent 2200 TapeStation (Agilent, Santa Clara, CA) and qPCR, using Library Quantification Kit (KK4854, Kapa Biosystems, Wilmington, MA), and multiplexed by pooling equimolar amounts of each library prior to sequencing. Libraries were sequenced on an Illumina NovaSeq to an average depth of 30M pair-end (PE, 60M total) 150 base-paired (bp) reads.

#### RNAseq data processing

FastQC was used to determine the data quality of the sequencing data. The read pairs were then aligned to the human reference genome (GRCh37, Ensembl release 75, n=47 ) by STAR 2.5.3[1] with following parameters "--outFilterMultimapNmax 1 --outFilterMismatchNoverLmax 0.1 --alignEndsType local". STAR aligner was also used to quantify gene-level read counts based on gene annotations inhuman reference genome (GRCh37, Ensembl release 75). Picard (http://broadinstitute.github.io/picard/) and RNAseqQC [2] were used for quality control of the data. Samples with less than 20M estimated library size as annotated by Picard were removed. A set of neuronal marker genes were tested to confirm the cellular stage of the cell cultures. Samples with failed QC were removed from subsequent analyses. Counts per million (CPM) were calculated based on the number of uniquely mapped reads. Genes with 0.5 CPM cut-off in 50% of samples in at least one condition in a particular comparison were analyzed in differential expression (DE) and correlation analyses.

#### Differential Gene Expression

Differential expression analysis was performed using R package DESeq2 version 1.16.1[3]. The CPM filtered gene counts were used as input for DESeq2 analysis. DE was performed within each cell and edit type and edited samples (DEL$^{het}$ and DEL$^{hom}$) were compared to corresponding wild type samples. PCA was used to determine the clustering of samples in all the comparisons. To account for unknown sources of variation in the expression data, surrogate variables (SVs) were estimated using Surrogate variable analysis (SVAseq) package with ~ genotype as full model and ~ 1 as reduced model . The estimated SVs were incorporated into the DESeq2 model as ~ genotype + SVs to identify DEGs. To this end, DESeq2 estimated size factors (controlling for differences in the sequencing depth of the samples), the dispersion values for each gene, and fitted a generalized linear model and applied Wald test to calculate differential expression p-value. These steps resulted in estimated log2 fold changes and p values, which were corrected for multiple testing using Benjamini- Hochberg adjusted p-values (FDR). The significant DEGs were selected at FDR < 0.1. Estimated SVs were also used to calculate the SVA corrected counts for co-expression analysis. We computed the statistical significance of overlap of DEGs from two tissues by applying a one-tailed Fisher's exact test. We defined overlapping DEGs in two categories based on DEG's direction of dysregulation, overlapping DEGs dysregulated in the same direction in both tissues. MEF2C was excluded from this comparison.

#### Differential Exon Expression

DEXSeq[4,5] was used to access differentially expressed exons of MEF2C genes. DEXSeq uses HTSeq to count the number of reads mapping to each exon. DEXSeq uses a generalized linear model on the negative binomial distribution and takes biological variation into account to estimate the size factors and dispersions, fit the dispersion-mean relation, and test for differential exon usage. DEXSeq detects differential exon usage with high sensitivity, and gives log2 fold changes against the wild type lines and p-values based on Fisher's exact test. The p-values are corrected following the Benjamini-Hochberg-procedure.

#### Co-expression Analysis

Co-expression network analysis was performed using R package WGCNA[6] for each cell type separately using the signed network type for which we used log transformed CPM filtered and SVA corrected counts. Soft power was selected such that the scale-free topology fit ($R^2$) > 0.8. Smallest module size was set to 50. Merge of the modules with similar eigen gene profiles was performed (Similarity > 70%). Module membership for each gene was re-evaluated based on the module membership p-value; genes with p-value > 0.01 were marked as unassigned (Module 0).

#### Functional enrichment

The enrichment of gene ontologies for DEGs and genes in co-expressed modules was tested using one-tailed Fisher's exact test against the mSig-DB (v7.4) curated lists[7,8] and phenotype-informed literature data including: gene sets and modules previously published with functional associations with neurological phenotype, synaptic activity, and *MEF2C* function. Mouse Model RNAseq DEG Validation[9]; Synaptic genes: Syngo v1.1[10]; *MEF2C* targets: ENCODE LCL ChIPseq[11]; DNA Binding and Transcription Modules[12]; Synaptic Activity Modules[12]; Neuroepithelial Precursor and Neuron Expression Modules[13]. The resulting p-values were corrected for multiple tests using Benjamini-Hochberg correction. The PPI network was generated and annotated for functional association using stringApp[14] in Cytoscape.

#### ATACseq Sample and Library Preparation from WT iNs and NSCs

ATAC-seq was applied to wildtype iPS-derived NSC (n=6) and iN (n=6) clones, with cell lines independently differentiated for each cell type as described above. ATAC-seq was performed as previously described (Buenrostro et al. 2015) with minor modifications. Briefly, 50,000 cells were washed with ice-cold PBS, then re-suspended in 50ul cold Lysis Buffer [10 mM Tris-HCl (pH 7.5), 10 mM NaCl, 3 mM MgCl2, 0.1% (vol/vol) IGEPAL, 0.1% (vol/vol) Tween-20, 0.01% (vol/vol) Digitonin] and kept on ice for 3 minutes. 1 ml Wash

Buffer [10 mM Tris-HCl(pH 7.5), 10 mM NaCl, 3 mM MgCl2, 0.1% (vol/vol) Tween-20] was added and samples were centrifuged for 10 minutes at 4°C and 500g. After lysis, the nuclei were re-suspended in 50 μL transposition reaction mix using reagents from Tagment DNA TDE1 Enzyme and Buffer Kits (Illumina) [2X Tagment DNA Buffer, 0.1% (vol/vol) Tween-20, 0.01% (vol/vol) Digitonin, 2.5 μLl Tn5 Transposase] and incubated for 30 min at 37°C with constant agitation at 1000rpm. The transposed DNA was purified with MinElute Reaction Cleanup kit (Qiagen) and eluted in 20 μL Elution Buffer. DNA fragments were PCR amplified using NEBNext High-Fidelity 2X PCR Master Mix (New England Biolabs) and standard Nextera primers for 15 cycles. Agencourt AMPure XP beads (Beckman Coulter) were used for double size selection using 0.5X and 1.8X bead solution to sample solution (v:v) ratios. The size-selected libraries were run on the TapeStation D1000 tape and reagents (Agilent) in order to determine fragment size profile. Libraries were multiplexed, pooled and sequenced on an Illumina NovaSeq, generating an average depth of 100M PE 150 bp reads.

#### ATACseq Peak Calling from WT iNs and NSCs

ATACseq peak calling followed ENCODE guidance (https://www.encodeproject.org/pipelines/). The paired-end reads were aligned to the human reference genome GRCh38 using Bowtie2 version 2.3.1[15], allowing up to 4 multiple maps. Alignments were first de-duplicated and the coordinates were adjusted according to TN5 transposase cut. Pseudo-replicates are pooled-replicates were generated. Narrow peaks were called by MACS2 version 2.2.7.1[16], with shift size of 75, smooth window size of 150 and p value cutoff of 0.01. Only the top 300,000 peaks were kept for the downstream analyses. IDR package (DOI: 10.1214/11-AOAS466) was then used to cross validate samples and generate conservative and optimal peaks. The genomic coordinates of peak regions were then transferred to the coordinates against GRCh37 using the UCSC tool *LiftOver*. It is worth noting that all peak regions were successfully transferred to GRCh37 with the same peak length.

#### Differentiation Marker Staining

iNs intended for staining were re-plated on Day 4 into 12-well plates onto Poly-L-Ornithine/Laminin-coated coverslips (neuVirto, GG-18-pre) and differentiated to Day 24 following the same protocol outlined above. NSCs intended for staining were passaged so P7 cells were grown on matrigel-coated coverslips (neuVitro, GG-18-pre). Cells were fixed using 4% paraformaldehyde (Sigma-Aldrich, F8775-500ML). Cells were permeablized through incubation in 0.5% Triton X-100, followed by incubation in blocking buffer (2% FCS in 0.1% DPBS Tween-20). Primary antibodies were diluted based on manufacturer suggestion in blocking buffer and incubated on paraflim overnight at 4 degrees. Coverslips were washed 3 times with 0.1% DPBS Tween-20. Secondary antibody was diluted based on manufacturer suggestion in 0.1% DPBS Tween-20 and incubated in dark. Coverslips were washed once with DPBS, followed by addition of Hoechst solution (1μg/mL), and subsequent DPBS wash. Coverslips were mounted on glass slides using Mowiol mounting solution. Images were analyzed using Fiji[17]. All washes and incubations were conducted at room temperature while rocking unless otherwise noted. NSCs were stained for Nestin, Pax6, and Sox1. iNs were stained for Map2, Glut1, and Glut2.

#### Western Blot

Western blot analyses were performed on human lymphoblastoid cells and iPS-derived neural stem cells (NSCs). Recombinant Human MEF2C protein (Abcam, ab114241) was used as positive control and iPSC extracts were used as negative control, since this cell type does not express MEF2C. Recombinant anti-MEF2C rabbit monoclonal antibody (Abcam, ab211493) was used to address MEF2C protein. Since the immunogen used to generate this antibody is C-terminal (from 250 amino acid position towards the C-terminal portion of the protein), the MEF2C protein region corresponding to the antibody's epitope is not encompassed by the CRISPR engineered Type I and Type II *MEF2C* gene deletions. Cells were harvested, pelleted and washed with Dulbecco's phosphate-buffered saline (DPBS) prior to lysis. Nuclear fractions were isolated using NE-PER™ Nuclear and Cytoplasmic Extraction Reagents supplemented with Halt™ Protease Inhibitor Cocktail and the protein concentration of resultant samples was determined using Pierce™ BCA Protein assay kit (all from Thermo Fisher Scientific). Loading samples were prepared from equal amounts of protein (30-65ug) in Bolt™ Sample Reducing Agent and Bolt™ LDS sample buffer (both from Thermo Fisher Scientific). The protein was denatured by heating the mixture at 70°C for 10 minutes, separated by electrophoresis on Bolt™ 4-12% Bis-Tris Plus gels (Thermo Fisher Scientific) and wet-transferred onto Immobilon-P PVDF transfer membranes (Millipore Sigma). The membranes were blocked for 1h at room temperature with Odyssey Blocking buffer in PBS (LI-COR Biosciences) and probed overnight at 4°C in recombinant Anti-MEF2C antibody (Abcam ab211493; 1:300 and 1:150) and Anti-PCNA antibody (Abcam ab29; 1:1000) diluted in Odyssey blocking buffer. After 3 washes in TBS/Tween-20 (Boston BioProducts), the membranes were incubated in IRDye secondary antibodies (LI-COR Bioscience; 1:10,000) diluted in Odyssey blocking buffer at room temperature for 1h. Imaging of the immunolabelled bands was performed using the Odyssey Western Blotting system and quantitatively analyzed using Image Studio™ Lite software (both from LI-COR Biosciences). The MEF2C signal from multiple isoforms (50-52KDa) undetected in iPSC extracts were normalized by the PCNA signal.

#### Multi-Electrode Array (MEA)

48-well MEA plates (Axion Biosystems, M768-tMEA-48B) were prepared by coating wells with 50uL 0.1% Polyethylenimine (Millipore-Sigma, 408727) dissolved in Borate Buffer (Boston Bioproducts, BB-66-500) and sterile filtered with a 0.2µm filter prior to use. Coated wells were incubated at 37C for 1hr. Wells were then washed 6-10x with tissue culture-grade water (Boston Bioproducts, 7732-18-5) to remove residual PEI solution. Once excess water was removed, plates were left under UV light for at least 10min and stored at RT overnight to dry out completely. MEA plates were used within 3 days of coating.

To prepare iNs for electrophysiology readings we largely followed manufacturer protocol with a few modifications. Briefly, TRA1+ iPS lines underwent 1 post-thaw passage prior to further manipulation. All iPS lines for MEA underwent Ngn2 transduction as a single batch to eliminate viral transduction batch effect using the same transduction and selection steps described in “Differentiation if Induced Neurons (iNs)” above. On Day 5 of iN differentiation, cells were detached using Accutase (STEMCELL, 07920), counted using a Countess Cell Counter (ThermoFisher, A27977), and resuspended in NMM supplemented with BDNF, NT-3, Doxycyline and Laminin as outlined in manufacturer protocols (Axion Biosystems: Culturing Human iPSC-derived Excitatory Neurons on Microelectrode Arrays: Maestro Pro MEA) in order to distribute $6x10^4$ cells per well in a volume of 10uL directly to the center of each MEA plate well. Plated cells were left to adhere for 1hr at 37C prior to the addition of 200uL NMM supplemented with BDNF, NT-3, and Doxycyline per well. 200uL NMM supplemented with BDNF, NT-3, Doxycycline, and Ara-C was added on Day 6 for a total volume of 400uL per well. Each well underwent a half media change (removal and addition of 200uL) every 3 days with NMM supplemented with BDNF and NT-3 until Day 24. Doxycycline use was discontinued after Day 10 and Ara-C was only added on Day 6.

Day 27 – Day 60, cells continued to undergo half media change every 3 days but media was changed to BrainPhys (following manufacturer protocol: Axion Biosystems: Culturing Human iPSC-derived Excitatory Neurons on Microelectrode Arrays: Maestro Pro MEA) to promote spiking activity. MEA plate readings were conducted every 3 days, 18-24hr following each media change. Readings were conducted at 37C with 0% $CO_2$ for 15min using a Axion Biosystems Maestro Pro. Manufacturer set thresholds for spike and network burst calling were used.

#### 4C Sample Preparation

We performed UMI-4C according to the protocol by Schwartzman et al[18]. In brief, NSCs (5x10^6 cells per sample) and iNs (0.5-3x10^6 cells per sample) were detached using Accutase (StemCell, 00-4555-56), neutralized with PBS and pelleted, resuspended in a PBS/10% FBS solution and crosslinked using formaldehyde (final concentration 2%) (Sigma-Aldrich, F1635-500ML) for 10 min. The reaction was quenched on ice using glycine (final concentration 0.125M). Cells were then centrifuged, washed and resuspended in cold lysis buffer (50mM Tris-HCl, 150mM NaCl, 5mM EDTA, 0.5% NP-40, 1% Triton X-100 and 1 tablet protease inhibitor per 10mL (cOmplete, Mini, Sigma-Aldrich, 11836153001)) and incubated for 10 min. After centrifugation, pellets were washed, snap-frozen in liquid nitrogen and stored at -80°C until further processing.

Frozen pellets were resuspended in a 500µL pre-diluted DpnII buffer, 15µL pre-heated 10% SDS and incubated on a thermomixer for 1h at 37°C, shaking at 900 RPM. We halved these volumes and those mentioned below for smaller cell pellets (originating from 0.5-1x10^6 cells). After adding 150µL 10% Triton X-100 the solution was incubated again (1h, 37°C, 900 RPM). We digested the chromatin using 600U DpnII (NEB, R0543L) in three stages (200U for 2h, 200U overnight, 200U for 2h) at 37°C and 900 RPM. The solution was incubated at 65°C for 20 min to inactivate the restriction enzyme and put on ice. Next, the chromatin was ligated by adding 4000U of T4 DNA ligase (NEB, M0202M) and 10x T4 DNA ligase buffer to a total volume of 1300µL and incubating the solution overnight at 16°C and 300 RPM. Ligated chromatin was de-crosslinked by incubation with 8µL proteinase K (20mg/mL, Qiagen, 19131) (overnight, 65°C, 300 RPM). A 3C template was then purified using 1x Ampure XP beads (Beckman Coulter, A63881).

#### 4C Library Preparation and Sequencing

Up to 4µg 3C template per sample was sheared using microTUBE snap-cap tubes (Covaris, 520045) in a Covaris M220 sonicator to an average fragment length of 300bp. UMI-4C sequencing libraries were generated using the NEBNext Ultra II library prep kit (NEB, E7645L). For each sample, library prep was performed in four parallel reactions with a maximum input of 1000ng per reaction, including a size selection targeting fragments with a length of 300-400bp (unless input was <100ng) and 4-8 cycles of PCR enrichment depending on the input. Next, two nested PCR reactions were performed to enrich for fragments captured by the viewpoint of interest, both using 2µL 10mM Illumina enrichment primer 2 and either a viewpoint-specific “upstream” (reaction 1, 2µL 10mM) or “downstream” (reaction 2, 2µL 10mM) primer. For each sample, we performed up to 8 nested PCR reactions in parallel with an input of 100-200ng per reaction using the KAPA2G Robust ready mix (Sigma-Aldrich, KK5702) in a volume of 50µL. PCR program: 3 min 95°C, 20 cycles (18 cycles for reaction 2) of 15 sec 95°C, 15 sec 55°C, 60 sec 72°C and final elongation of 5 min 72°C. Between PCR reactions the product was cleaned up using 1x AmpureXP beads and eluted in 21µL. The final PCR product was cleaned up using 0.7x AmpureXP beads and eluted in 25µL. Reactions per sample were pooled and library concentration was quantified via qPCR using the KAPA SYBR FAST qPCR Master Mix (Sigma-Aldrich, KK4602). UMI-4C libraries were pooled and sequenced on an Illumina HiSeq or NovaSeq (paired-end, 2x150 cycles).

#### 4C Data Processing – Delta Map Generation

Reads were mapped to the GRCh37 human reference genome and filtered based on the presence of the downstream primer sequence (20% mismatch allowed). Two primer sequences/viewpoints were used within the *MEF2C* promoter. ViewPoint1 (VP1) was used primarily with homozygous deletion samples. ViewPoint2 (VP2) was designed for reads to capture a phasing SNP identified from whole genome sequencing (WGS) of the GM08330 iPSC background line used for generation of all CRISPR lines used herein. VP2 was used primarily with heterozygous deletion samples to allow for allelic phasing and discrimination of contact differences from the wildtype and deletion alleles present in these samples. For VP1, “GAAAGTCCTTCTAAAGTCGG” and “GATC” were set as the bait sequence and bait pad, respectively. For VP2, sequences “AGTGGCCTCAACATTTAG” and “GCAACGGTCCCAAGGCTTCACTTGAAAGGGGAAGGTTTATTTTATAGGCAAGTACAACTAGATC” were used as the bait sequence and bait pad for the G-allele while sequences “AGTGGCCTCAACATTTAG” and “GCAACTGTCCCAAGGCTTCACTTGAAAGGGGAAGGTTTATTTTATAGGCAAGTACAACTAGATC” were used as the bait sequence and bait pad for the T-allele. For heterozygous samples processed with the VP2 viewpoint primer, fastq files were split based on the presence of this SNP and analyzed separately. The resulting fastq files were used as input to the UMI-4C R package (https://github.com/tanaylab/umi4cpackage) to generate genomic interaction tracks representing UMI counts (i.e. unique interactions) per genomic restriction fragment. The package was then used to generate smoothed, viewpoint-specific interaction profiles for the region of interest. For each profile, interaction counts were normalized to the total UMI count within the profile and the adaptive smoothing parameter (win_cov) was scaled to this statistic as well (with a minimum of 10 and maximum of 180).

#### 4C Data Processing – FDR-Corrected Differential Contact Annotation

Differential contacts were investigated within GRCh37 chr5:86,750,000-90,25,0000. A sliding window with size of 5,000 bp and step of 200bp were applied to this region, which quantified the 4C signals in the sub-regions from each contact track. For *i*-th window (or sub-region) in *j*-th track, the 4C signal $C_{i,j}$ was measured as:

Where $d_k$ represents the read depth at position $k$ of that window. For $j$-th track, the normalization factor $F_j$ was calculated as:

$F_j = S_j / M$

Where  represents the read depth at position chr5:$k$ in $j$-th track;  represents the sum of base-wised depth within chr5:86,750,000-90,25,0000 in $j$-th track and $M$ represents the median value of $\{S_1, S_2, S_3, \ldots S_j\}$. Then the following generalized linear regression:

~ NB(mean $=U_{i,j}$, dispersion = $\alpha_i$)
$\log(U_{i,j} / F_j) \sim G_j$

was fit under negative binomial (NB) distribution using DESeq2 v1.24.0 (PMID: 25516281). For samples derived from View Point 1, $G_j$ represents the genotype of $j$-th track and Wald test was applied to examine whether there was a significant difference between WT samples and each type of CRISPR samples. For samples derived from View Point 2, $G_j$ represents the allele-specific genotype of $j$-th track and Wald test was applied to examine whether there was a significant difference between WT alleles and the alleles with each type of CRISPR manipulation. The subregions with FDR < 0.1 were considered as having significantly differential contacts.

### Supplemental Methods

#### Section 1: *MEF2C* direct disruption molecular QC:

##### Section 1.1: Design of MEF2C deletion iPSC lines:

We generated two types of coding deletions within the gene *MEF2C,* referred to herein as Type I and Type II. Both these deletions were designed to create loss of function (LoF) mutations to assay functional effects of tiered allelic loss of *MEF2C* in human neural cell lines. Both Type I and Type II deletions target the same 5' position, upstream of the canonical promoter of *MEF2C*, but different 3' positions within the gene, resulting in 122kb and 131kb deletions respectively (**Figure 1A**). Both heterozygous and homozygous states of each deletion type were recovered. We intentionally designed all *MEF2C* deletions to exclude the Proximal Boundary position, which is located within an intron of *MEF2C* targeted at GRCh37 chr5:88,025,314-88,025,895. This design consideration was included in order to allow for direct, unconfounded comparison between functional effects of the Proximal Boundary vs *MEF2C* direct deletion. Guide sequences and target positions are listed in **Supplementary Table 1.**

##### Section 1.2: Deletion confirmation in *MEF2C* iPSC lines:

We utilized breakpoint PCR amplification followed by Sanger sequencing in addition to ddPCR to confirm genotypes, clonality, and allelic ratio of each iPSC line. PCR primers and Sanger sequences are included in **Supplementary Table 2.**

In order to use Western blot to quantify MEF2C protein in our cell lines, we first validated our MEF2C antibody using known positive and negative controls. In a control blot with inputs from cytoplasmic protein fraction from lymphoblastoid cell lines (LCLs), nuclear protein fraction from LCLs, nuclear protein fraction from iPSCs, and recombinant MEF2C protein, we confirmed the specificity and sensitivity of our MEF2C antibody, as shown in **Supplementary Figure 1A.** We observed quantifiable MEF2C bands in nuclear fraction LCL lysates and used nuclear protein fraction for all Western Blot inputs. We used protein input extracted from LCLs for our initial antibody validation given known expression of MEF2C in this cell type and technical ease of culturing and extraction.

Relative protein expression was calculated using PCNA as a calibrator. At least two replicate clones per genotype were included for quantification. Two isoform sizes of MEF2C were observed on Western Blot (50/51kDa and 52kDa) summed up as MEF2C protein quantification. Relative protein MEF2C/PCNA ratio per sample was averaged across wildtype samples included in the same blot and used as the denominator in relative ratio calculations per sample per blot. Relative ratios were then compared across blots as outlined in **Supplementary Figure 1**B,C Protein quantification calculations included in **Supplementary Table 3**. Western blots protein quantification matched the expected zygosity and effect of LOF mutation on protein levels ( **Supplementary Figure 1D).**

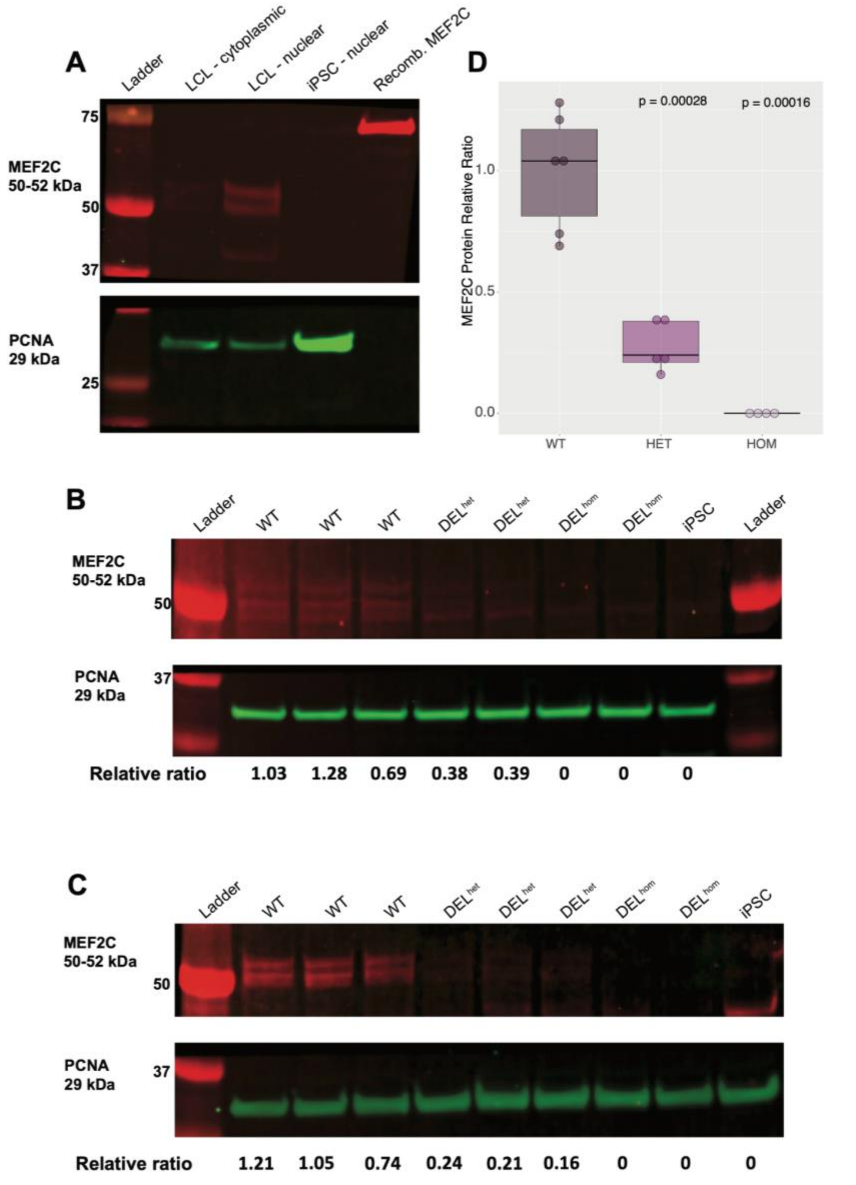

Supplementary Figure 1: MEF2C protein quantification in direct deletion NSCs

**A.** Activity of MEF2C antibody tested against multiple controls. Antibody demonstrated sensitivity against recombinant MEF2C-FLAG reported at 78kDa in size and specificity given lack of band in nuclear fraction of iPSC lysate, a cell type that does not express MEF2C. Cytoplasmic and nuclear fractions of LCLs tested with nuclear fraction utilized for subsequent experimental blots. The band from recombinant MEF2C protein is observed higher than MEF2C observed in LCLs given that the recombinant protein is attached to a FLAG tag. **B.** MEF2C protein abundance quantified from Type I deletion NSCs with direct heterozygous and homozygous deletion of MEF2C relative to matched controls. Clones represented in Type I deletion blot include replicates of J (WT), K (DELhet), and H (DELhom). **C.** MEF2C protein abundance quantified from Type II deletion NSCs with direct heterozygous and homozygous deletion of MEF2C relative to matched controls. Clones represented in Type II deletion blot include M (WT), N (WT), R (DELhet), and X (DELhom). For both Type I and Type II blots at least two technical replicates of each deletion type were included for quantification. Total MEF2C (50/51 + 52kDa bands) was used in MEF2C quantification relative to PCNA (29kDa band) for all. iPSC lysate included as negative control given lack of MEF2C expression. **D.** Boxplot of relative MEF2C protein expression per genotype, including both Type I and Type II deletions. P-values calculated using t-test relative to wildtype mean. Despite the observed increase in mRNA expression for exons outside the deletion, we observe the reduction or total loss of overall MEF2C expression at the protein level in heterozygous and homozygous deletion lines respectively in NSCs (Supplementary Figure 2). For both Type I and Type II deletions, homozygous deletion yielded no detectable MEF2C protein while heterozygous deletion resulted in ~60-85% expression reduction. It's important to note that the gene region that encodes the antibody's epitope is not encompassed by the CRISPR deletion on the MEF2C gene (**see antibody information in Material and Methods**).

#### Section 1.3: Confirmation of MEF2C expression reduction in direct disruption CRISPR edits:

In order to confirm our direct *MEF2C* deletion CRISPR models accurately represented the expected heterozygous (50%) and homozygous (0%) expression levels of *MEF2C* relative to wildtype, we evaluated the mRNA expression of *MEF2C* in NSCs and iNs using RNAseq as well as characterized MEF2C protein expression in NSCs using Western blot.

We also determined exon usage per genotype in iNs and NSCs (**Supplementary Figure 1**, details in **Materials and Methods**). We used the reads mapping to *MEF2C* exons, to estimate the exon expression in $DEL^{het}$, $DEL^{hom}$ and WT samples using R DEXSeq package DEXSeqin R[4,5].

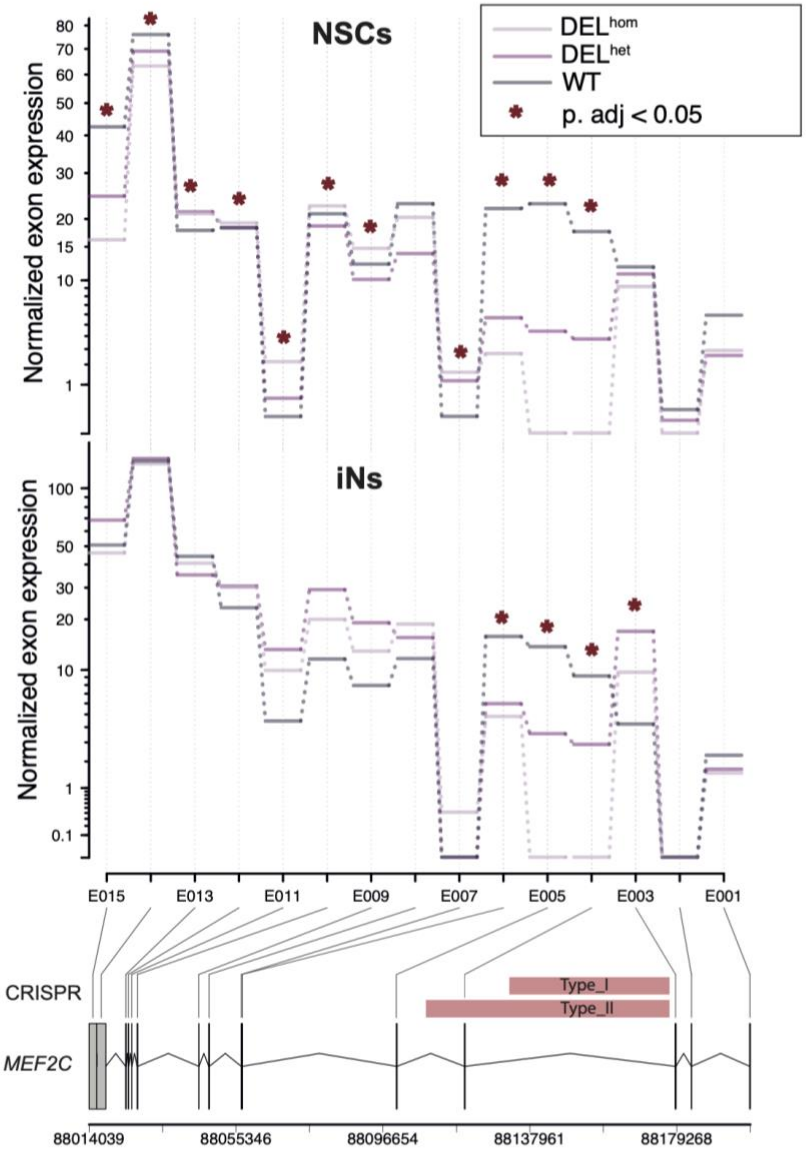

Supplementary Figure 2: MEF2C exon usage in direct deletion CRISPR lines

Exon usage of CRISPR vs wildtype NSCs and iNs was calculated using DEXseq[5]. Exons edited out by Type I or Type II *MEF2C* deletion indicated by horizontal bars (bars correspond to the scale of the gene model at the bottom of the figure). Exons with significantly differential expression in either $DEL^{het}$ or $DEL^{hom}$ vs wildtype demarcated by asterisks (FDR<0.05). In NSCs, we observed a significant down regulation of exons (E004-E007), which were contained within the deleted region (FDR<0.05) but observed an over-expression of exons from E009 to E015, upstream of the deleted region. For iNs, we observed down-regulation of (E003-E005), confirming down-regulation of exons within deleted gDNA. The expression of exons located within each deletion type was consistent with the allelic ratio, but increased expression was observed for exons outside of the deleted region. In contrast to the other three types of deletions, the homozygous Type II deletion demonstrated reduced expression of all exons relative to wildtype, regardless of position relative to deletion. Interestingly, expression of later exons rose to >2x that of wildtype expression for exons 3' to the site of deletion (FDR < 0.05), with NSCs displaying more marked expression increases than iNs of matched CRISPR lines. We again interpreted this variable expression as erroneous transcription that would not generate canonical MEF2C protein, which we confirmed using Western blot.

#### Section 1.4: Differentiation marker expression validation in *MEF2C* iNs and NSCs:

We confirmed our iPS-derived neural differentiations by demonstrating the expression of known marker genes NSCs and iNs both at the RNA level from RNAseq (**Supplementary Figure 3**) as well as at the protein level with immunocytochemical staining (**Supplementary Figure 4**).

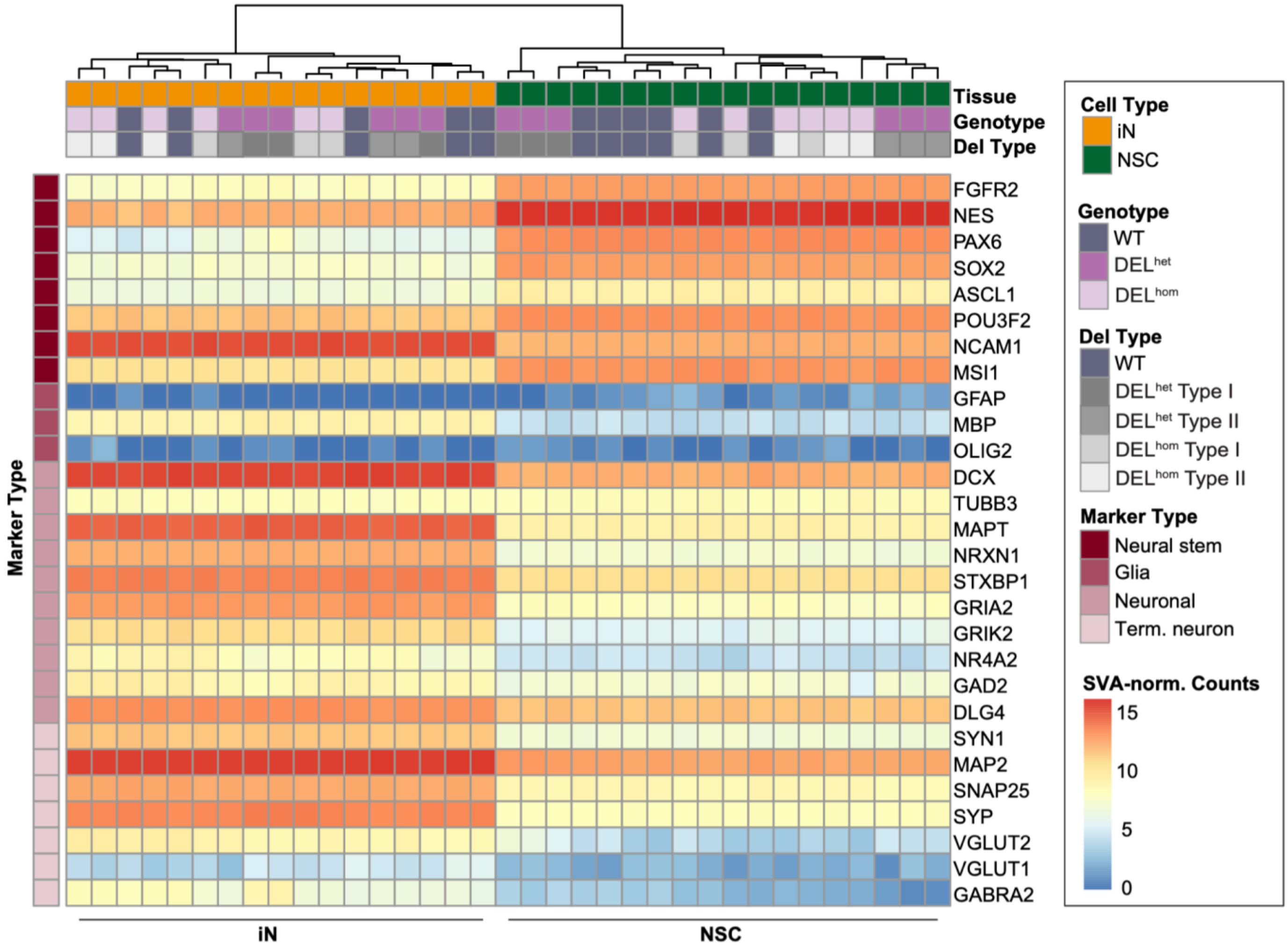

Supplementary Figure 3: Neuronal differentiation marker expression in MEF2C direct disruption iNs and NSCs as ascertained by RNAseq.

Expression of known neuronal marker genes curated from the literature based on SVA-normalized counts from each replicate iN and NSC sample from the direct *MEF2C* disruption CRISPR set. “Term. neuron” represents markers observed in terminally differentiated neurons.

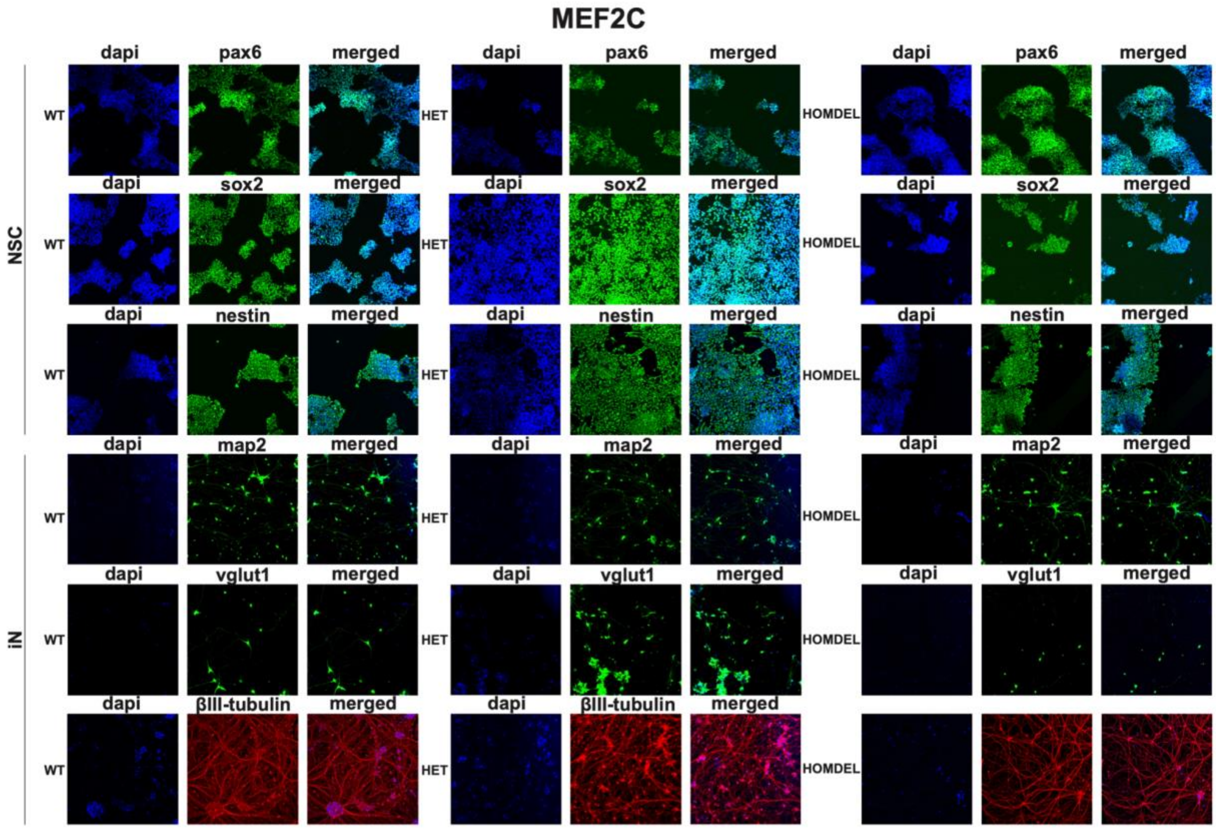

Supplementary Figure 4: Differentiation marker expression in MEF2C direct disruption iNs and NSCs as ascertained by immunocytochemical protein staining.
Marker proteins SOX2, NES (Nestin), and PAX6 stained alongside DAPI in NSCs. Marker genes MAP2, BetaIII-tubulin, and VGLUT1 are stained alongside DAPI in iNs.

#### Section 2: *MEF2C* direct disruption multi-electrode array (MEA):

##### Section 2.1: Analysis of *MEF2C* direct disruption iNs for multi-electrode array (MEA):

Two 48-well MEA plates were included in the analysis reflected in **Figure 3**, referred to as MEF2C MEA Plate 1 and MEF2C MEA Plate 2. The 48 total wells on Plate 1 contained 20 wells of wildtype cells (10 wells clone J, 10 wells clone N), 18 wells of heterozygous cells (9 wells clone K, 9 wells clone R), and 10 wells of homozygous cells (10 wells clone X). The 48 total wells on Plate 2 contained 20 wells of wildtype cells (10 wells clone N, 10 wells clone M), 18 wells of heterozygous cells (9 wells clone K, 9 wells clone R), and 10 wells of homozygous cells (10 wells clone X). Replicate wells of clone N were included on both Plates 1 and 2 as a wildtype comparator set. Considering wells from both Plates 1 and 2 we plated a total of 40 replicate wildtype wells, 36 replicate heterozygous deletion wells, and 20 replicate homozygous deletion wells. Well filtering was conducted to retain wells with at least 50% active electrodes (requiring 8 of 16 total electrodes per well passing activity threshold, as defined by manufacturer) for downstream analyses. For MEF2C MEA Plate 1 and 2 respectively, 44/48 and 47/48 wells passed electrode activity filtering for at least one day of measurements. Differentiation day reflecting measurements of focus for downstream analyses was selected based on date with highest average post-filtering weighted mean firing rate for wells containing wildtype cells. This date was Day 55 for Plate 1 (average weighted mean firing rate for WTs 3.34Hz) and Day 51 for Plate 2 (average weighted mean firing rate for WTs 2.96Hz)

(**Supplementary Figure 5-6**). Combined analysis of MEF2C MEA Plates 1 and 2 was conducted by normalizing datapoints per well by average of plate-matched wildtypes. P-values calculated using paired t-test comparing mean value of each measure vs matched wildtype.

We also include the results of our MEA pilot experiment to demonstrate replication of the same reported synaptic trends in a smaller but independent experiment (**Supplementary Figure 7**). The MEF2C MEA Pilot Plate contained a total of 12 wildtype replicate wells (6 wells clone J, 6 wells clone N), a total of 12 heterozygous deletion wells (6 wells clone K, 6 wells clone R), and a total of 6 homozygous deletions wells (6 wells clone X). Well filtering was conducted to retain wells with at least 50% active electrodes (requiring 8 of 16 total electrodes per well passing activity threshold, as defined by manufacturer) for downstream analyses. Differentiation day reflecting measurements of focus for downstream analyses was selected based on the date with the highest average post-filtering weighted mean firing rate for wells containing wildtype cells. Day 41 average weighted mean firing rate = 2.10Hz. For MEF2C MEA Pilot Plate, 12/30 wells passed electrode activity filtering on differentiation Day 41. We exclude the MEF2C MEA Pilot results from our primary analysis given the lower number of well replicates but include as a Supplementary secondary validation given the clear demonstrated reproducibility of resultant synaptic effects reported.

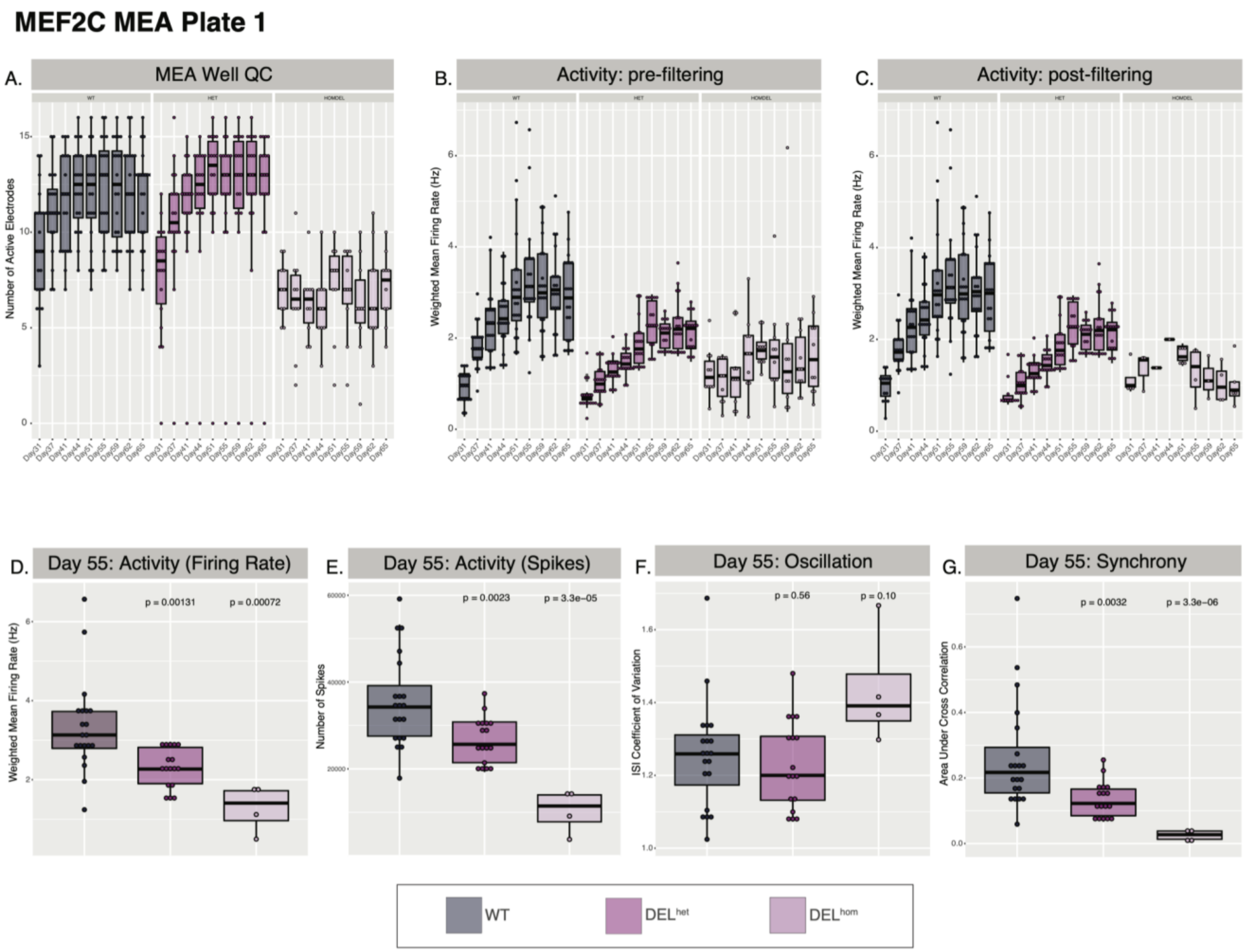

Supplementary Figure 5: MEF2C MEA Plate 1.

**A.** Well QC demonstrating number of active electrodes per genotype per MEA reading day. Weighted mean firing rate per genotype per MEA reading day shown before **B.** and after **C.** filtering for wells with at least 8 active electrodes. **D-G.** measurements per genotype on Day 55, focus day given highest wildtype firing.

#### MEF2C MEA Plate 2

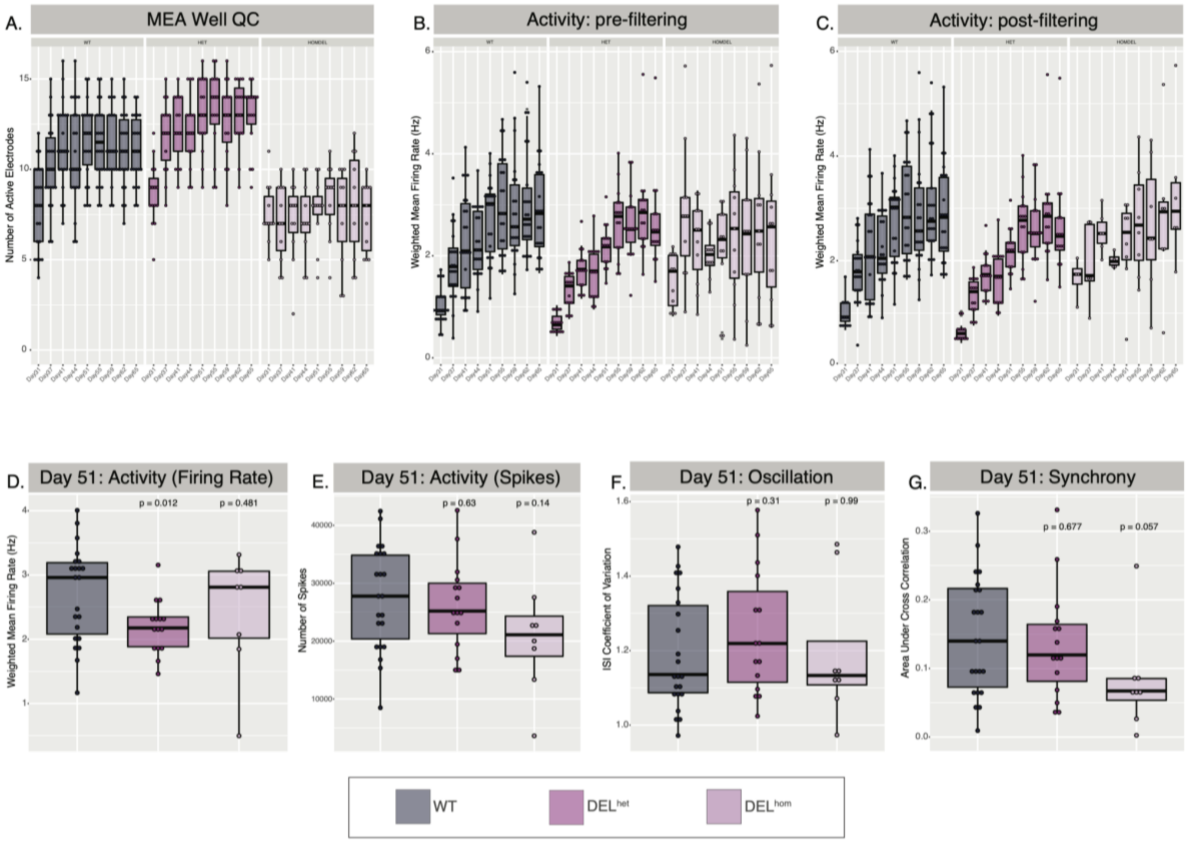

Supplementary Figure 6: MEF2C MEA Plate 2

**A.** Well QC demonstrating number of active electrodes per genotype per MEA reading day. Weighted mean firing rate per genotype per MEA reading day shown before **B.** and after **C.** filtering for wells with at least 8 active electrodes. **D-G.** measurements per genotype on Day 55, focus day given highest wildtype firing.

#### MEF2C MEA Pilot Plate

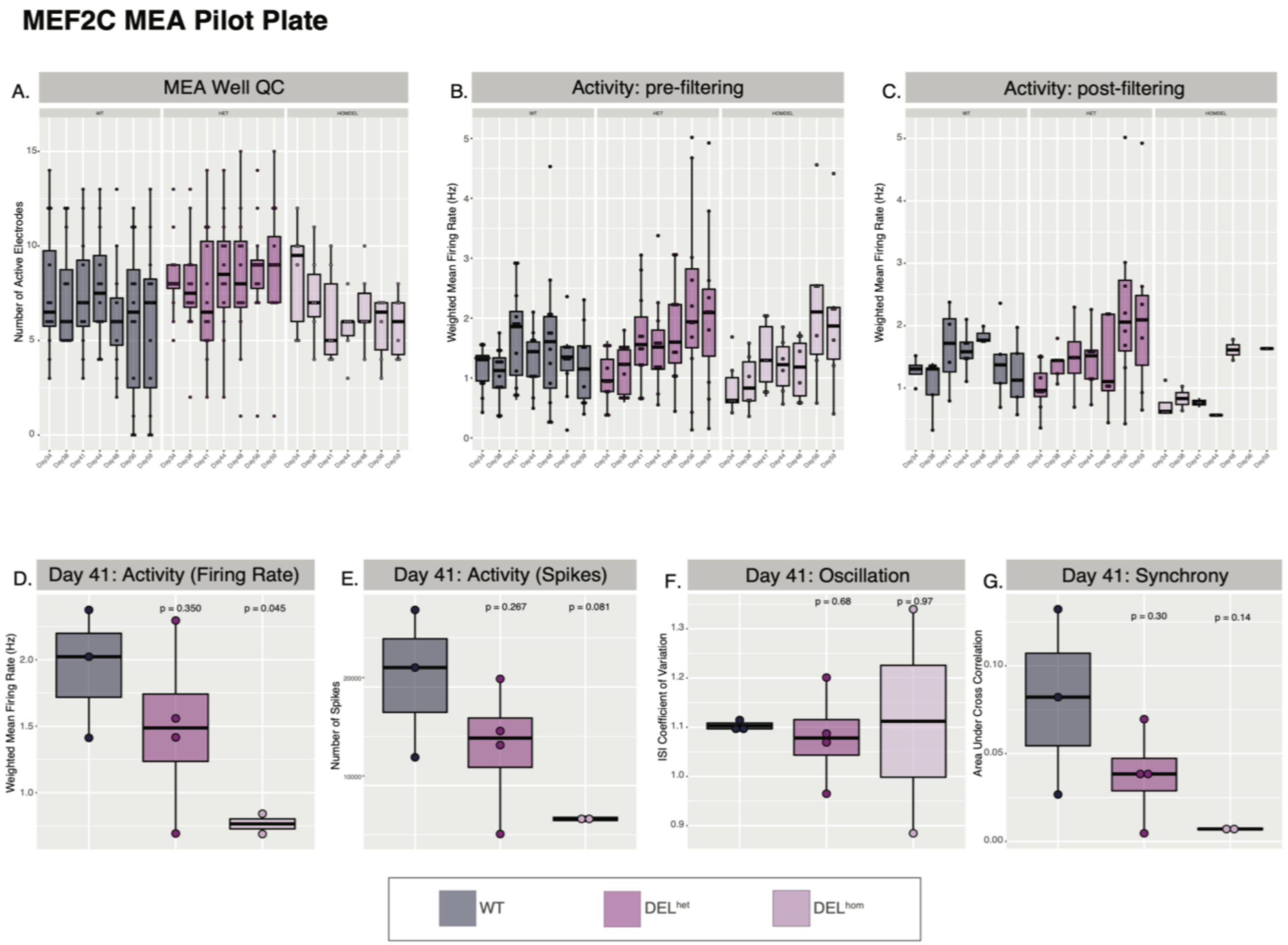

Supplementary Figure 7: MEF2C MEA Pilot Plate

**A.** Well QC demonstrating number of active electrodes per genotype per MEA reading day. Weighted mean firing rate per genotype per MEA reading day shown before **B.** and after **C.** filtering for wells with at least 8 active electrodes. **D-G.** measurements per genotype on Day 55, focus day given highest wildtype firing.

#### Section 3: Characterization of 5q14.3 positional effect CRISPR targets

##### Section 3.1: Determination of Proximal and Distal Boundary targets for CRISPR

In light of the population genetic evidence pointing to 5q14.3 as a strong positional effect locus, we sought multiple lines of functional genomic evidence to determine the 2D and 3D regulatory elements to target for CRISPR dissection. We hypothesized the 5q14.3 BCA case rearrangements indirectly caused *MEF2C* haploinsufficiency via disruption of distal regulatory elements responsible for proper *MEF2C* expression. Given this, we moved to identify regulatory elements for dissection most reasonably capable of driving broadly acting haploinsufficiency based on both experimentally validated interaction with *MEF2C* and annotated tissue/cell type usage. More specifically, we aimed to primarily consider elements with evidence of regulating *MEF2C* that were active in the brain but also largely cell/tissue type invariant. We deemed these patterns of activity to be phenotype-relevant while also having the highest likelihood of causing systemic haploinsufficiency rather than cell type-specific dysregulation. While we recognize that cell type specific disruption could most certainly cause haploinsufficiency-driven pathogenesis, we first prioritized the lower hanging fruit of broad disruption prior to the far more complex cell type specific interrogations.

In consideration of 2D regulatory elements, we first turned to functionally validated enhancers annotations defined previously in D'haene et al, 2018[19]. Briefly, enhancers of *MEF2C* were initially identified using 4Cseq in SH-SY5Y neuroblastoma cell line and validated in zebrafish using a luciferase reporter assay. A total of 16 distal enhancers (e1-e16) were identified based on both long-range interaction *in vitro* and capability of enhancer activity *in vivo.*

For consideration of 3D regulatory elements, we were initially guided by HiC maps and ChIPseq annotations of 3D structural proteins (CTCF and SMC3) from GM12878 LCLs[11,20]. We started with LCLs given that our starting material for the putatively 5q14.3 TAD-disrupting BCA cases was from LCLs as well. Through the integration of HiC, motif sequence mapping, structural protein ChIPseq data, we identified the sites we refer to herein as the Proximal and Distal Boundary elements. We first narrowed our search space by using TAD and loop definitions from the literature[21,22]. We then sought to refine the 3D boundary definitions for CRISPR perturbation by identifying the CTCF binding site positions based on motif sequence directionality as well as sites of CTCF occupancy in LCLs using HiGlass.io[23], as visually demonstrated in **Figure 4B.** We identified the directly oriented CTCF binding site positions within the annotated *MEF2C*-containing loop supported by both CTCF and SMC3 occupancy based on LCL ChIPseq, considering the importance of CTCF directionality based on previous reports[22]. We furthermore observed the utilization of the Proximal and Distal Boundary elements in a diversity of cell types as demonstrated by CTCF ChIPseq in **Supplementary Figure 8.** Of note, our 3D Distal Boundary element annotation encompasses enhancer e13.

We first sought evidence in support of the identified 2D and 3D *MEF2C*-associated regulatory elements being utilized in the human brain. To this end, we considered open chromatin regions (OCRs) identified by ATACseq in multiple brain-related cell and tissue types defined by the Brain Open Chromatin Atlas (BOCA:https://bendlj01.u.hpc.mssm.edu/multireg/)[24] and psychENCODE[25]. We utilized neuronal (NueN+) and non-neuronal (NeuN-) cell OCR calls from 14 brain sections included in BOCA. In addition, we considered neuronal (NueN+) and non-neuronal (NueN-) cell OCR calls from 2 brain regions (Broadmann's area 9 and Nucleus accumbens). A total of 32 NueN+ and NueN- brain tissue inputs were considered across both studies. We observe direct (minimum 1bp) OCR overlap for 31/32 total inputs with the Proximal Boundary and 26/32 inputs for the Distal Boundary. Enhancers e8 and e15 represent the non-coding enhancers with the highest overlap with the considered ATAC OCRs, both overlapping OCRs identified in 15/32 tissue inputs. These observations highlight the 3D boundary elements as consistent sites of open chromatin, suggesting their consistent utilization in cell types observed in the brain. Notably, enhancer e16 is observed as an OCR in 32/32 ATAC inputs but given that this enhancer overlaps the promoter of the broadly expressed *CETN3* gene, we cannot confirm whether this region is a OCR due to transcription rather than distal regulatory activity from ATACseq alone (**Supplementary Figure 9**).

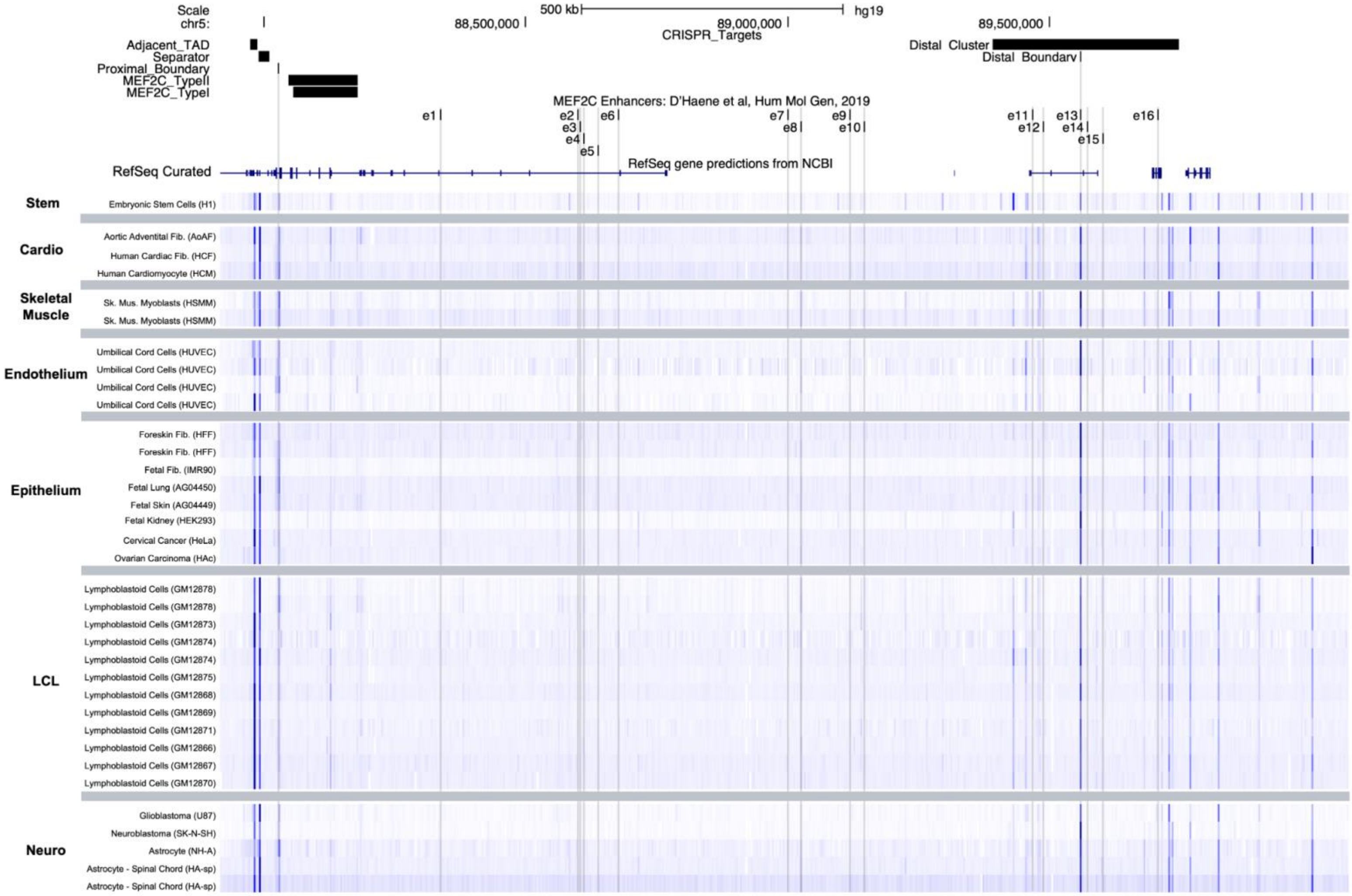

Supplementary Figure 8: CTCF ChIPseq from ENCODE cell and tissue types.

ChIPseq tracks annotating CTCF binding sites. Tracks configured to fixed heatmap scale minimum of 0 and maximum of 100 using WashU Epigenome Browser[26].

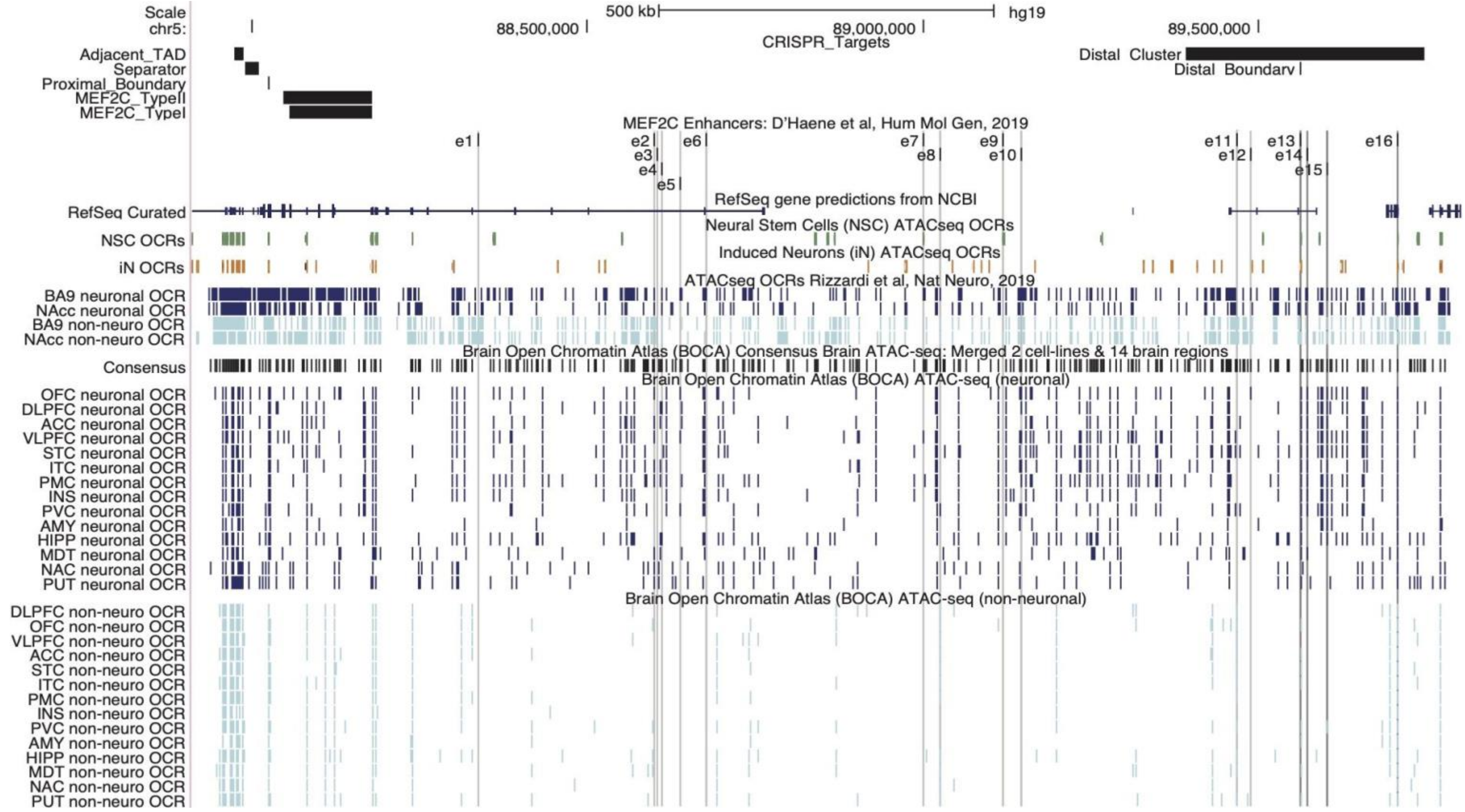

Supplementary Figure 9 : ATACseq open chromatin regions in neural tissue and cell types.

Open chromatin regions (OCRs) called from ATACseq in neural tissue from psychENCODE[25] and BOCA[24] shown alongside OCRs called from ATACseq in wildtype iNs and NSCs. In 34 considered brain-related cell and tissue types, we observe most consistent overlap between annotated *MEF2C* enhancers and e13-e16, which correspond with both the coordinates of the Distal Boundary and Distal Cluster CRISPR targets.

In addition to publicly available ATACseq OCRs, we also compared our identified *MEF2C*-related 2D and 3D regulatory elements of interest with OCRs in our wildtype iN and NSC neural models (1bp minimum overlap). We considered these comparisons of important technical relevance to determine the regulatory elements we would be able to reliably assay in our *in vitro* models systems. Both the Proximal and Distal Boundary positions directly overlap OCRs from iN and NSC ATACseq. Of the 16 functionally validated enhancers, only 2, e13 and e16, overlap with OCRs annotated in both iNs and NSCs, with e7 and e9 overlapping only NSC OCRs (**Supplementary Figure 9**). As mentioned previously, e13 directly overlaps the Distal Boundary and e16 overlaps the promoter of *CETN3*, making observations from the latter potentially the result of active transcription rather than distal regulatory usage.

We further interrogated the *MEF2C* regulation potential of the 2D and 3D regulatory elements of interest in our wildtype iNs and NSCs by assaying their long-range contact frequency with the *MEF2C* promoter using UMI-4C. We observed the Proximal and Distal Boundary elements emerging as two of the strongest long-range contacts with the *MEF2C* promoter in both iNs and NSCs. These results continued to underscore the 3D regulatory elements as being both of interest due to human brain usage and technically accessible for CRISPR dissection in our *in vitro* models (**Supplementary Figure 10**).

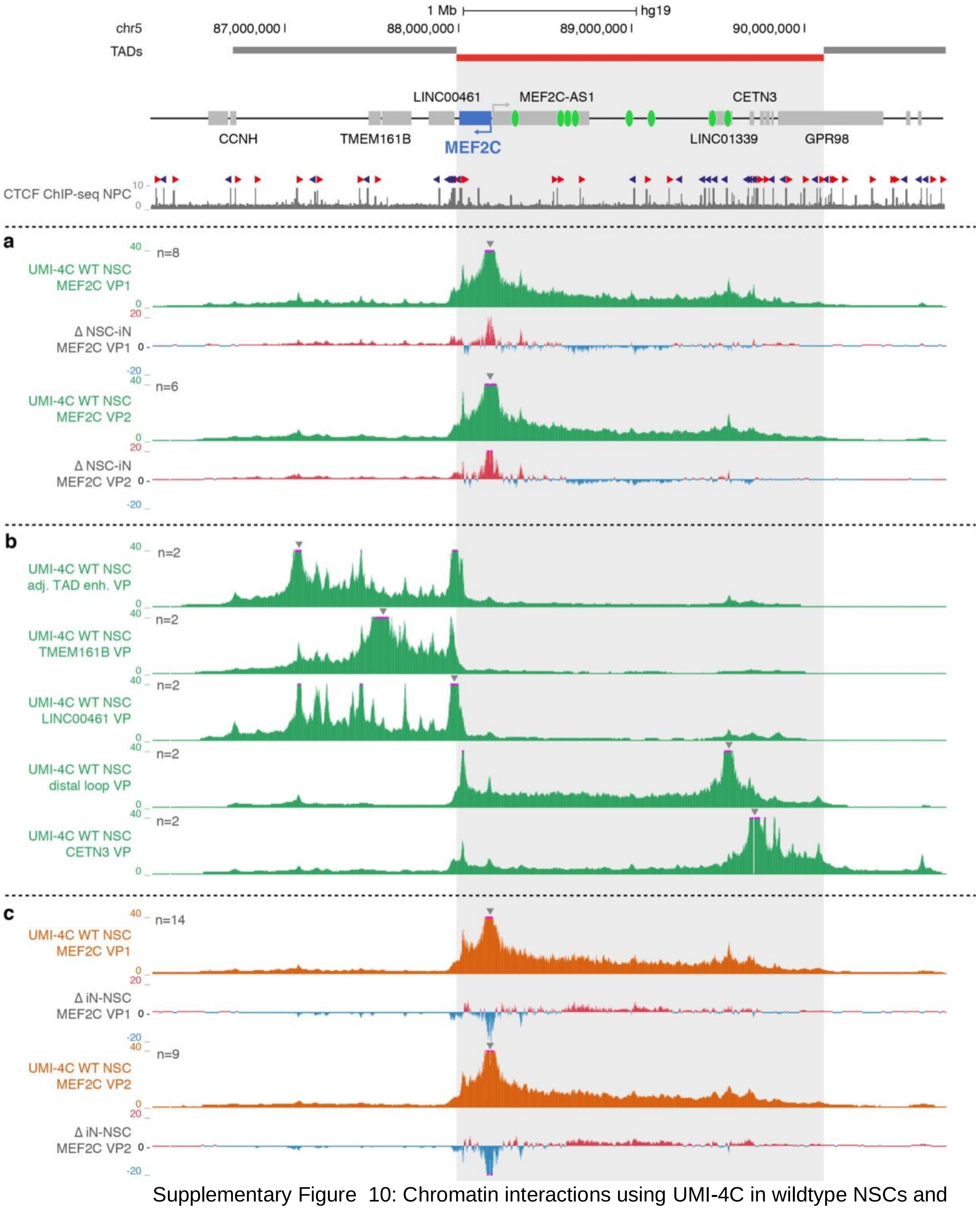

Supplementary Figure 10: Chromatin interactions using UMI-4C in wildtype NSCs and iNs

Normalized and smoothed UMI-4C interaction counts and subtraction (∆) profiles. UMI-4C viewpoints (VPs) are indicated by arrows, n represents the number of biological replicates. MEF2C locus with topologically associating domain (TAD) structure, annotated enhancer elements (green ovals) and CTCF binding in neural progenitor cells (NPCs, available from ENCODE) on top. (a) Wildtype (WT) neural stem cells (NSCs), MEF2C VP 1 and 2. (b) WT NSCs, alternative VPs. (c) WT induced neurons (iNs), MEF2C VP 1 and 2.

Having identified consistent neuro-usage of our 3D boundary elements alongside moderate usage of a few *MEF2C* enhancers, we next aimed to consider more refined regulatory annotations from ChIPseq in a broad array of cell and tissue types. The goal with this layer of consideration was to identify which regulatory elements displayed broad cell/tissue type usage while better understanding the biological nature of their activity beyond simply openness of chromatin. Specifically, we considered H3K27Ac, H3K4me1, and CTCF annotations from publicly available ChIPseq peaks generated as part of the ENCODE project in a range of cell and tissue types. We considered peaks from 107, 105, and 133 cell and tissue type samples for H3K27Ac, H3K4me1, and CTCF ChIPseq respectively. We observed e1-e16 displaying highly variable and limited corroboration by the enhancer element ChIPseq marks (H3K27Ac and H3K4me1) in different cell and tissue types. In contrast, the positions of the 3D boundary elements, the Proximal and Distal Boundaries, emerged largely cell type invariable based on CTCF ChIPseq, with 129/133 and 133/133 considered samples displaying annotated peaks overlapping each boundary respectively. These findings highlighted the 3D regulatory elements as targets of interest based on their broad cell type usage. The publicly available data discussed above is interactively viewable in the UCSC session below: http://genome.ucsc.edu/s/kianamohajeri/5q14.3_ATAC_ChIP.

Through the synthesis of these analyses, we observe that not only do the Proximal and Distal Boundary emerge as positions of consistently open chromatin and CTCF binding across brain and non-brain cell and tissue types, but they are also two of the most prominent long-range contacts in our iNs and NSCs, highlighting these sites as cell type invariant 3D functional elements with evidence of *MEF2C* regulation (**Figure 4** and **Supplementary Figure 10**). Not surprisingly, the functionally validated enhancers e1-e16 are variably corroborated by brain and non-brain cell and tissue types, underscoring that while these enhancers have demonstrated strong evidence of regulating *MEF2C*, their roles and essentiality vary from cell type to cell type.

Based on these findings, we sought to interrogate the overarching and consistent 3D functional element landscape responsible for orchestrating gene-enhancer interactions in lieu of a dissection of individual enhancer elements. To this end, we targeted the Proximal and Distal Boundary elements as our main experimental sites for CRISPR dissection. Given nuanced considerations of cell type specificity individual enhancers require, we saw CRISPR disruption of the 3D regulatory elements as a preliminary opportunity to address questions of distal regulation of *MEF2C*, which would inform future work on individual enhancer contributions.

#### Section 3.2: Design of five positional effect CRISPR target positions within 5q14.3 locus

As outlined above, we targeted the Proximal Boundary and Distal Boundary positions for deletion, which represent the primary experimental lines in this study. Furthermore, we also designed CRISPR deletions at 2 additional positions within the 5q14.3 locus at the outset of our experiment which included a negative control: disruption of a non-*MEF2C* containing TAD boundary (Adjacent TAD) and a region devoid of CTCF binding sites (Separator). We expanded upon the initial 4 positional effect targets with the additional Distal Cluster CRISPR deletion, aimed at determining the potential contribution of CTCF buffering in the prevention of *MEF2C* dysregulation by deletion of the Distal Boundary. Guide sequences and target positions are listed in **Supplementary Table 1 (Figure 4A).**

Adjacent TAD

We designed the deletion of the site we termed Adjacent TAD to demonstrate the effect of a 3D boundary element within 5q14.3 that we did not expect to have an effect on *MEF2C* expression. The Adjacent TAD is a deletion of the distal boundary of the TAD upstream and adjacent to the *MEF2C*-containing TAD. This 13kb deletion does not overlap any genes but does remove the 5’ end of *LINC00461*. Given that the Adjacent TAD deletion targeted the boundary of a domain not inclusive of *MEF2C*, we expected this deletion to have no effect on *MEF2C* expression. This deletion encompasses 2 occupied CTCF binding sites based on ChIPseq in GM12878 LCLs as demonstrated in **Figure 4E.**

Separator
The Separator region is a non-coding site between the Adjacent TAD and *MEF2C*-containing TAD, representing a position within 5q14.3 devoid of occupied CTCF binding sites. Given the lack of annotated 3D functional elements within this segment, we did not expect deletion of this position to have any effect on *MEF2C* expression and used this target as a negative control to demonstrate a large (21kb) deletion of gDNA within 5q14.3 alone was insufficient dysregulate *MEF2C* as demonstrated in **Figure 4E**.

Proximal Boundary
As outlined above, we sought to target the proximal end of the loop harboring *MEF2C* as one of two primary CRISPR targets focused on determining whether disruption of 3D structure was sufficient to dysregulate *MEF2C*. The Proximal Boundary CRISPR targeted a single occupied CTCF binding site within a 3' intron of *MEF2C*. The 581bp target site sits 181bp and 132bp away from the immediately upstream and downstream exons, respectively. This deletion was not expected to disrupt nearby splice sites based on in silico prediction. Three occupied CTCF binding sites are actually present within two neighboring introns of *MEF2C*. We chose to target the single position with the highest level of occupancy based on CTCF ChIPseq in GM12878 LCLs in lieu of removing all 3 occupied sites, along with the separating exon, with a single deletion. The singular site deletion did not disrupt coding sequence, creating an appropriate model for assaying non-coding effects on *MEF2C* expression as demonstrated in **Figure 4E**.

Distal Boundary
As outlined above, we sought to target the distal end of the loop harboring *MEF2C* as one of two primary CRISPR targets focused on determining whether disruption of 3D structure was sufficient to dysregulate *MEF2C*. The Distal Boundary CRISPR represents a 3.3kb deletion targeting a single occupied CTCF binding site >1.3Mb distal to the *MEF2C* promoter. This deletion also partially overlaps *LINC01339* as shown in **Figure 4E**.

Distal Cluster
This target represented a follow-up CRISPR set which was generated after our observations of the Distal Boundary disruption yielding a lack of robust differential *MEF2C* down-regulation. The goal of this deletion was to remove not only the distal end of the 3D loop containing *MEF2C*, but to also eliminate the properly (reverse) oriented but unoccupied CTCF binding sites. We hypothesized these nearby CTCF binding sites were being utilized in the absence of the canonical, normally occupied binding site within the Distal Boundary site, allowing the 3D loop structure to still form and buffering against *MEF2C* dysregulation. To test this, we designed a 354kb deletion. This deletion was designed to address the aforementioned questions related to 3D regulatory disruption but we also note that 6 enhancers (e11-e16) are encompassed by this deletion and in fact directly overlap the positions of the CTCF binding sites. Given this inherent intertwining of 2D and 3D functional elements within the 5q14.3 locus, we highlight that both types of effects must be considered in explaining functional effects of the Distal Cluster deletion. This deletion is also directly disruptive to *CETN3, MIR3660,* and *LINC01339* as shown in **Figure 4E**.

#### Section 3.3: Genotype characterization of 5q14.3 iPSC CRISPR lines

In order to confirm precise editing and homogeneity of each cellular model generated, we used orthogonal genotyping methods of PCR, ddPCR, and Sanger sequencing at the iPS stage prior to carrying lines forward for pluripotency selection, differentiation, and functional assays.

Unique PCR assays were designed for each target to capture wildtype, deletion, and direct tandem duplication alleles generated by CRISPR. Each PCR assay was first tested in gDNA extracted from bulk transfected iPSCs to confirm efficiency and accuracy of the genotyping assay as well as guide efficiency. Following PCR and Sanger screening, we used the secondary measure of ddPCR to further confirm genotype as well as ensure lines were clonal and without evidence of mosaicism from allele balance. PCR primer and ddPCR probe sequences are listed in **Supplementary Table 2.** Droplet counts and allelic ratios obtained per edited iPSC clone are displayed in **Supplementary Figures 11-12.** Genotyping data from ddPCR alongside breakpoint and wildtype allele Sanger Sequence are also included in **Supplementary Table 4** with editing efficiencies per target site displayed in **Supplementary Figure 13**.

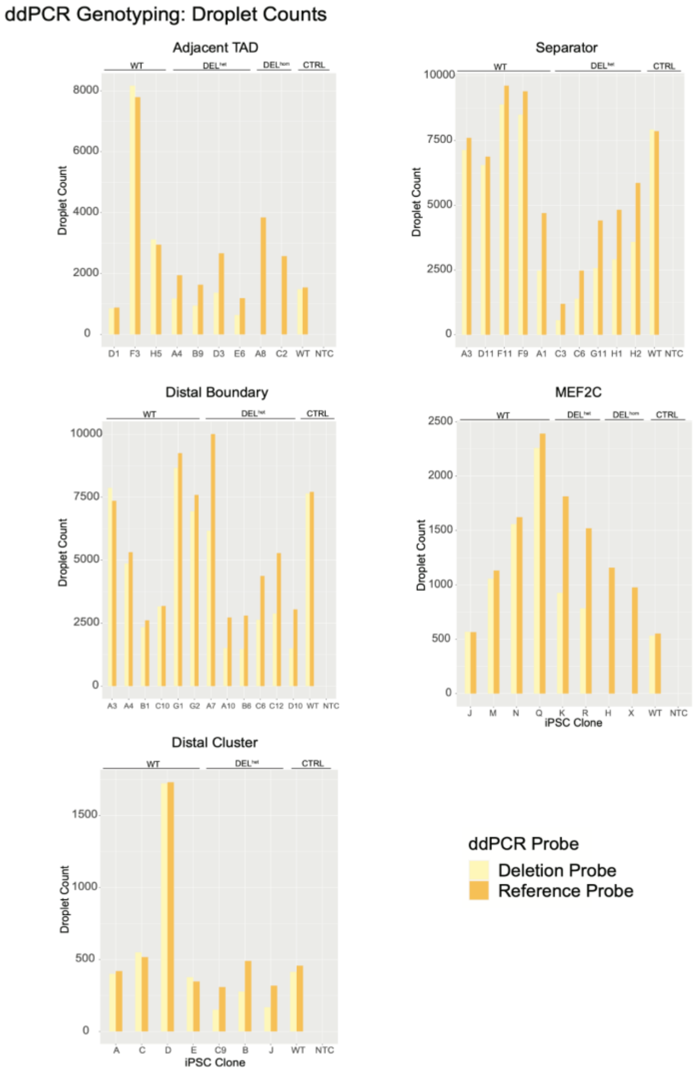

Supplementary Figure 11 : ddPCR droplet counts from reference and deletion probes.

gDNA per iPSC clone underwent ddPCR. Relative number of droplets positive for gDNA from sequence overlapping deletion vs sequence of RNaseP used for determination of allelic ratios.

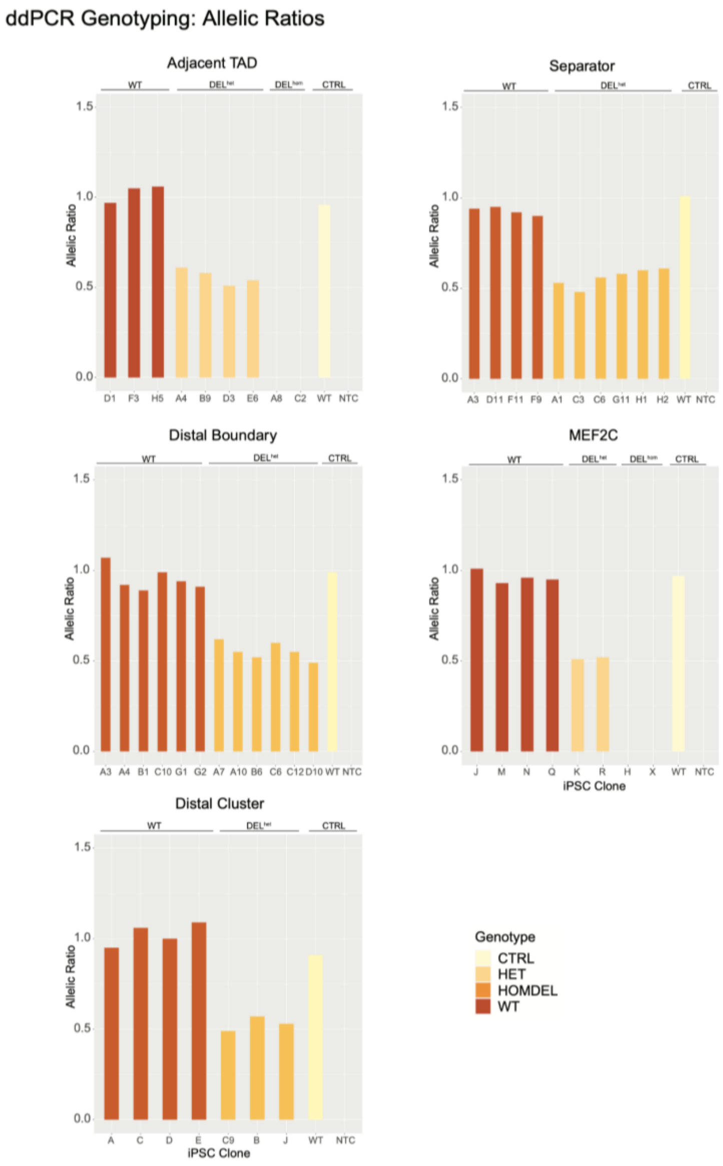

Supplementary Figure 12 : ddPCR Allelic Ratios per clone per target.
Using droplet counts generated by ddPCR, calculated relative allelic ratios of each target comparing counts for target site vs counts for RNaseP.

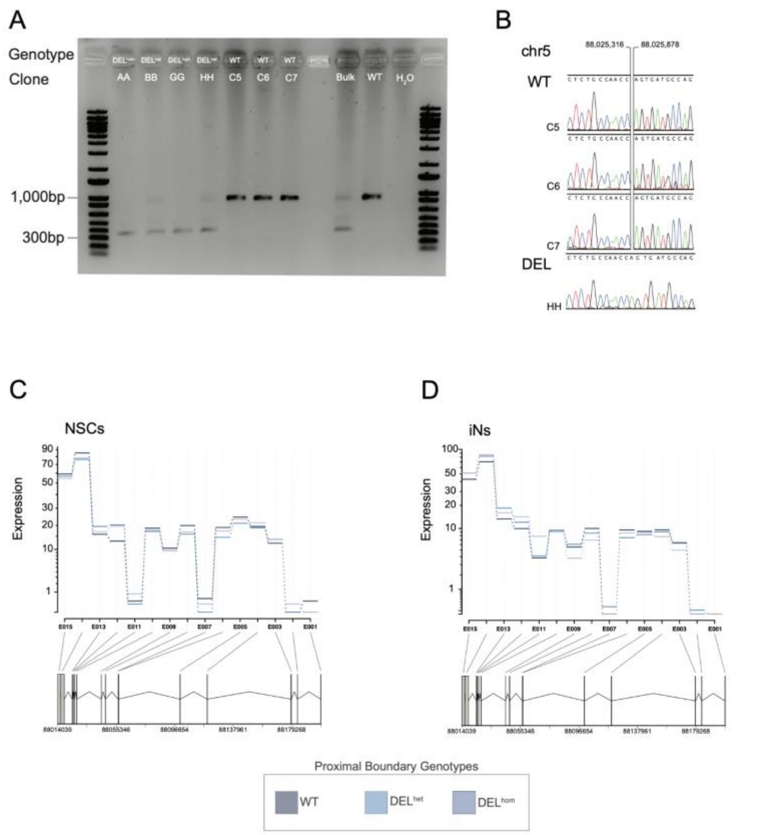

Supplementary Figure 13 : Proximal Boundary characterization.
**A.** PCR characterization of wildtype (906bp) and deletion (325bp) iPS clones, corresponding to the targeted 581bp deletion. Bulk edited iPSC DNA, DNA from un-transfected iPSCs, and water included as controls. **B.** Sanger traces of wildtype and gel extracted deletion band. (**C,D).** Exon usage of *MEF2C* from NSCs and iNs in **Proximal Boundary** DEL$^{het}$ and DEL$^{hom}$ vs WT samples.

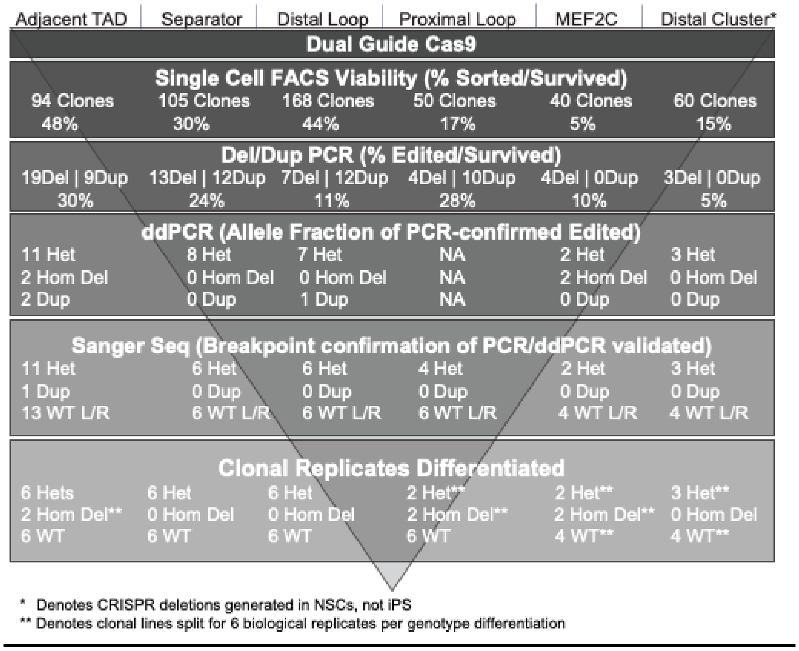

Supplementary Figure 14 : CRISPR target editing efficiencies by locus.

Breakdown of viabilities and editing efficiencies per target locus for each nested genotyping method used. Biological replicate iPSC lines representing independent editing events were retained for subsequent differentiation. For genotypes where 6 biological replicates were not obtained at the iPS stage, iPS lines were split prior to differentiation and technical replicates were differentiated for final retention of 6 differentiated lines per genotype per locus.

#### Section 3.4: Differentiation marker expression validation in positional effect iNs and NSCs

iN and NSC generation for positional effect CRISPR lines was confirmed using immunocytochemical staining of known protein markers of differentiation (**Supplementary Figure 15-21**).

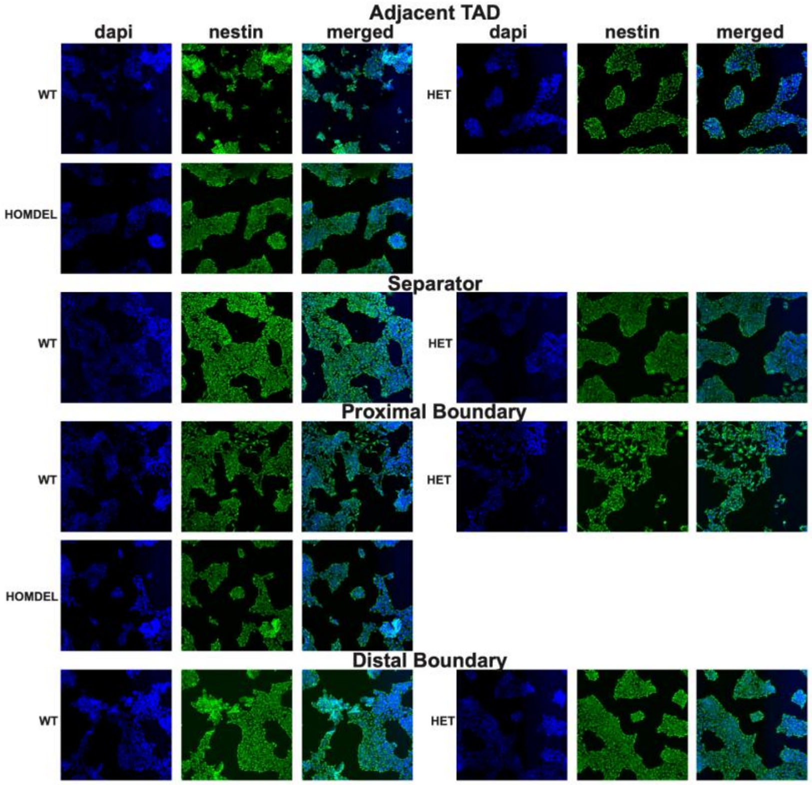

Supplementary Figure 15 : Nestin (NES) staining of positional effect CRISPR NSCs.

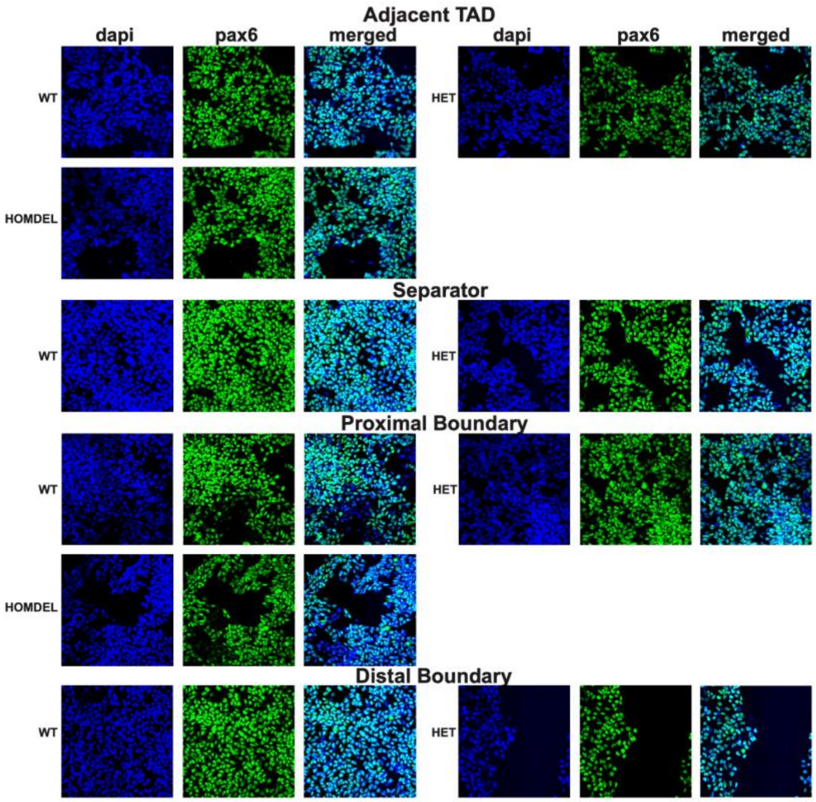

Supplementary Figure 16 : PAX6 staining of positional effect CRISPR NSCs.

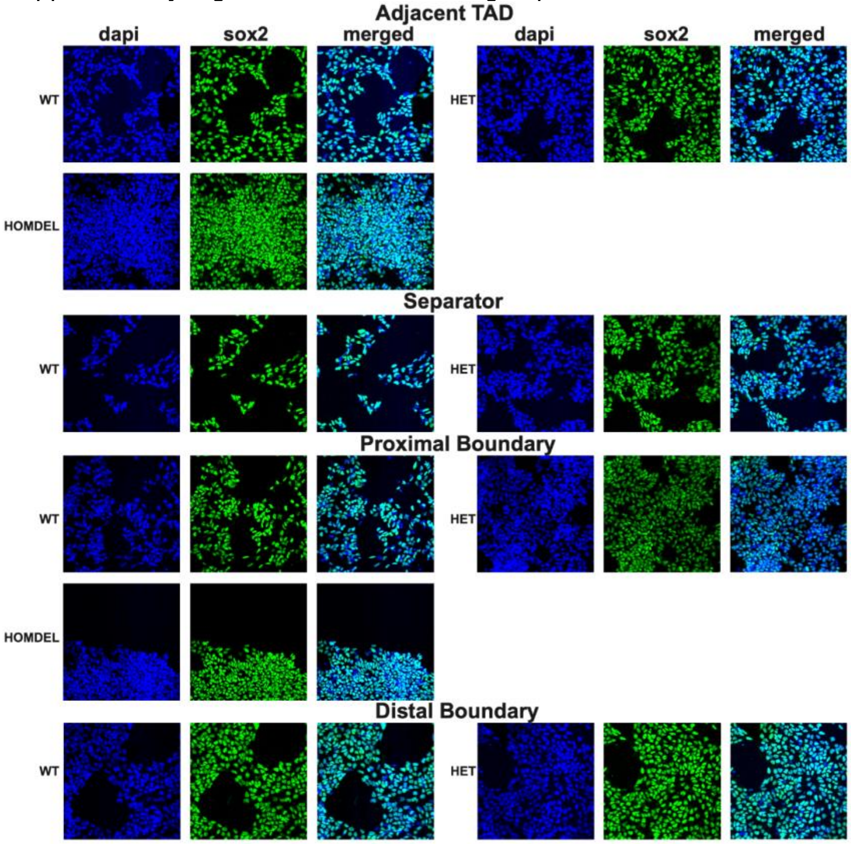

Supplementary Figure 17 : SOX2 staining of positional effect CRISPR NSCs.

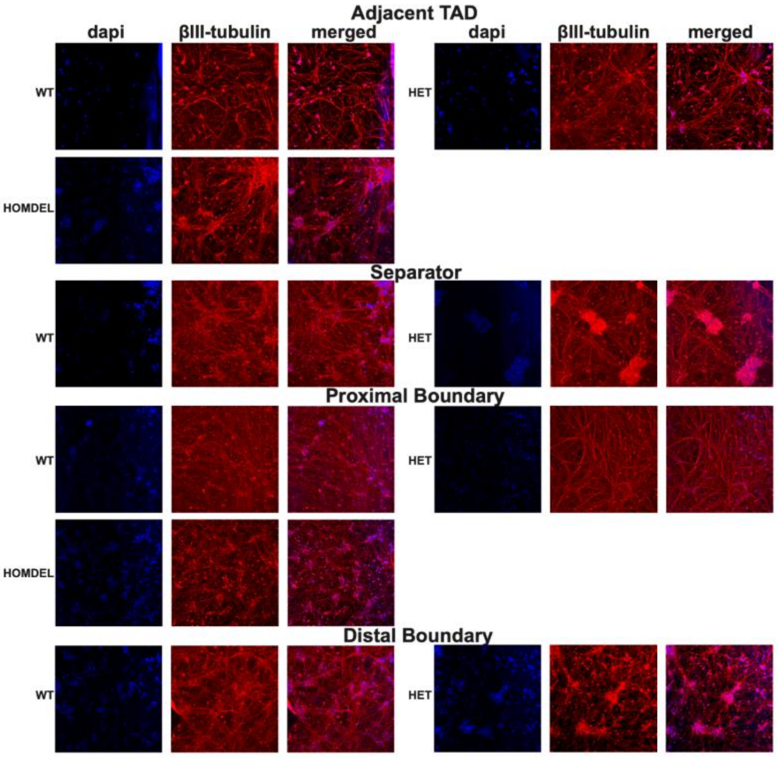

Supplementary Figure 18 : BetaIII-tubulin (TUBB3) staining of positional effect CRISPR iNs.

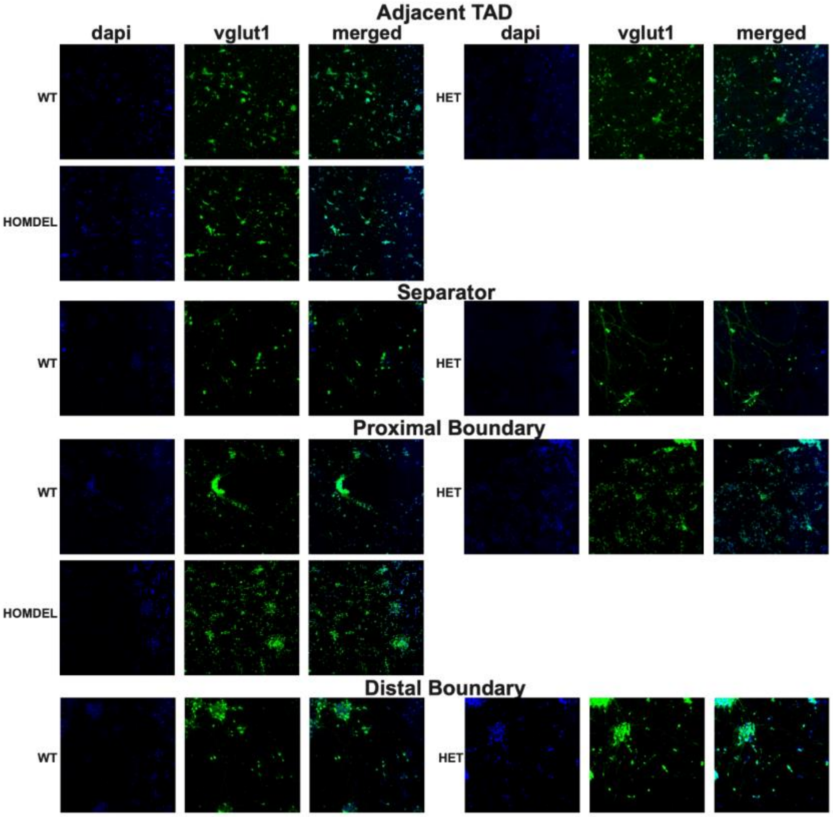

Supplementary Figure 19 : VGLUT1 staining of positional effect CRISPR iNs.

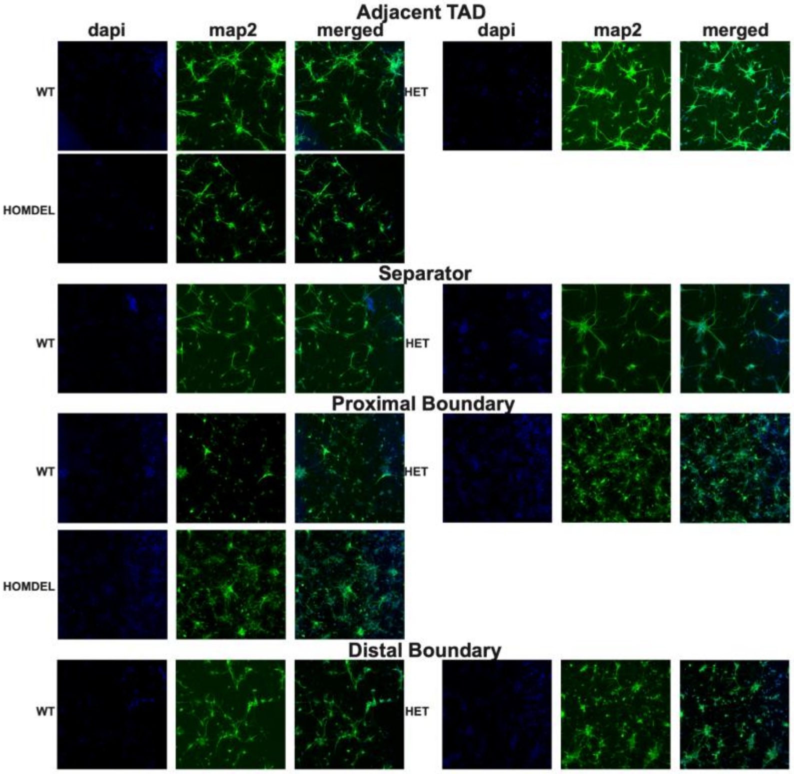

Supplementary Figure 20 : MAP2 staining of positional effect CRISPR iNs.

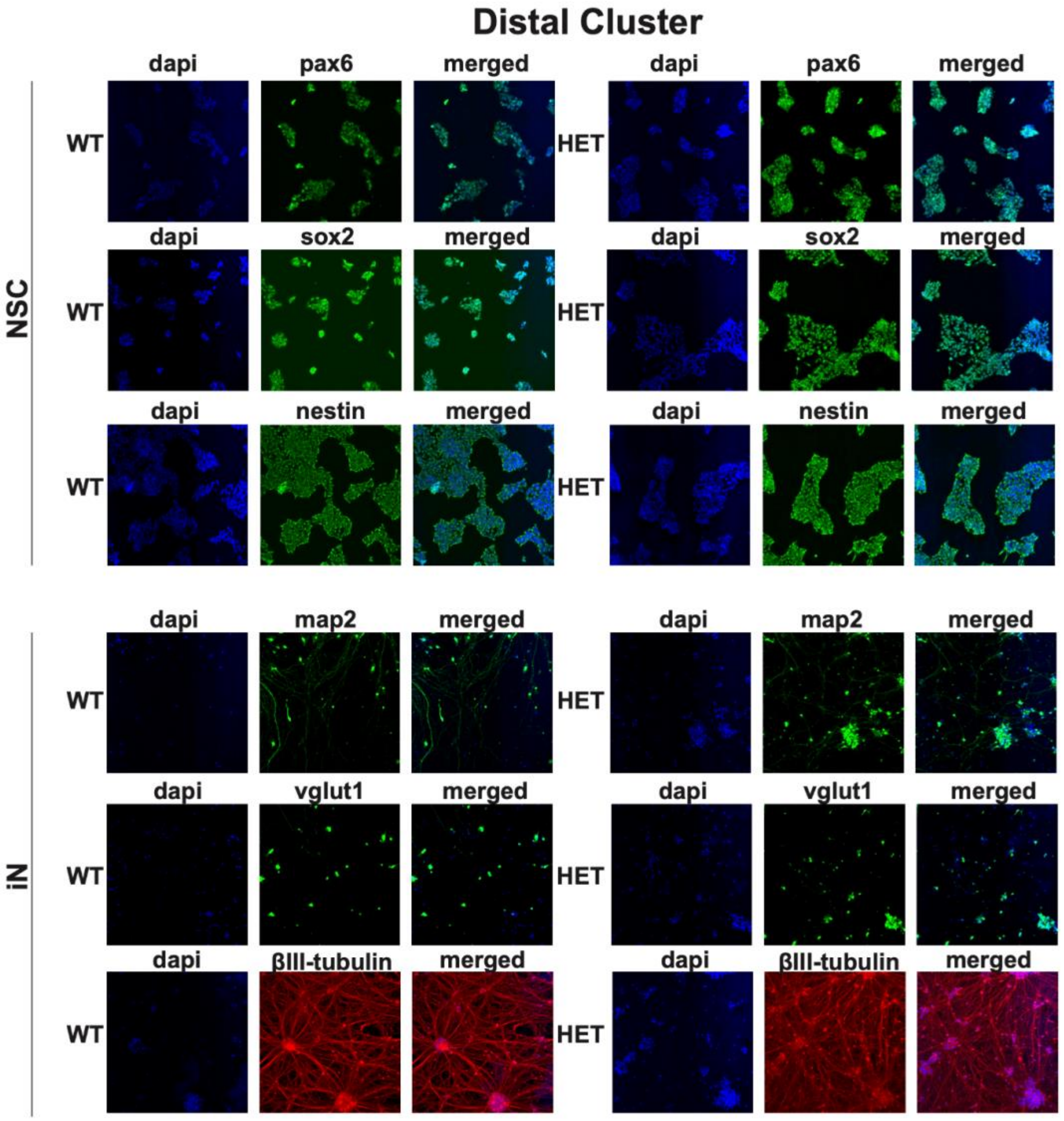

Supplementary Figure 21 : Staining of iN and NSC marker genes in Distal Cluster NSCs and NSC-derived iNs.

#### Section 3.5: MEF2C protein expression in positional effect CRISPR NSCs

We sought to follow-up on the observed *MEF2C* expression effects from RNAseq and consider MEF2C protein expression in NSCs using Western Blot for a subset of clonal lines from each CRISPR set. Using the same materials, methods, and analyses as the direct gene deletion lines, we calculated relative MEF2C protein expression per genotype per CRISPR set. We observe concordant differential expression results between RNA quantification from RNAseq and protein quantification from Western blot for most sets, as demonstrated in **Supplementary Figure 22**). The one exception is the Proximal Boundary NSCs, in which we observe a 40-60% reduction of MEF2C expression in $DEL^{hom}$ lines, in contrast to the WT levels of $DEL^{het}$ and $DEL^{hom}$ expression for these NSCs quantified by RNAseq.

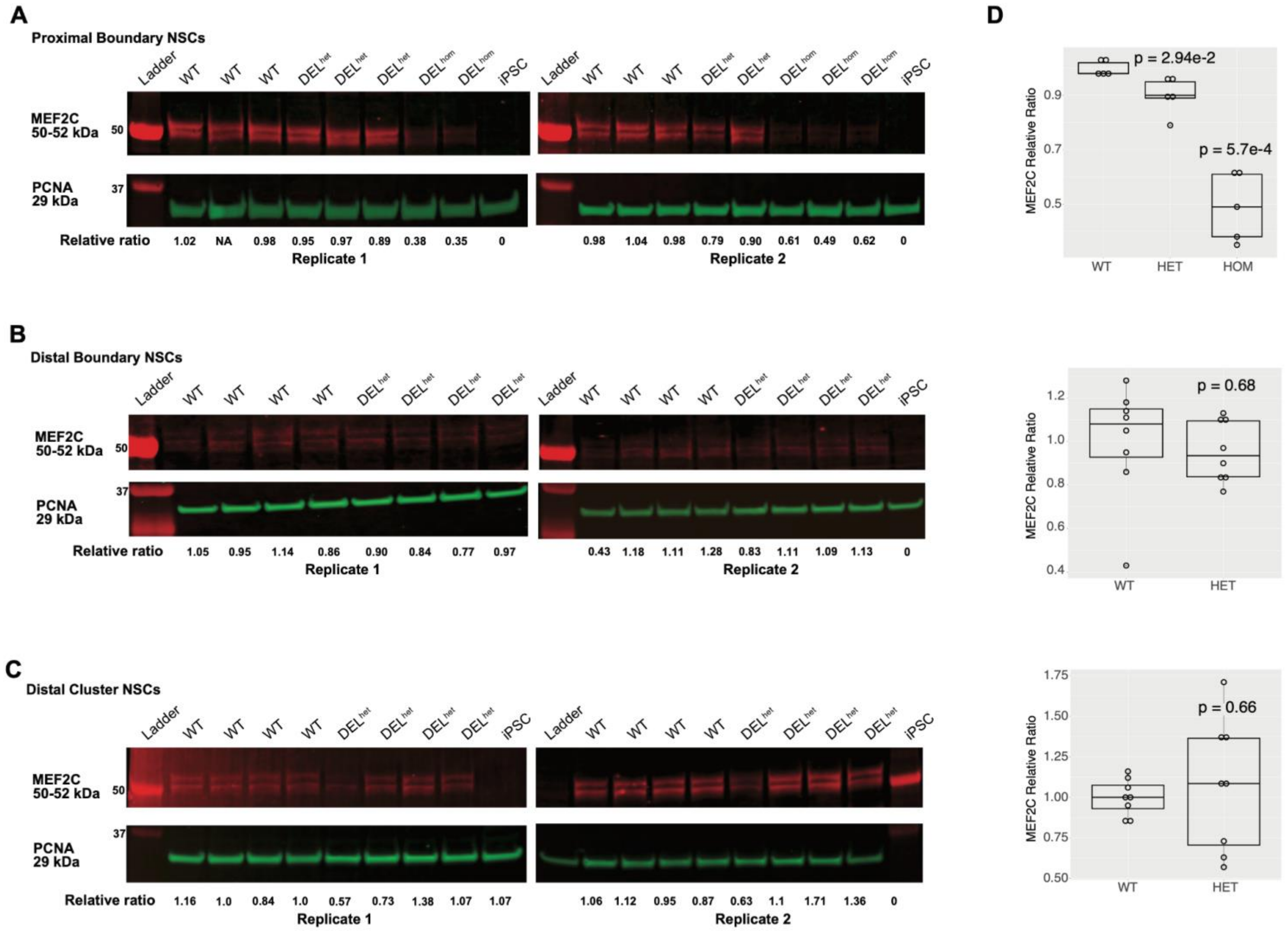
A
Proximal Boundary NSCs
Ladder
WT
DEL het
DEL hom
iPSC
MEF2C 50-52 kDa
50
PCNA 29 kDa
37
Relative ratio
1.02 NA 0.98 0.95 0.97 0.89 0.38 0.35 0
Replicate 1
0.98 1.04 0.98 0.79 0.90 0.61 0.49 0.62 0
Replicate 2
B
Distal Boundary NSCs
1.05 0.95 1.14 0.86 0.90 0.84 0.77 0.97
Replicate 1
0.43 1.18 1.11 1.28 0.83 1.11 1.09 1.13 0
Replicate 2
C
Distal Cluster NSCs
1.16 1.0 0.84 1.0 0.57 0.73 1.38 1.07 1.07
Replicate 1
1.06 1.12 0.95 0.87 0.63 1.1 1.71 1.36 0
Replicate 2
D
MEF2C Relative Ratio
p = 2.94e-2
p = 5.7e-4
WT
HET
HOM
p = 0.68
p = 0.66

Supplementary Figure 22 : MEF2C protein quantification for NSCs from 5q14.3 CRISPR lines.

**A.** MEF2C protein abundance quantified from NSCs with heterozygous and homozygous deletions to the Proximal Boundary relative to matched controls in two independent replicate blots. Clones represented in each blot include replicates of C8 (WT), C7 (WT), C6 (WT), BB ($DEL^{het}$), HH ($DEL^{het}$), AA ($DEL^{hom}$), and GG ($DEL^{hom}$). **B.** MEF2C protein abundance quantified from NSCs with heterozygous and homozygous deletions to the Distal Boundary relative to matched controls. Clones represented in each blot include replicates of A4 (WT), G1 (WT), C10 (WT), G1 (WT), A7 ($DEL^{het}$), and A10 ($DEL^{het}$). **C.** MEF2C protein abundance quantified from NSCs with heterozygous and homozygous deletions to the Distal Cluster relative to matched controls. Clones represented in each blot include replicates of A (WT), C9 ($DEL^{het}$), J ($DEL^{het}$), and B ($DEL^{het}$). iPSC lysate included as negative control given lack of MEF2C expression. **D.** Boxplots representing relative MEF2C protein expression per CRISPR group. P-values calculated using t-test relative to wildtype mean. Total MEF2C (50/51kDa + 52kDa bands) was used in MEF2C quantification relative to PCNA (29kDa band) for all.

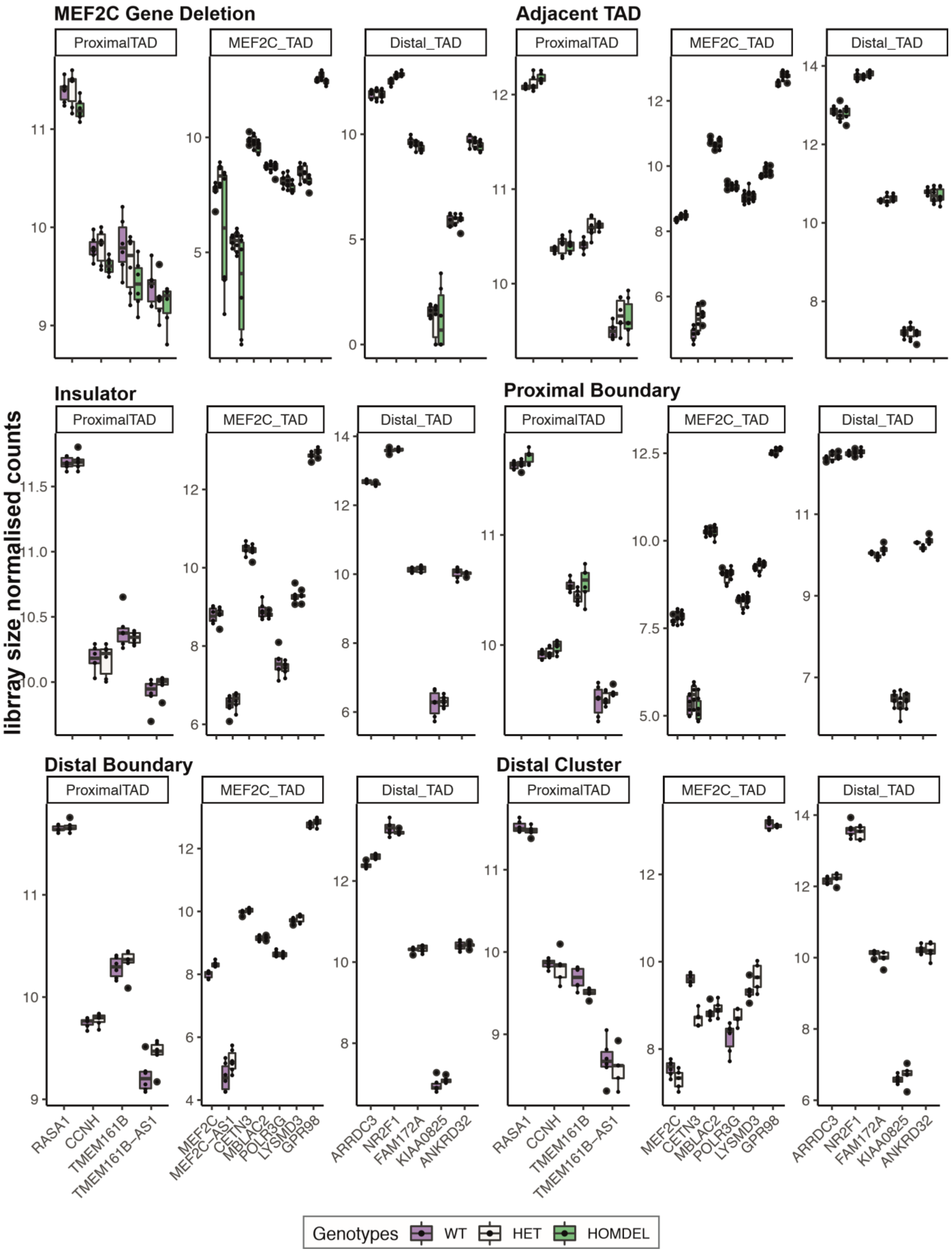

Supplementary Figure 23 : CRISPR NSCs expression boxplots

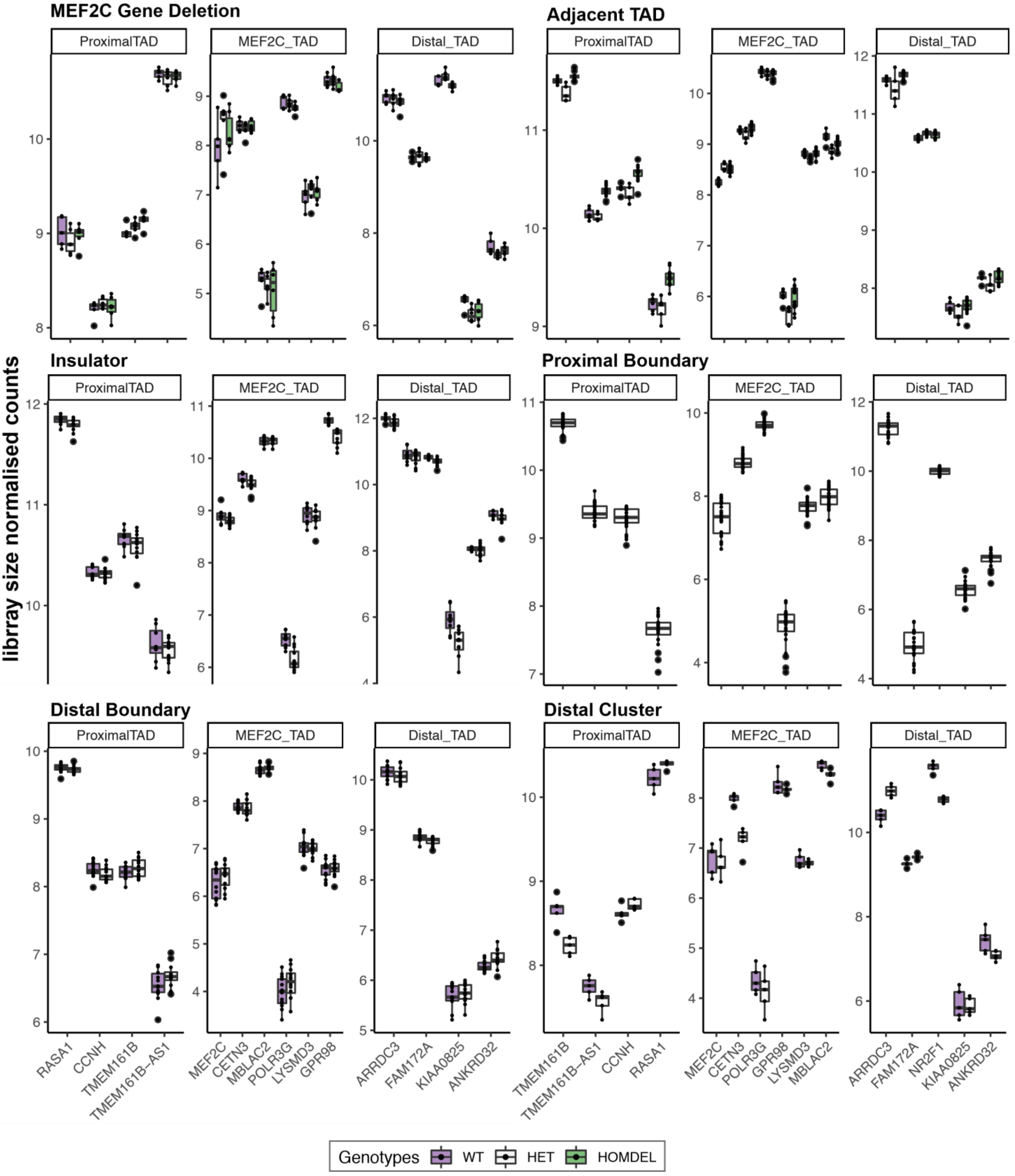

Supplementary Figure 24 : CRISPR iNs expression boxplots

#### Section 3.6: mRNA expression of local genes within 5q14.3 in CRISPR iNs and NSCs

In addition to our primary interest of determining whether 3D functional element disruption was sufficient to dysregulate *MEF2C* expression, we were also interested in surveying local gene expression effects within 5q14.3. Using SVA-normalized counts from RNAseq, we first considered mean and variance of expression per genotype per CRISPR differentiated set of 16 total genes distributed across the 5q14.3 locus. This included genes within the *MEF2C* containing TAD and both TADs immediately up and downstream of this 3D domain. For most CRISPR sets, we observe minor, if any, gene expression changes as a result of each deletion relative to matched wildtype, with most deviations from matched wildtype amounting to 15-20% gene expression change. This is true both for *MEF2C* specifically as well as other local genes in the locus. The one exception is the Proximal Boundary CRISPR differentiated to iNs. In this CRISPR set we observe a 45% (FDR <0.1) down-regulation of *MEF2C* in homozygous deletions of the Proximal Boundary alongside a genotype-specific expression decay for genes up to 1Mb distal to the perturbation site (**Supplementary Figure 23,24).** All local and global differentially expressed genes identified from RNAseq are submitted to GEO as processed data.

#### Section 3.7: Identification of distal contacts with *MEF2C* promoter using UMI-4C

We report aggregate read coverage per per individual CRISPR set (**Supplementary Figure 28-: 4Cseq coverage maps from viewpoint of *MEF2C* promoter in wildtype and CRISPR cell lines**). Given technical variables that allow for higher cellular input per sample for NSCs (5 million cells targeted) vs iNs (1 million cells targeted), we observe higher overall read coverage across the 5q14.3 locus in NSCs vs iNs. Allele-indiscriminate maps were constructed using baits against VP1 within the MEF2C promoter region whereas baits against VP2 allowed construction of allele-specific contact maps.

Data tracks that contributed to the creation of our 4Cseq figures are interactively viewable on the UCSC genome browser:
http://genome.ucsc.edu/s/kianamohajeri/5q14.3_CRISPR_4C_Coverage
http://genome.ucsc.edu/s/kianamohajeri/5q14.3_CRISPR_4C_LogFC

See these UCSC sessions for the processed data (normalized and smoothed, per sample and pooled, + subtraction profiles):

4C_MEF2C_final_WT
4C_MEF2C_final_proxTAD (adjacent TAD)
4C_MEF2C_final_insul
4C_MEF2C_final_proxLoop
4C_MEF2C_final_mef2c
4C_MEF2C_final_distLoop
4C_MEF2C_final_distLoopLRG (distal cluster)

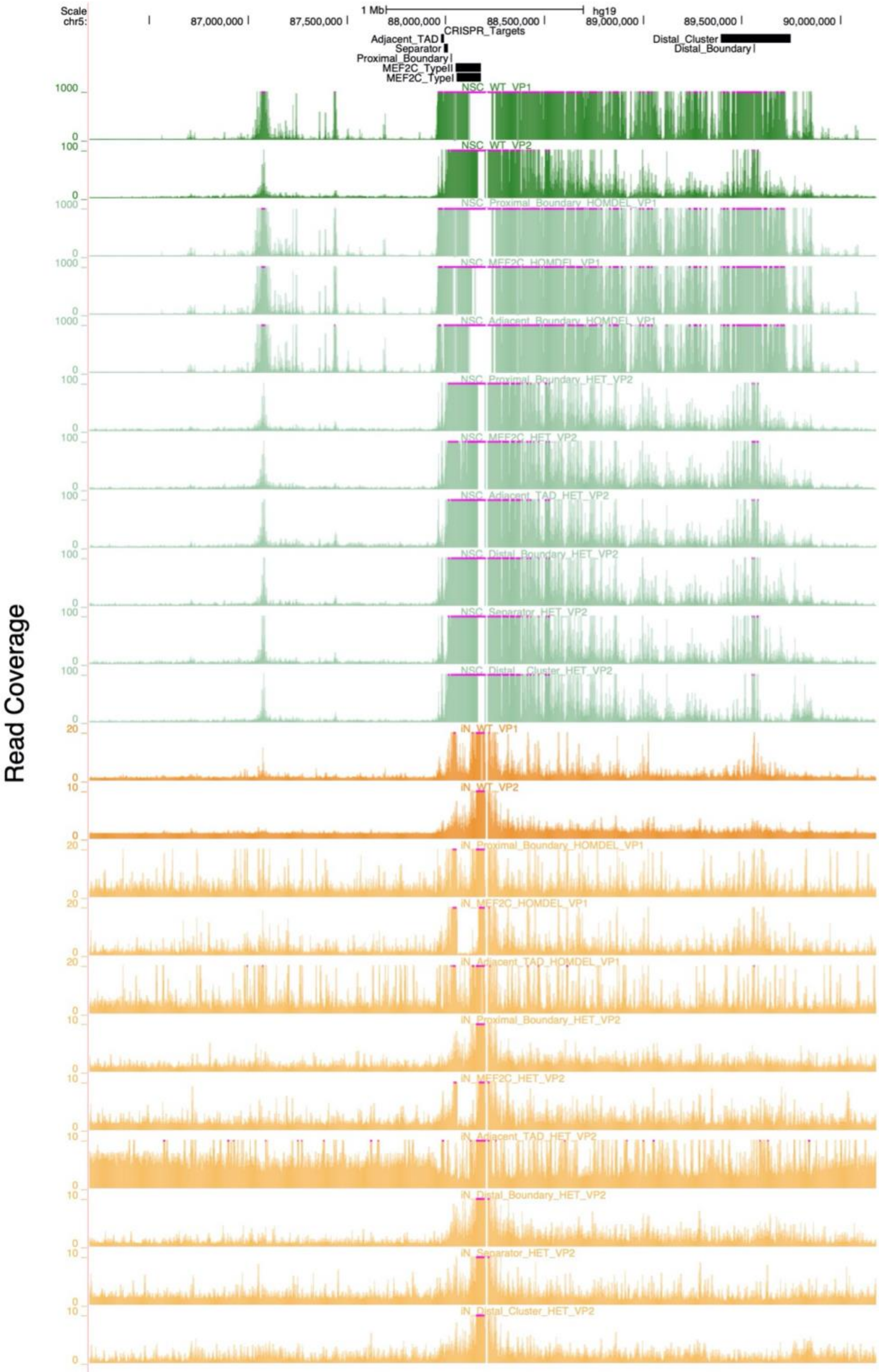

Supplementary Figure 25 : 4C coverage maps from viewpoint of MEF2C promoter in wildtype and CRISPR lines.

Clone replicates per genotype per locus displayed in aggregate on single track. Green tracks represent NSCs with iNs in orange.

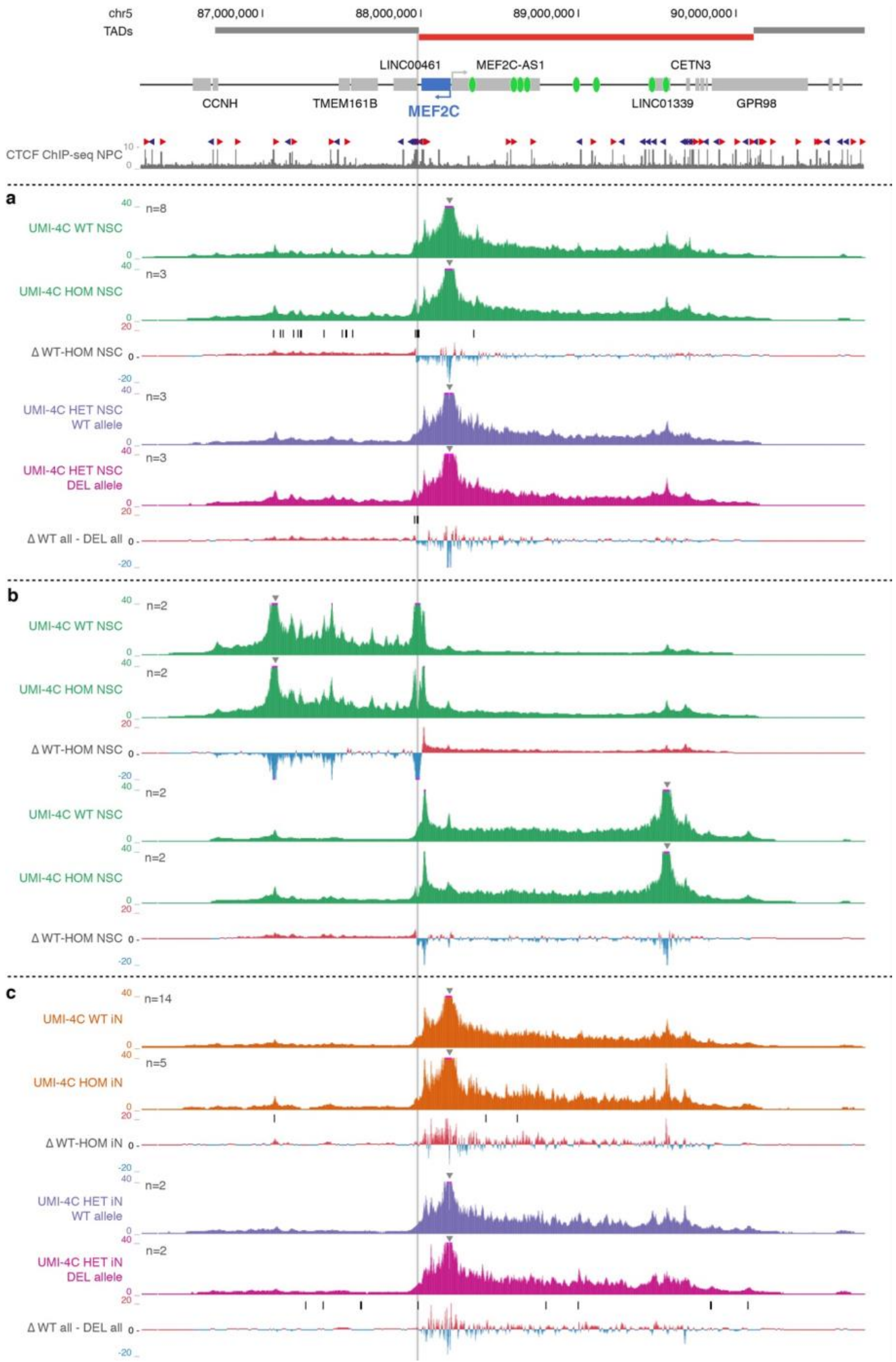

Supplementary Figure 26 : Chromatin interactions using UMI-4C in Adjacent TAD NSCs and iNs.

Normalized and smoothed UMI-4C interaction counts and subtraction (Δ) profiles. UMI-4C viewpoints (VPs) are indicated by arrows, the deletion by a grey bar, significant differential interactions (FDR 0.1) by black boxes. N represents the number of biological replicates. MEF2C locus with topologically associating domain (TAD) structure, annotated enhancer elements (green ovals) and CTCF binding in neural progenitor cells (NPCs, available from ENCODE) on top. (a) Wildtype (WT) neural stem cells (NSCs) vs NSCs with homozygous (HOM) deletion of the Adjacent TAD (MEF2C VP1). Heterozygous (HET) deletion, WT vs deletion (DEL) allele (MEF2C VP2). (b) WT vs HOM NSCs from two alternative VPs. (c) WT vs HOM induced neurons (iNs) (MEF2C VP1). WT vs DEL allele of HET iNs (MEF2C VP2).

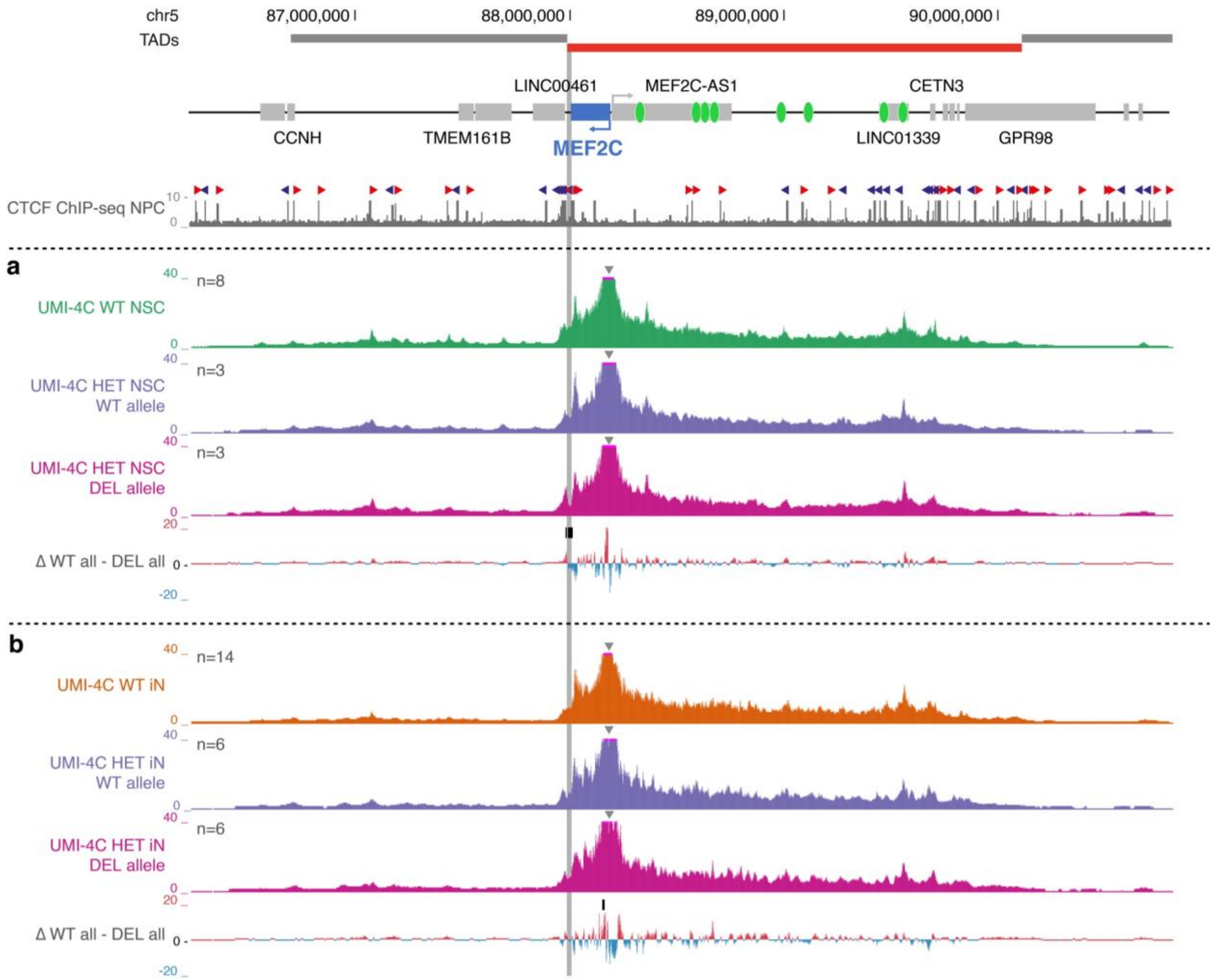

Supplementary Figure 27 : Chromatin interactions using UMI-4C in Separator NSCs and iNs.

Normalized and smoothed UMI-4C interaction counts and subtraction (Δ) profiles. UMI-4C viewpoints (VPs) are indicated by arrows, the deletion by a grey bar, significant differential interactions (FDR 0.1) by black boxes. N represents the number of biological replicates. MEF2C locus with topologically associating domain (TAD) structure, annotated enhancer elements (green ovals) and CTCF binding in neural progenitor cells (NPCs, available from ENCODE) on top. (a) Heterozygous (HET) deletion of Separator region in neural stem cells (NSCs), wildtype (WT) vs deletion (DEL) allele (MEF2C VP2). (b) HET deletion of Separator region in induced neurons (iNs), WT vs DEL allele (MEF2C VP2).

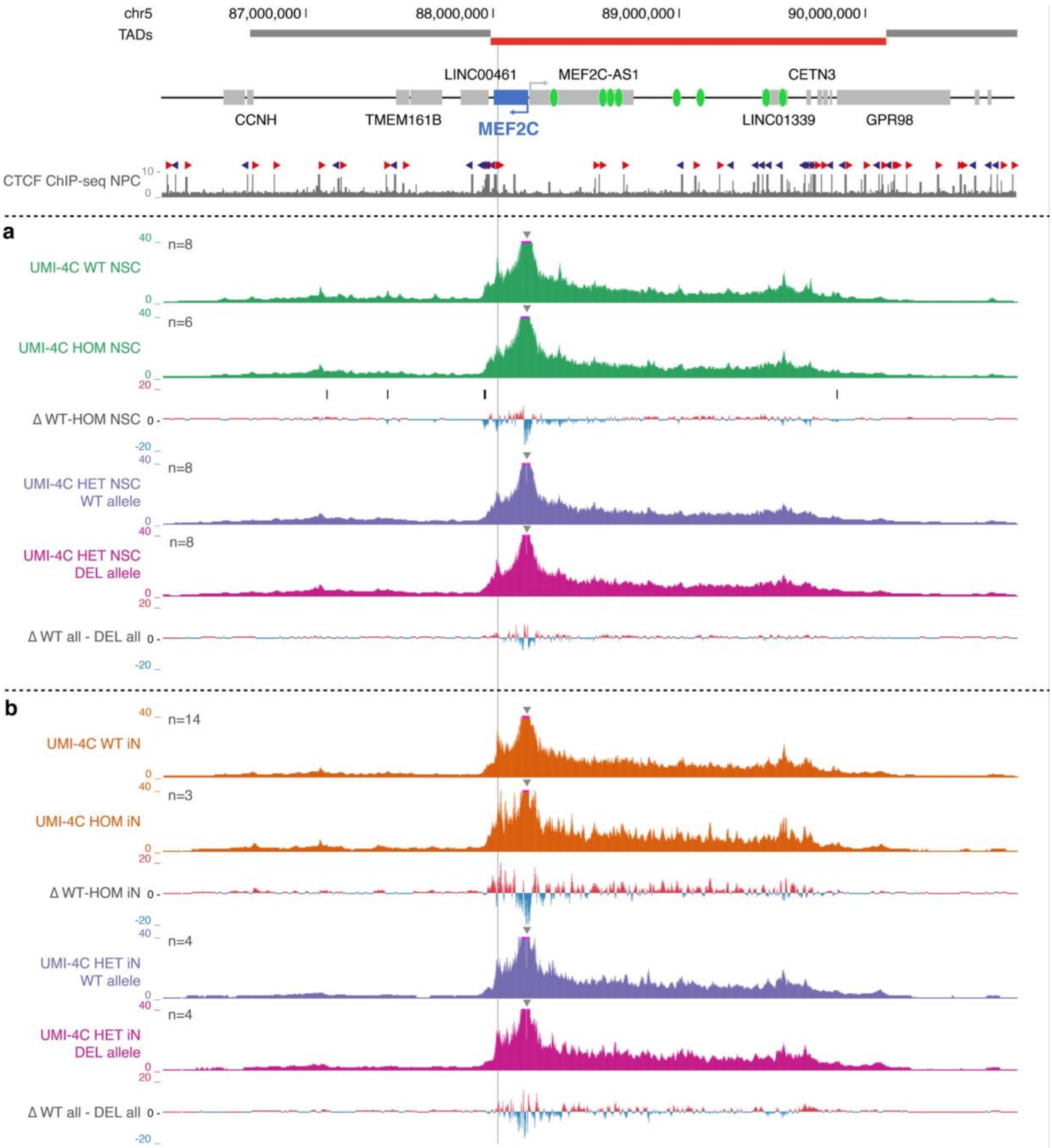

Supplementary Figure 28 : Chromatin interactions using UMI-4C in Proximal BoundaryLoop NSCs and iNs.

Normalized and smoothed UMI-4C interaction counts and subtraction (Δ) profiles. UMI-4C viewpoints (VPs) are indicated by arrows, the deletion by a grey bar, significant differential interactions (FDR 0.1) by black boxes. N represents the number of biological replicates. MEF2C locus with topologically associating domain (TAD) structure, annotated enhancer elements (green ovals) and CTCF binding in neural progenitor cells (NPCs, available from ENCODE) on top. (a) Wildtype (WT) neural stem cells (NSCs) vs NSCs with homozygous (HOM) deletion of the Proximal Loop (MEF2C VP1). Heterozygous (HET) deletion, WT vs deletion (DEL) allele (MEF2C VP2). (b) WT vs HOM induced neurons (iNs) (MEF2C VP1). WT vs DEL allele of HET iNs (MEF2C VP2).

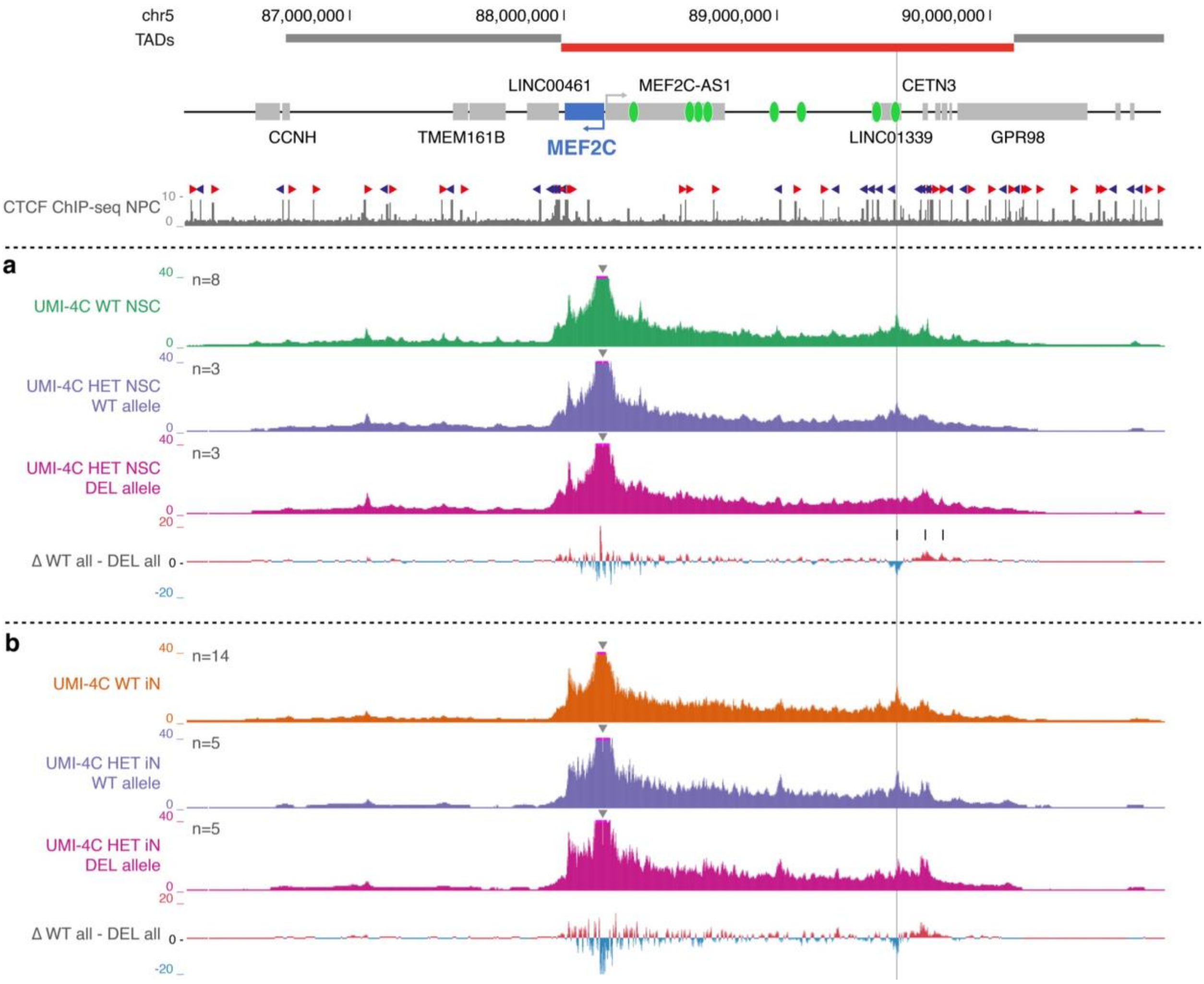

Supplementary Figure 29 : Chromatin interactions using UMI-4C in Distal BoundaryLoop NSCs and iNs.

Normalized and smoothed UMI-4C interaction counts and subtraction (Δ) profiles. UMI-4C viewpoints (VPs) are indicated by arrows, the deletion by a grey bar, significant differential interactions (FDR 0.1) by black boxes. N represents the number of biological replicates. MEF2C locus with topologically associating domain (TAD) structure, annotated enhancer elements (green ovals) and CTCF binding in neural progenitor cells (NPCs, available from ENCODE) on top. (a) Heterozygous (HET) deletion of the Distal Loop in neural stem cells (NSCs), wildtype (WT) vs deletion (DEL) allele (MEF2C VP2). (b) HET deletion of the Distal Loop in induced neurons (iNs), WT vs DEL allele (MEF2C VP2).

#### Section 3.8: Proximal Boundary CRISPR multi-electrode array (MEA) assessment of synaptic activity

We conducted MEA on the Proximal Boundary CRISPR iNs and matched controls using methods outlined in **Materials and Methods**, following the same protocols used for MEA on direct disruption *MEF2C* iNs. Two 48-well MEA plates were included in analyses reflected in **Supplementary Figures 30-31**, referred to as Proximal Boundary MEA Plate 1 and Proximal Boundary MEA Plate 2. The 48 total wells on Plate 1 contained 16 wells of wildtype cells (8 wells clone C5, 10 wells clone C6), 12 wells of heterozygous cells (6 wells clone BB, 6 wells clone HH), 12 wells of homozygous cells (8 wells clone AA, 4 wells clone GG), and 8 media-only wells. The 48 total wells on Plate 2 contained 16 wells of wildtype cells (8 wells clone C5, 8 wells clone C7), 12 wells of heterozygous cells (6 wells clone BB, 6 wells clone HH), 12 wells of homozygous cells (8 wells clone AA, 4 wells clone GG), and 8 media-only wells. Replicate wells of clone C5 were included on both Plates 1 and 2 as a wildtype comparator set.

Considering wells from both Plates 1 and 2 we plated a total of 32 replicate wildtype wells, 24 replicate heterozygous deletion wells, and 24 replicate homozygous deletion wells. Well filtering was conducted to retain wells with at least 50% active electrodes (requiring 8 of 16 total electrodes per well passing activity threshold, as defined by manufacturer) for downstream analyses (**Supplementary Figures 30-31**). Clones C6 (WT) and BB (DEL$^{het}$) were removed from downstream analyses given lower than threshold electrode activity through half of differentiation.

Combined Plate 1 and Plate 2 analysis was conducted by first normalizing all data points by wildtype average for a given measure on each day per each respective plate. P-values calculated using paired t-test comparing mean value of each measurement vs wildtype (**Supplementary Figure 32**)

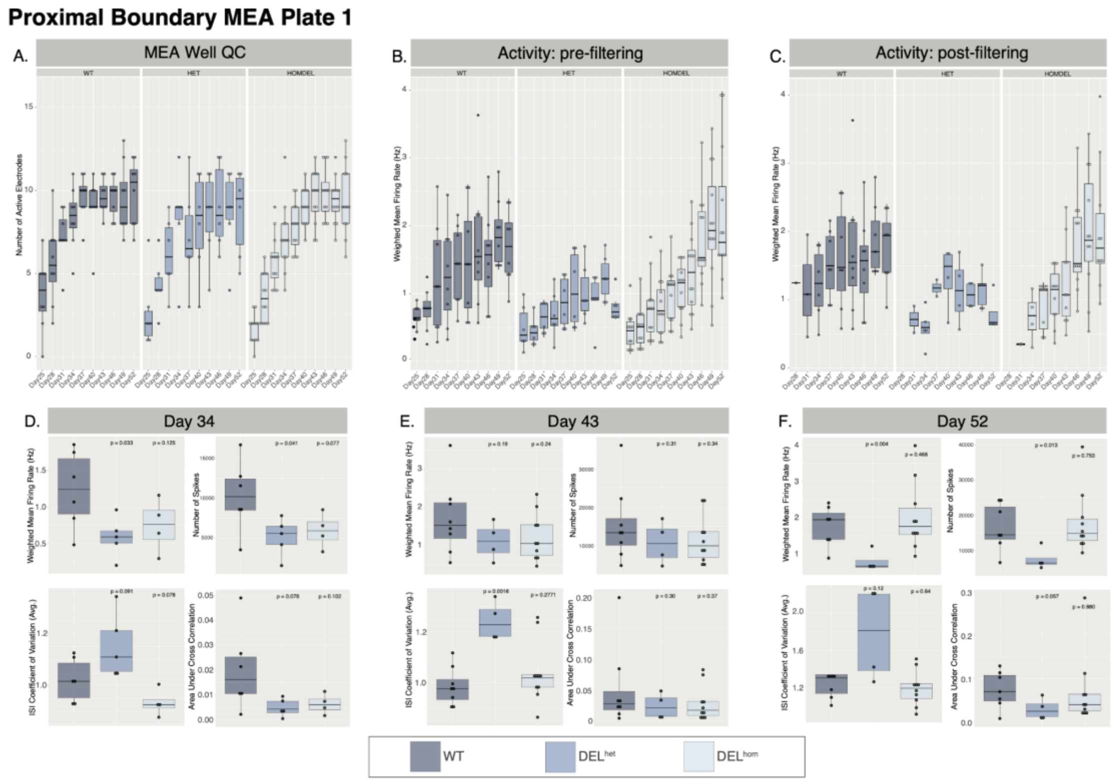

Supplementary Figure 30 : MEA on iNs with Proximal Boundary deletion (Plate 1). Well QC demonstrating number of active electrodes per genotype per MEA reading day. Weighted mean firing rate per genotype per MEA reading day shown before **B.** and after **C.** filtering for wells with at least 8 active electrodes. **D-G.** measurements per genotype on early (Day 34), mid (Day 43), and late (Day 52) differentiation timepoints.

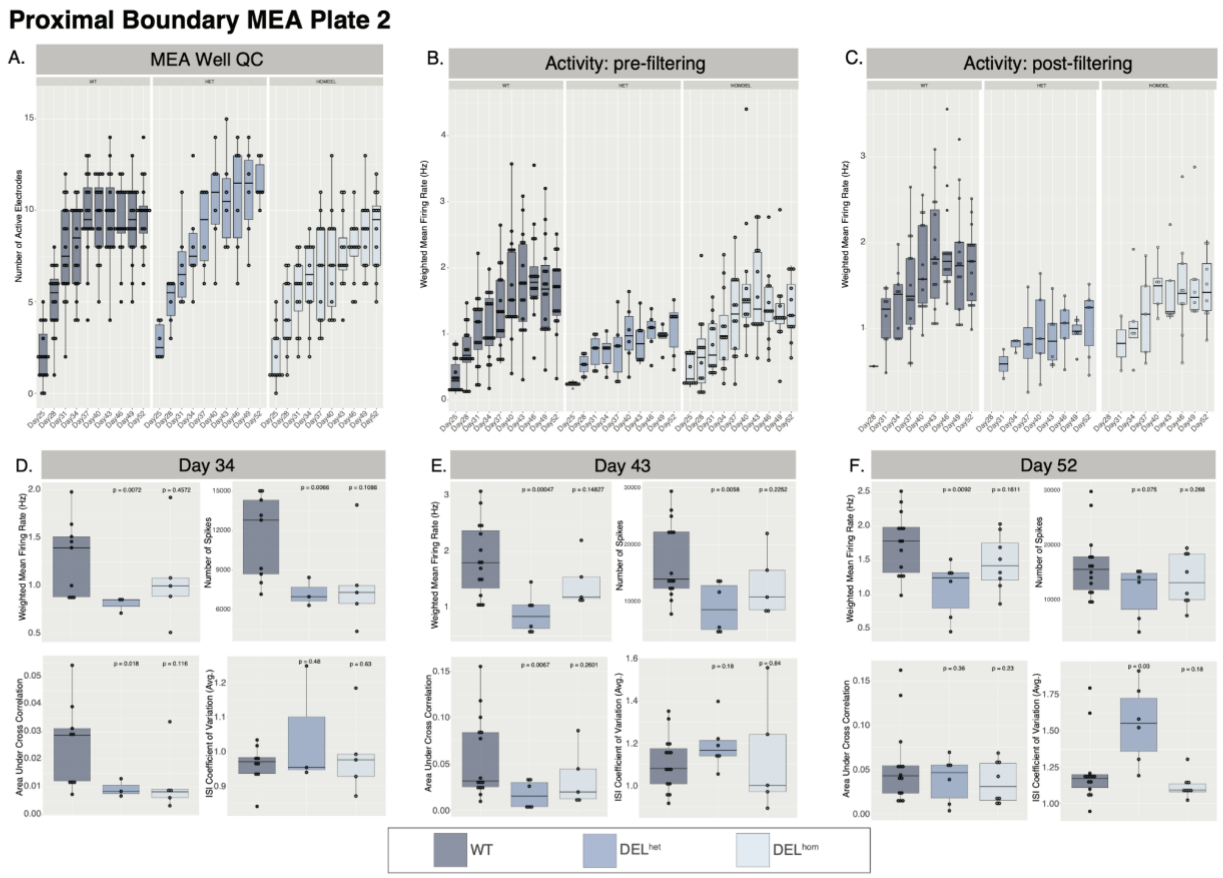

Supplementary Figure 31 : MEA on iNs with Proximal Boundary deletion (Plate 2). Well QC demonstrating number of active electrodes per genotype per MEA reading day. Weighted mean firing rate per genotype per MEA reading day shown before **B.** and after **C.** filtering for wells with at least 8 active electrodes. **D-G.** measurements per genotype on early (Day 34), mid (Day 43), and late (Day 52) differentiation timepoints.

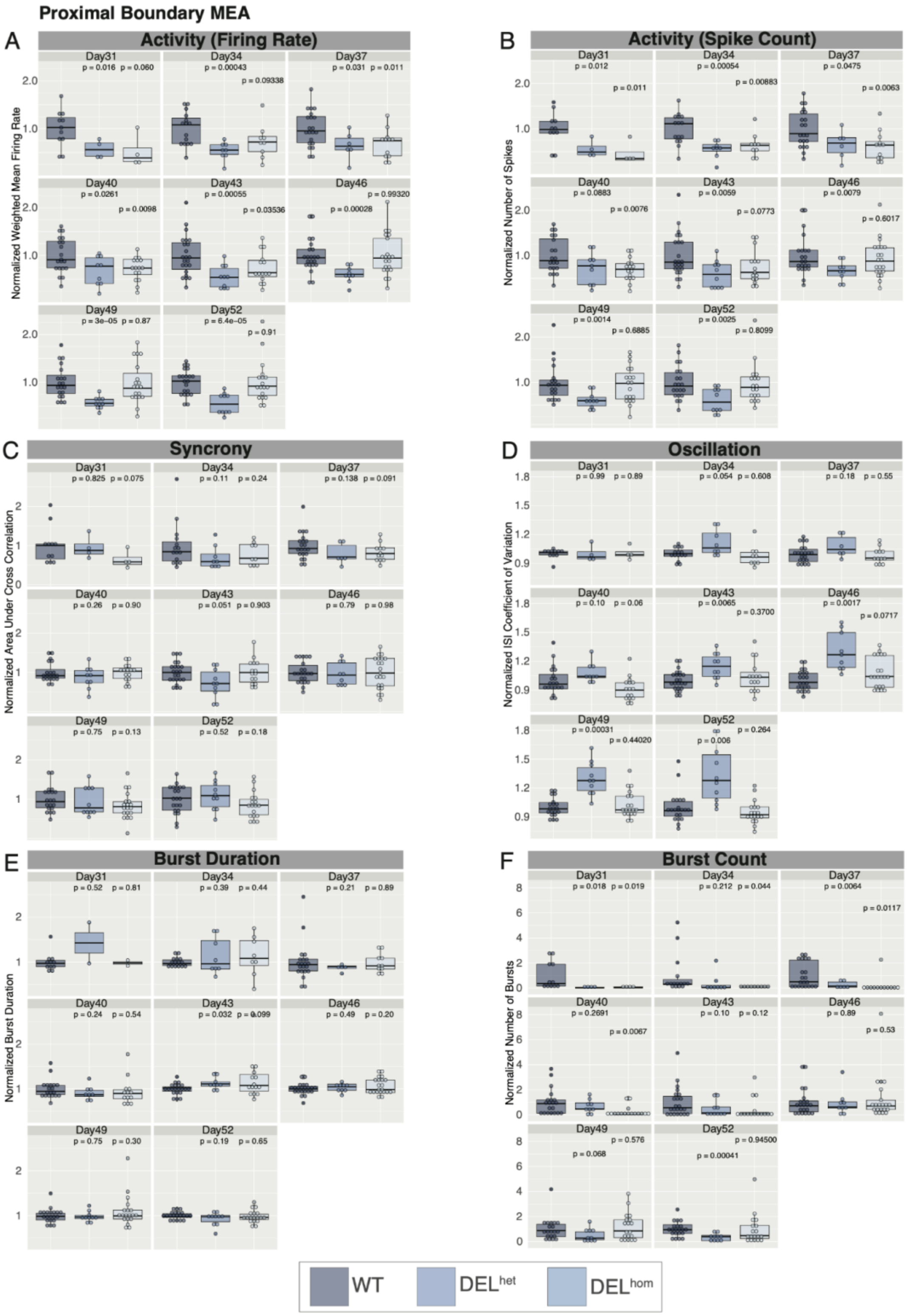

Supplementary Figure 32 : Combined time course on iNs with Proximal Boundary Deletion.

Datapoints normalized relative to respective wildtype measurement average per plate and plotted collectively. Weighted mean firing rate (**A**), spike count (**B**), synchrony (**C**), Oscillation (**D**), burst duration (**E**), and burst count (**F**) shown per day per genotype. P-values calculated using t-test against wildtype mean.

### Supplementary Tables

(Supplied as excel sheet)
Supplementary Table 1: CRISPR/Cas9 guide sequences used in this study
Supplementary Table 2: ddPCR probe and PCR primer sequences used for genotyping in this study
Supplementary Table 3: MEF2C protein quantification calculations from Western blot
Supplementary Table 43: Genotyping results per clone from ddPCR and Sanger sequencing
Supplementary Table 5: Overview of iN and NSC differentiation batches included and functional followup
Supplementary Table 6: Sample list of iNs and NSCs for RNAseq
Supplementary Table 7: Sample list of iNs and NSCs for 4Cseq
